# The wheat *Seven in Absentia* gene is associated with increases in biomass and yield in hot climates

**DOI:** 10.1101/726802

**Authors:** Pauline Thomelin, Julien Bonneau, Chris Brien, Radoslaw Suchecki, Ute Baumann, Priyanka Kalambettu, Peter Langridge, Penny Tricker, Delphine Fleury

**Author notes:** Corresponding author: Penny Tricker, Plant Genomics Centre, Hartley Grove, Urrbrae, South Australia, 5064.

## Abstract

Wheat productivity is severely reduced by high temperatures. Breeding of heat tolerant cultivars can be achieved by identifying genes controlling physiological and agronomical traits with high temperature and using these to select superior genotypes, but no gene underlying genetic variation for heat tolerance has previously been described. We completed the positional cloning of *qYDH.3BL*, a quantitative trait locus (QTL) on bread wheat chromosome 3B associated with increased yield in hot and dry climates. The delimited genomic region contained 12 putative genes and a sequence variant in the promoter region of one gene - *seven in absentia, TaSINA*. This was associated with the QTL’s effects on early vigour, plant biomass and yield components in two distinct wheat populations grown under various growth conditions. Near isogenic lines carrying the positive allele at *qYDH.3BL* under-expressed *TaSINA* and had increased vigour and water use efficiency early in development, as well as increased biomass, grain number and grain weight following heat stress. A survey of worldwide distribution indicated that the positive allele became widespread from the 1950s through the CIMMYT wheat breeding programme but, to date, has been selected only in breeding programmes in Mexico and Australia.

**Significance statement:** Wheat is the world’s most widely grown crop and a staple of human diet. Even brief episodes of high temperature in the growing season cause severe yield reductions. Finding and deploying genes for heat stress tolerance in new varieties is a priority for food security with climate change. We narrowed a genetic locus to a small genomic region where genetic variation was present only in one gene that showed clear differences of expression and improved yield and physiology under stress in the populations. Using diagnostic markers to track the positive haplotype in nearly 750 accessions, we found many regions where the allele could be used in breeding programmes to increase wheat’s heat tolerance.

## Introduction

As one of the world’s major crops providing 20 % of human food, bread wheat (*Triticum aestivum* L.) has a vital role in food security^1^. It is also the most widely produced crop, grown worldwide in a variety of climates including harsh environments. Wheat productivity is affected by growing season drought and heat stresses, often in combination, which can cause almost complete yield loss in some cases^2,3^. Due to the increasing occurrence of drought and heat stress, particularly in the Mediterranean region, the USA, India and Australia, which are amongst the largest producers of bread wheat in the world, improving yield under these stresses is a priority^4,5^. Targeting crop productivity in regions affected by drought and heat is believed to be the best strategy to reach the 1.6 % yield improvement per year required to meet the food needs of an increasing world population^6^.

One way to deliver new, high yielding varieties is to discover and deploy genes associated with grain yield variation in stress-prone environments through breeding. Many genetic studies have focussed on the identification of quantitative trait loci (QTL) associated with yield variation under abiotic stress and, in wheat, several genes have been identified for tolerance of salinity, cold, aluminium and boron toxicity^7–10^. Genetic variation in wheat yields in dry and hot conditions has been extensively studied using bi-parental populations^11–27^ but no genes underlying QTL for heat and drought tolerance have yet been successfully identified. Grain yield is a complex trait, highly influenced by the environment and, although QTL for yield have been identified, only a few have been used in breeding programmes due to inconsistent performance of the QTL in different environmental conditions.

We focussed on a QTL located on chromosome 3BL, *qYDH.3BL*, identified in a doubled-haploid (DH) population from the cross between the Australian wheats RAC875 and Kukri. These lines were selected for their contrasting physiological responses in Mediterranean-like climates: RAC875 is a water conservative line, whereas Kukri depletes water more quickly under stress^28^. In the RAC875 x Kukri DH population, a multi-environment analysis of 21 field trials showed a strong genotype by environment (G x E) interaction at the locus^29^. The QTL was expressed in the deep soil of northern Mexico with the RAC875 allele positively contributing to yield, thousand grain weight and early vigour. The Kukri allele at *qYDH.3BL* was associated with increased yield in southern Australian trials, but only when plants were irrigated^11,29^. Parent et al.^30^ subsequently observed that the allele effect at *qYDH.3BL* was dependent on growth temperatures. In this study, we fine mapped the QTL to a short region of 690 Kbp sequence which includes 12 putative genes. Furthermore, we identified sequence variants and differential expression of a *SINA* gene consistent with the positive effects of one allele on early growth in unstressed conditions and plant biomass following heat stress. We identified and tracked the allele in a worldwide collection of wheat accessions to enable its future use.

## Results

The QTL *qYDH.3BL*, originally found in the spring wheat RAC875 x Kukri DH population^29^ was also present in a Drysdale x Gladius recombinant inbred line (RIL) population^31^. The analysis of these two populations across multiple trials in Australia and Mexico showed that at this locus the allele from RAC875 and Drysdale increased yield relative to the Kukri and Gladius allele (Fig. 1a, b). The multi-environment analysis of both populations showed that the high confidence QTL interval was flanked by the markers AWG43_1 and AWG38 (Fig. 1a, b). The RAC875 allele was positively associated with a 10 % yield increase (Fig. 1b). We fine mapped *qYDH.3BL* in RAC875 by anchoring the interval defined by the multi-environment analysis onto the v1.0 reference sequence of cv. Chinese Spring (IWGSC RefSeq v1.0)^32^. We then aligned the whole genome shotgun sequencing data of the four parental lines^33^ against the reference genome to identify sequence polymorphisms. Comparison between the genotypes RAC875-Drysdale vs Kukri-Gladius enabled us to identify new sequence variants. Seventy-four single nucleotide polymorphism (SNP)-based markers (named AWG and ADW hereafter) and InDel markers were added to the previously described RAC875 x Kukri RIL map^29^. The interval delimited by AWG43_1 and AWG38 defined a ∼1.5 Mbp sequence in the reference sequence assembly, containing 29 gene models, 17 described as high confidence genes (HC) and 12 as low confidence genes (LC) (Fig. 1c). We found that this interval was smaller, only ∼1 Mbp, in the RAC875 genome (Fig. 1c, Supplementary Fig. 1).

**Fig. 1.**
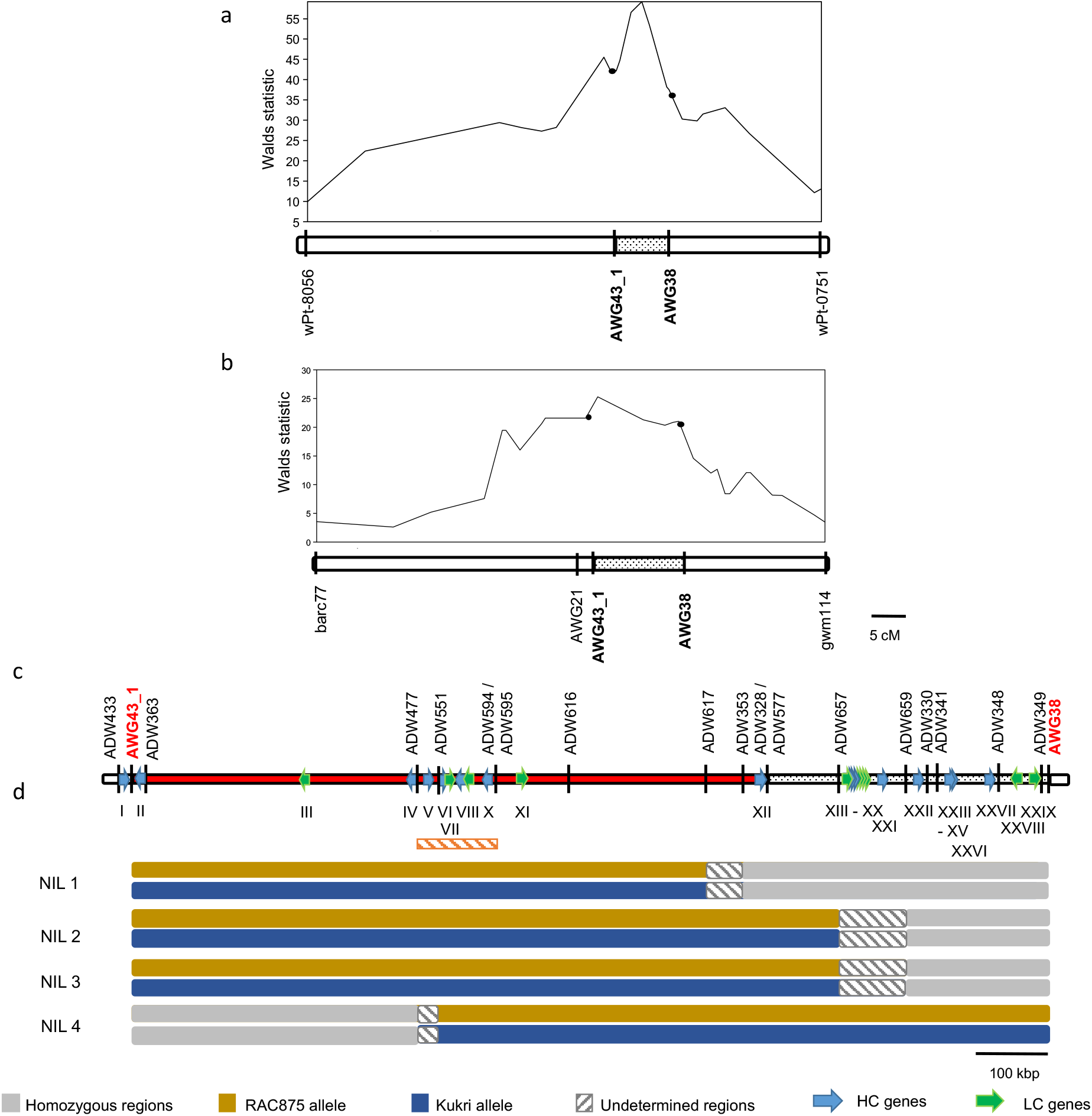
Genomic structure of *qYDH.3BL* in wheat. **a**, QTL analysis of the Drysdale x Gladius recombinant inbred lines (RILs) in 5 field trials in Australia in 2009 and 2010. **b**, QTL analysis of RAC875 x Kukri (RILs) in 4 field trials conducted in 2011 and 2012 in Ciudad de Obregon, Mexico, delineating the QTL peak between the markers AWG43_1 and AWG38. **c**, Physical interval of *qYDH.3BL* in RAC875 defined an interval of ∼ 1 Mbp. HC: high confidence gene models, LC: low confidence gene models. The orange, hatched rectangle highlights the interval where RAC875 local assembly has been done. **d**, Genotypes of RAC875 x Kukri RILs with residual heterozygosity at *qYDH.3BL* used to generate near-isogenic lines (NIL).

The QTL was highly significant for yield in Mexican field trials with heat treatment (by late planting with irrigation by flooding)^29^. These trials were conducted in the northern part of Mexico in the Sonora region where the climate is hot and dry: temperatures at flowering time averaged 25 - 28°C for conventional sowing and 30 - 35°C for late sowing; there is no rainfall during the growing season and water is provided by irrigation; the soil profile is deep with an effective soil depth of 110 - 120 cm^34^. The QTL was most significant when flood irrigation and heat stress were applied to the crop which suggested that the QTL gave an advantage in deep soil where water is stored at depth.

To identify the genes underlying the QTL, higher genetic map resolution was required and new recombinants needed to be grown in conditions where the QTL has the strongest effects. As southern Australian soils are shallow, we used a deep soil-mimic platform to test the hypothesis that a deep soil profile, characteristic of the Mexican environments, triggered *qYDH.3BL* expression. Forty-four and 20 RAC875 x Kukri RILs previously grown in Mexican field trials^29^ were planted in the deep soil platform in August and September 2014, respectively; late planting dates enabled us to phenotype the plants under combined drought and heat stress (Supplementary Fig. 2 and 3). The analysis of the September trial showed a significant positive effect of the RAC875 allele on most measured traits, increasing early vigour, stem biomass, plant height, spike length and biomass, grain and spike number per plant, spikelets per spike, chlorophyll content and seminal roots number, by at least 19 % (Table 1). No effects were observed on stomatal density, nor on carbon and nitrogen isotope discrimination in mature grains (data not shown). These results showed that *qYDH.3BL*’s effects in the deep soil-mimic platform were similar to those observed in Mexican field trials. A quantitative analysis of soil water potential and air temperature effects on *qYDH.3BL* had previously found that the allele effect was correlated to temperatures^30^. The RAC875 allele increased individual seed weight, biomass and harvest index when temperatures were above 25°C around flowering time. As expected, we found that the RAC875 allelic effects were larger under the hotter conditions experienced by plants sown in September (Table 1) compared to those planted in August (Supplementary Fig. 4).

**Table 1.** Effects of the RAC875 allele at *qYDH.3BL* in 20 RAC875 x Kukri RILs and four lines (RAC875, Kukri, Drysdale and Gladius) sown in September 2014 in the deep soil-mimic platform (n = 3).

| Phenotypic traits | Allelic effect* (%) | *p* value |
| --- | --- | --- |
| Early vigour per plant | 19.3 | 0.03 |
| Plant height | 20.8 | 0.03 |
| Spike number per plant | 20.8 | 0.03 |
| Grain number | 21.3 | 0.02 |
| Spike biomass | 21.3 | 0.02 |
| Relative chlorophyll content in flag leaf | 21.6 | 0.02 |
| Spikelet per spike | 25.1 | 0.01 |
| Seminal roots | 27.2 | 0.01 |
| Stem biomass | 27.6 | 0.01 |
| Spike length | 29.0 | 0.01 |

\*allelic effect calculated as the % of variance explained by the RAC875 compared to the Kukri allele.

Out of 2,000 RAC875 x Kukri RILs, we found 30 lines with recombination breakpoints in the interval delimited by the AWG43_1 and AWG38 markers and grew them in the deep soil-mimic platform with heat stress to fine map *qYDH.3BL*. In 2015, maximum temperatures at flowering time ranged between 34.8°C and 38.8°C (Supplementary Fig. 2). Spike length and biomass, stem biomass and early vigour were significantly higher in plants carrying the RAC875 allele at the ADW594 – ADW577 interval (Fig. 2a). Forty-four Drysdale x Gladius RILs were also phenotyped in dry and hot conditions, using the deep soil-mimic platform. The single-marker analysis showed the positive allele from Drysdale associated with increases in flag leaf length and single grain weight that narrowed the interval in this population to markers AWG 43_1 and AWG38 (Fig. 2b).

**Fig. 2.**
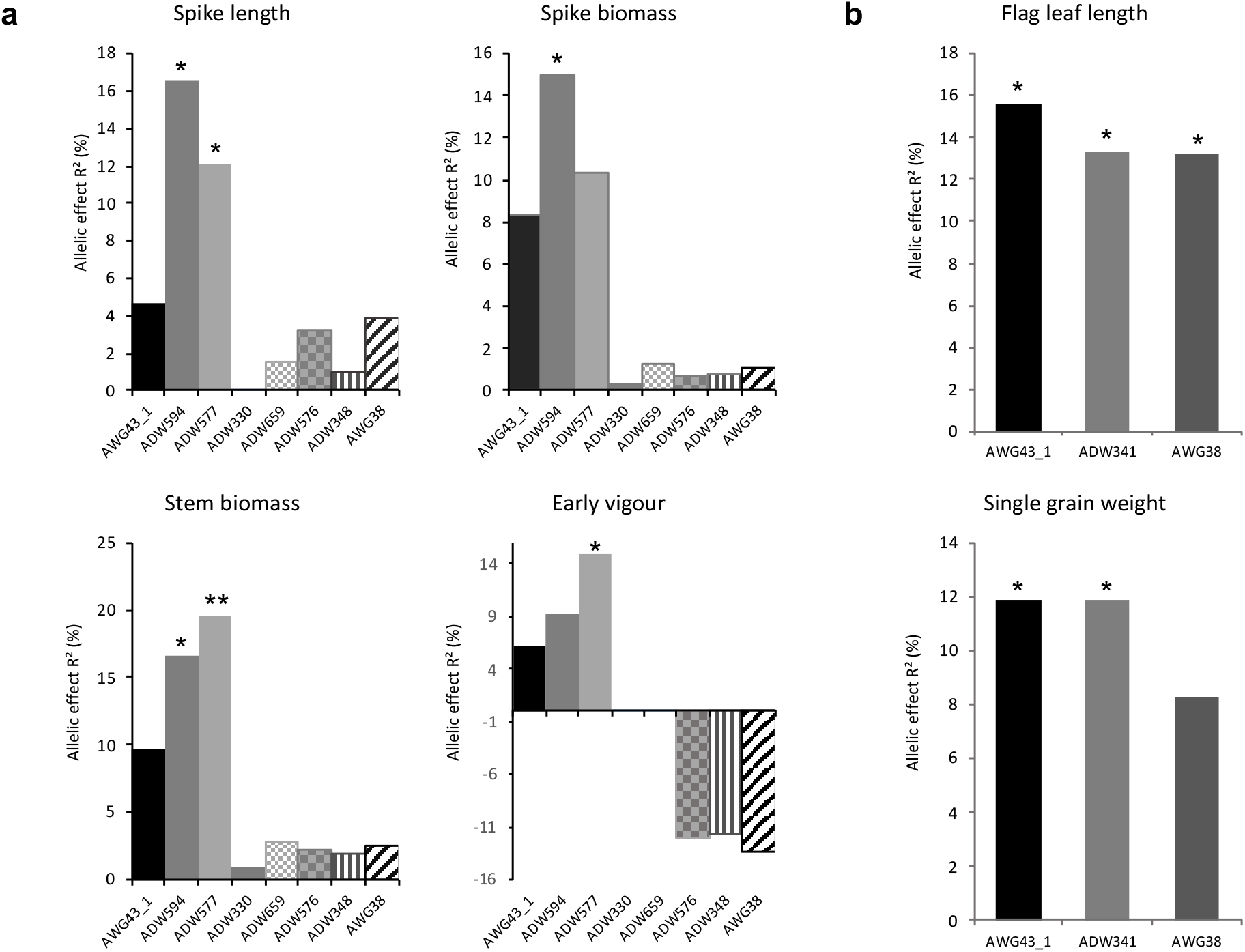
Single marker analysis in 30 RAC875 x Kukri RILs and 44 Drysdale x Gladius RILs segregating for *qYDH.3BL* and grown in dry and hot conditions in the deep-soil platform (2015). **a**, Positive effects were associated with the RAC875 allele while negative effects were associated with the Kukri allele. **b**, Positive effects were associated with the Drysdale allele while negative effects were associated with the Gladius allele. * : p value < 0.05; ** : p value < 0.01

The eight recombination breakpoints in the QTL region in the 30 RAC875 x Kukri RILs enabled us to fine map *qYDH.3BL* to a ∼690 Kbp sequence interval delimited by markers ADW594 and ADW577 which contained 12 annotated genes in the Chinese Spring reference sequence (Table 2). We also performed a local assembly of the RAC875 genomic sequence where no additional or missing annotated genes were detected in the interval compared to the Chinese Spring reference sequence (Supplementary Fig. 1).

**Table 2.** Genes annotated in the Chinese Spring reference sequence v1.0 690 Kbp interval of *qYDH.3BL*. The putative function of each gene was retrieved by homology with rice and brachypodium. HC = high confidence gene; LC = low confidence gene

|  | Gene ID | Position (bp) | Putative function based on rice orthologues | E value | Putative function based on brachypodium orthologues | E value | Expression data* |
| --- | --- | --- | --- | --- | --- | --- | --- |
| I | TraesCS3B01G570900HC | 802845950 - 802848503 | LOC_Os10g33910.1_Mitochondrial import inner membrane translocase subunit Tim16 | 4.1 e-42 | Bradi2g61480_mitochondrial import inner membrane translocase subunit TIM16 | 1.7 e-55 | expressed |
| II | TraesCS3B01G571000HC | 802848874 - 802852363 | LOC_OS02g17280_Gamma-secretase subunit APH-1B | 4.3 e-131 | Bradi3g10110_endopeptidase activity | 8.5 e-135 | expressed |
| III | TraesCS3B01G851200LC | 803363524 - 803364407 | - | - | - | - | - |
| IV | TraesCS3B01G572500HC | 803511461 - 803513157 | LOC_Os11g14410_Polygalacturonase | 8.5 e-142 | Bradi2g57427_Polygalacturonase / Pectinase | 8.2 e-87 | - |
| V | TraesCS3B01G572600HC | 803523701 - 803525213 | LOC_Os01g72720_expressed protein | 8.9 e-11 | Bradi2g61275_PF03478 Protein of unknown function | 4.8 e-56 | expressed |
| VI | TraesCS3B01G572700HC | 803540158 - 803540583 | - | - | Bradi1g36100_Glycosyl hydrolase, subfamily GH 28 | 1.6 e-22 | - |
| VII | TraesCS3B01G851300LC | 803580461 - 803581032 | LOC_Os01g72620_expressed protein | 1.6 e-22 | - | - | - |
| VIII | TraesCS3B01G572800HC | 803583095 - 803583731 | LOC_Os01g59540_GRF zinc finger family protein | 1.1 e-4 | Bradi5g21617 | 1.5 e-24 | - |
| IX | TraesCS3B01G851400LC | 803586310 - 803587263 | LOC_Os07g42000_expressed protein | 1.8 e-14 | - | - | - |
| X | TraesCS3B01G572900HC | 803628595 - 803630088 | LOC_Os01g03170_seven in absentia protein family protein | 3.3 e-34 | Bradi1g27970_ubiquitin-protein ligase activity | 6 e-57 | expressed |
| XI | TraesCS3B01G851500LC | 803715648 - 803716915 | - | - | - | - | - |
| XII | TraesCS3B01G573000HC | 804008796 - 804009926 | LOC_Os01g03170_seven in absentia protein family protein | 2.7 e-31 | Bradi1g27970_ubiquitin-protein ligase activity | 1.7 e-48 | - |

\*Expression data extracted from www.wheat-expression.com and in-house database

As no further recombinants were found in our RIL collections, we developed near-isogenic lines (NIL) from heterozygous inbred families with recombination breakpoints spanning the QTL interval using RAC875 x Kukri RILs with residual heterozygosity between AWG43_1 and AWG38 (Fig. 1d). Following crossing, these families segregate for the locus in contrasting pairs but are otherwise near isogenic. In 2017, the lines were phenotyped under combined drought and heat treatment in the deep soil-mimic platform for yield components and main tiller sap flow, a proxy for plant transpiration^35^. To avoid the confounding effects of phenology, we selected NIL2, 3 and 4 with similar flowering time as the RILs previously studied. In NIL2 family, the lines carrying the RAC875 allele had increased spike and stem biomass, increased grain weight per plant and increased average number of spikelets per spike compared to those carrying the Kukri allele. In NIL3 family, the lines carrying RAC875 allele had increased single grain weight compared to those carrying the Kukri allele. In NIL4 family, the lines carrying the RAC875 allele had increased spike and stem biomass, grain number, average number of spikelets per spike and spike number compared to those carrying the Kukri allele (Table 3). Mass sap flow was, on average, significantly lower in all NILs carrying the RAC875 allele compared to those carrying the Kukri allele at the QTL over the period (Fig. 3). The regression of normalized sap flow mass against mean daily temperatures was significant (P= <0.05) in all lines even though the proportion of the variance explained was low (adj. R^2^ = 0.11–0.33). Whilst the difference was not significant early on during temperate days (average temperature 20.5°C, Fig. 3a) or later in development (grain filling) during several cool days (average temperature 17.5°C, Fig. 3b), there was a significant difference in sap flow between RAC875-allele NILs versus Kukri-allele NILs during heat stress (average temperature 31.5°C) (Fig. 3c). At this time, sap flow in NILs carrying the RAC875 allele was always lower than in NILs carrying the Kukri allele as the RAC875-allele lines failed to increase sap flow in response to increasing temperature in comparison with the alternative allele.

**Table 3.** Analysis of variance of the NIL carrying the RAC875 or Kukri allele at *qYDH.3B* and grown in wheelie bins under drought and heat stress. The values represent the mean ± standard deviation.

| Traits | Early vigour<br>(cm<sup>2</sup>) | Spike<br>biomass (g) | Stem<br>biomass (g) | Grain weight per<br>plant (g) | Grain number | Number of<br>spikelets per<br>spike | Single grain<br>weight (mg) | Average spike<br>length | Spike<br>number | Tiller<br>number | Plant height<br>(cm) |
| --- | --- | --- | --- | --- | --- | --- | --- | --- | --- | --- | --- |
| NIL2-AA | 25.52 ± 3.59 | 1.95 ± 0.74 | 2.74 ± 0.72 | 1.08 ± 0.39 | 49.63 ± 16.30 | 18.04 ± 0.48 | 19.93 ± 3.81 | 9.23 ± 0.30 | 1.78 ± 0.83 | 3.67 ± 0.87 | 52.22 ± 3.85 |
| NIL2-BB | 23.75 ± 4.05 | 1.21 ± 0.34 | 1.81 ± 0.43 | 0.64 ± 0.20 | 37.75 ± 7.70 | 17.48 ± 0.47 | 16.65 ± 3.2 | 8.87 ± 0.51 | 1.38 ± 0.52 | 3 ± 0.53 | 48.39 ± 4.91 |
| df | 16 | 16 | 16 | 16 | 15 | 16 | 16 | 16 | 16 | 16 | 16 |
| F-test | ns | * | ** | * | ns | * | ns | ns | ns | ns | ns |
| NIL3-AA | 19.40 ± 2.81 | 1.77 ± 0.46 | 3.01 ± 0.78 | 1.17 ± 0.31 | 40.63 ± 10.2 | 16.25 ± 0.94 | 29.17 ± 5.23 | 8.89 ± 2.98 | 1.58 ± 0.51 | 3.62 ± 0.51 | 54.68 ± 5.11 |
| HIF3-BB | 17.66 ± 2.63 | 1.46 ± 0.97 | 2.58 ± 1.26 | 0.83 ± 0.60 | 31.08 ± 17.6 | 16.07 ± 1.17 | 22.5 ± 5.65 | 8.64 ± 2.94 | 1.64 ± 0.84 | 3.5 ± 1.02 | 50.9 ± 6.01 |
| df | 26 | 25 | 25 | 24 | 23 | 25 | 24 | 25 | 25 | 26 | 25 |
| F-test | ns | ns | ns | ns | ns | ns | ** | ns | ns | ns | ns |
| NIL4-AA | 24.81 ± 2.71 | 2.38 ± 0.91 | 3.05 ± 0.9 | 1.29 ± 0.55 | 67.36 ± 13.74 | 16.3 ± 0.39 | 17.10 ± 2.80 | 8.03 ± 0.41 | 2.75 ± 0.62 | 3.58 ± 0.79 | 54.05 ± 4.56 |
| NIL4-BB | 22.49 ± 2.76 | 1.69 ± 0.54 | 2.26 ± 0.55 | 0.96 ± 0.32 | 51.45 ± 14.44 | 15.74 ± 0.81 | 18.56 ± 2.92 | 7.88 ± 0.44 | 1.91 ± 0.7 | 3.27 ± 0.79 | 51.73 ± 5.62 |
| df | 22 | 22 | 22 | 21 | 21 | 22 | 22 | 21 | 22 | 22 | 22 |
| F-test | ns | * | * | ns | * | * | ns | ns | ** | ns | ns |

**Fig. 3.**
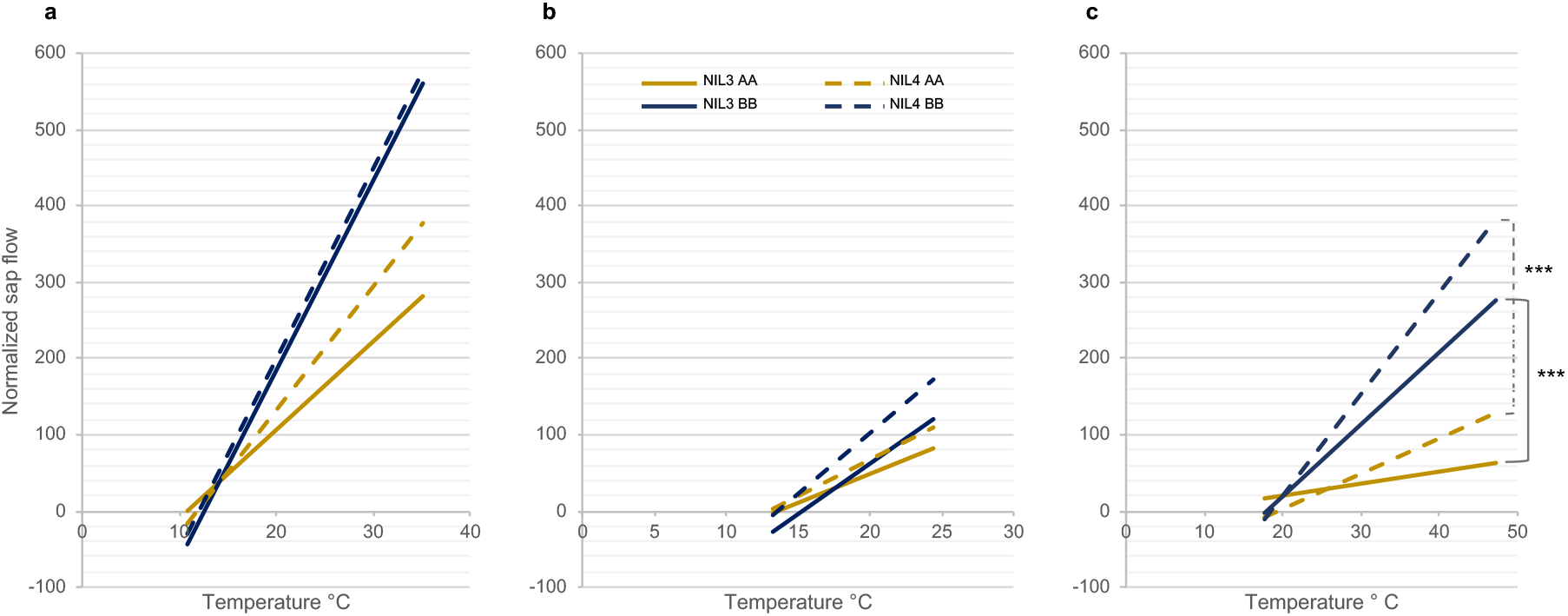
Regression of normalized sap flow of NIL3 and NIL4 families against temperature. **a**, temperate days. **b**, cool days. **c**, hot days. AA = RAC875 allele (gold), BB = Kukri allele (blue). Differences between AA and BB alleles were non-significant on temperate and cool days. *** : p<0.001 between AA and BB alleles on hot days.

To understand whether variation in water use was related to the observed increase in early vigour for the RAC875 allele at *qYDH.3BL*, we grew the NIL3 and 4 allele pair plants on an automated, gravimetric watering system with real-time imaging of early vegetative growth and measurement of water used per plant in unstressed conditions. Early vigour was increased in NIL4 lines carrying the RAC875 allele (Fig. 4) consistent with the allelic effect observed in the RIL experiments in deep soil (Table 1) and the increase in final biomass observed for this NIL (Table 3), but the effect was not significant in NIL3. In unstressed conditions, NIL4 with the RAC875 allele also had increased water use compared with the Kukri allele in complete contrast with its reduced sap flow during heat stress. The water use index (kpixel/ ml H_2_O) of NIL4 was increased in plants carrying the RAC875 compared with the Kukri allele, showing that this line’s early increase in biomass did not demand increased water use in unstressed conditions, *i.e.* that the RAC875 allele at the locus conferred both increased early vigour and increased water use efficiency in unstressed conditions.

**Fig. 4.**
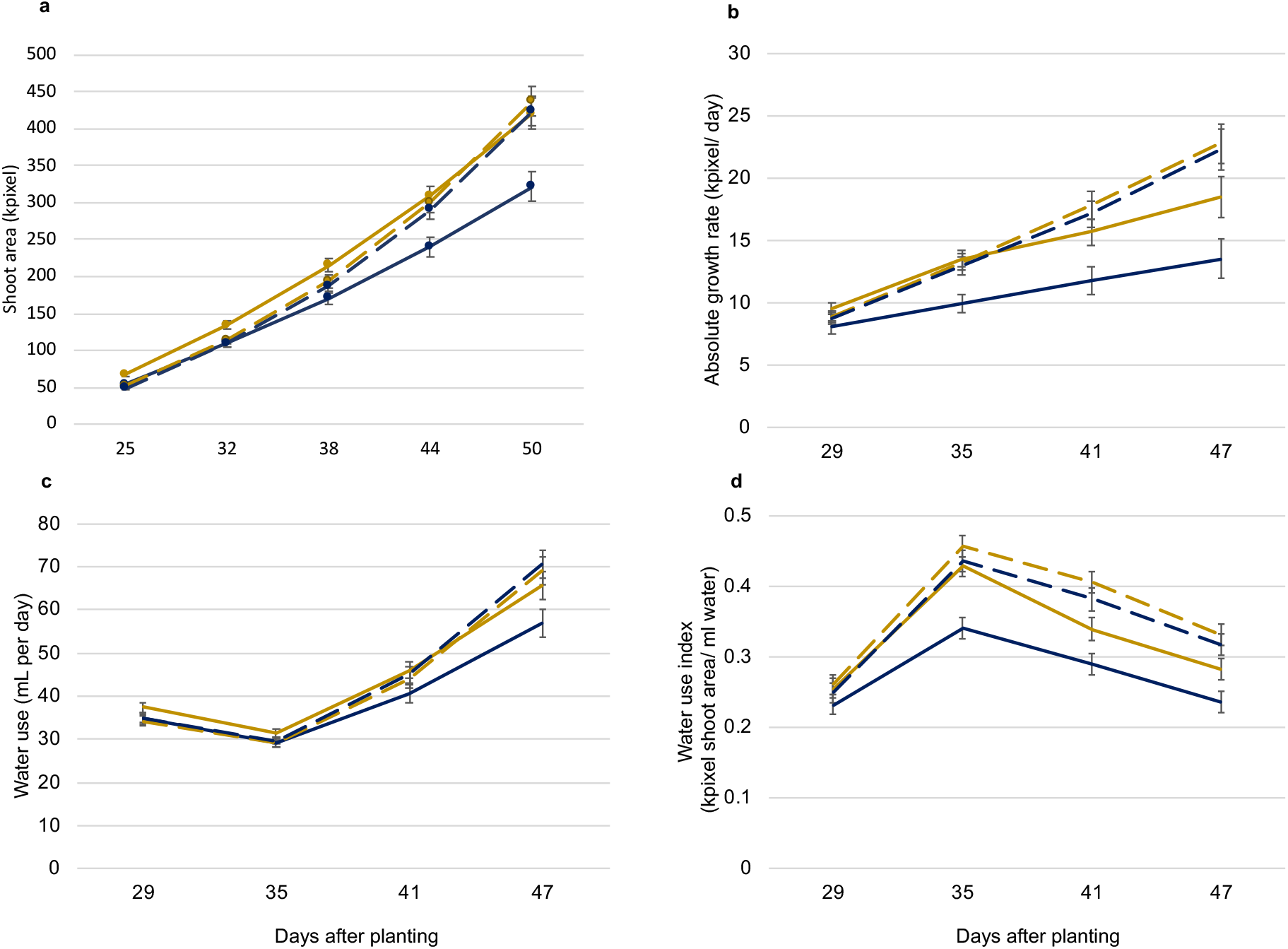
Early vigour and water use of NIL grown in unstressed conditions. **a**, estimated shoot area (kpixel) and **b**, absolute growth rate (kpixel/ day) of young NIL3 (solid line) and NIL4 (hatched line) family plants. AA = RAC875 allele (gold), BB = Kukri allele (blue). **c**, water use (mL/ day) and **d**, water use index (shoot area kpixel/ mL).

We then analysed the sequence of the 12 gene models present in the QTL interval delineated by ADW594 and ADW577 markers using whole genome sequencing datasets of RAC875, Kukri, Gladius and Drysdale. Using shotgun sequencing data of the four parental lines, no non-synonymous variation was identified in the protein sequences encoded by the 12 genes at the locus when the RAC875-Drysdale vs Kukri-Gladius translated sequences were compared. Therefore, functional variation was most likely to lie at the gene expression level. We next studied the expression profile of these gene models in the four NIL pairs that contrasted for yield components in deep soil. We did not find any evidence of gene expression for the four low confidence gene models, either in the wheat-expression database^36^ (http://www.wheat-expression.com) or in our in-house database^37^ that displays RNA-seq expression data in five tissues at three developmental stages^38^ (Table 2). We concluded that these sequences were likely pseudogenes. As expected, we found evidence of gene expression for the eight genes annotated as high confidence (Table 2).

We measured the expression of these eight genes in the four NIL pairs, expecting a difference in expression between the RAC875 and Kukri alleles for the gene(s) responsible for *qYDH.3BL* effect. As the QTL was associated with early vigour, when no treatment had been applied, we studied gene expression in seedlings grown in unstressed conditions. The expression profiles of genes I, II, IV, V, VI, VIII and XII in RAC875 and the NILs carrying the RAC875 allele were not significantly different from Kukri and the lines carrying the Kukri allele (Supplementary Fig. 5a-g). Gene X (*TaSINA 3B*) was the only gene with a differential expression pattern both between the parental lines and among the NILs (Fig. 5a). The gene was consistently highly expressed and statistically different in the lines containing the Kukri-Gladius allele compared to the lines containing the RAC875-Drysdale allele, in both shoot and root tissues of seedlings, except for NIL2 in shoot tissue (Fig. 5a and 5e). By contrast, gene X homeologs on chromosomes 3A and 3D had the same expression profile in the two parental lines and the NILs independently of the allele that they contained and the tissue analyzed (Fig. 5b and 5d).

**Fig. 5.**
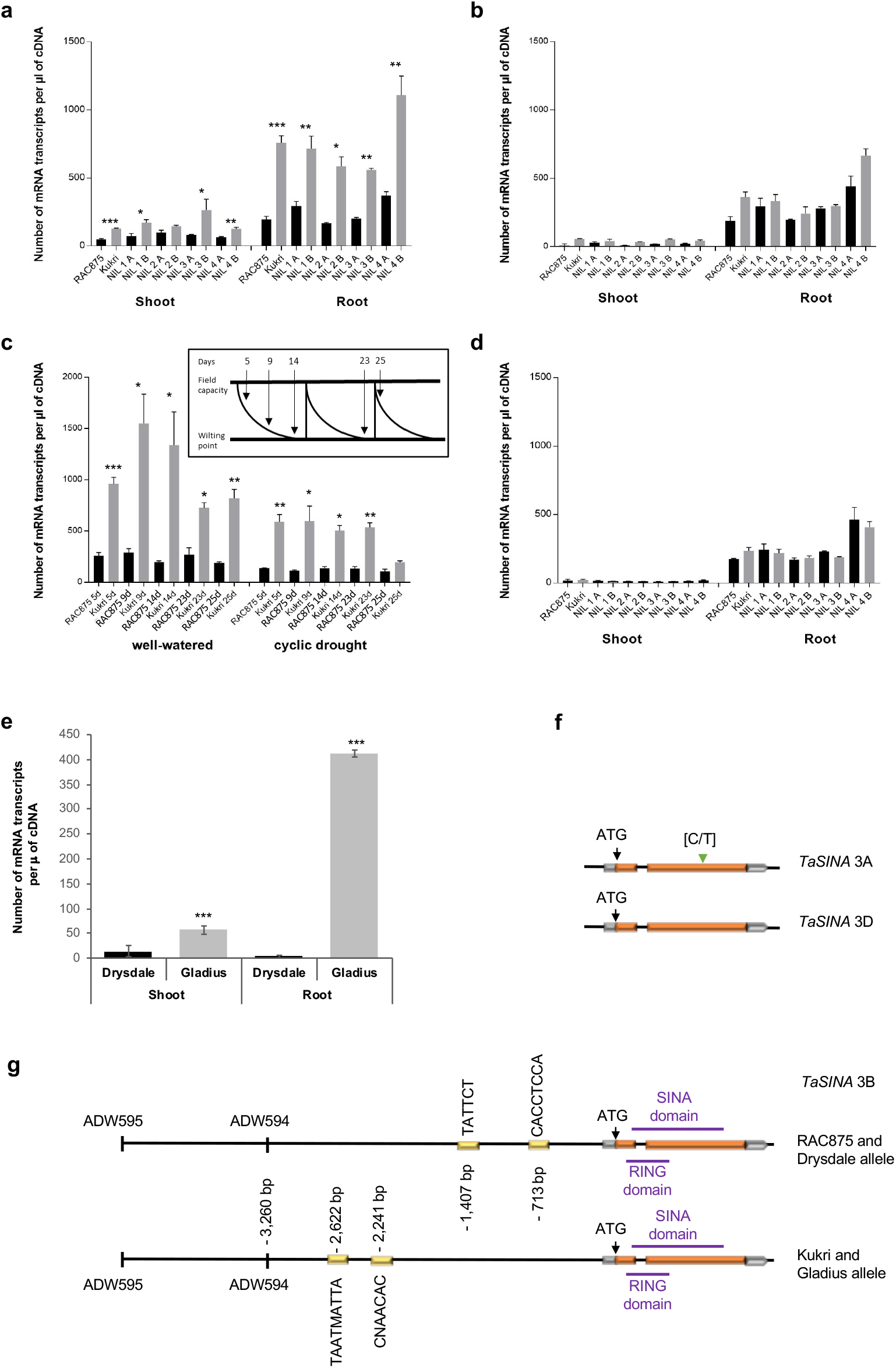
Expression analysis and gene structure of *TaSINA* and its homeologs. **a, e** Expression analysis of *TaSINA* 3B in seedling shoot and root tissues of RAC875, Kukri and NILs (**a**) and in Gladius and Drysdale (**e**). **c**, Expression of *TaSINA* 3B in RAC875 and Kukri under well-watered and cyclic drought conditions described in ^39^. Expression analysis of *TaSINA* 3A (**b**) and *TaSINA* 3D (**d**). “A’ corresponds to the RAC875 allele and the “B” corresponds to the Kukri allele. *: p value < 0.05; ** : p value < 0.01; *** : p value < 0.001. **f**, Exon-intron structure of *TaSINA* 3A and 3D. **g**, Exon-intron structure and promoter annotation of *TaSINA* 3B. Yellow boxes indicate the position of cis-acting elements present in the promoter region; green triangle, SNP position inducing the alanine to valine amino acid change; orange boxes represent the exons and the grey boxes the UTRs.

We also analyzed the expression profile of gene X in cDNA of RAC875 and Kukri grown in well-watered conditions and cyclic drought^39^. Gene X was significantly over-expressed in Kukri compared to RAC875 in both well-watered and drought treatments (Fig. 5c). Based on homology with rice and *Brachypodium*, gene X is homologous to a *seven in absentia (SINA)* gene encoding an E3 ubiquitin ligase protein, hereafter called *TaSINA*.

The translated Sanger sequencing of the *TaSINA* gene revealed two amino acid substitutions between the RAC875 and Kukri protein sequences that were not predicted to affect the protein function. We also found sequence polymorphisms in the promoter region with the predicted *cis*-acting elements CANBNNAPA and HDZIP2ATATHB2 present in Kukri and absent in RAC875 (Fig. 5g). In RAC875 and Drysdale promoter sequences, the insertion of a CCAC domain 713 bp upstream of the start codon created a SBOXATRBCS (CACCTCCA) motif and a −10PEHVPSBD (TATTCT) motif which were absent in the Kukri and Gladius promoter sequences (Fig. 5g). The sequences of *TaSINA* homeologs on chromosomes 3A and 3D were monomorphic between RAC875 and Kukri. The alignment of the TaSINA predicted protein sequence and its homeologs revealed an amino acid change in the protein sequence of the 3A copy (Fig. 5f). A SNP in the coding sequence of the 3A copy introduced the substitution of an alanine to a valine at the 227 amino acid position that was predicted to be deleterious for the protein function (Fig. 5e). This suggested that the 3A protein might be non-functional in both RAC875 and Kukri, although we did find evidence of gene expression in the roots.

We studied the worldwide allelic distribution of *TaSINA* alleles using the two closest SNP markers (ADW594 and ADW595). Alleles were homogeneously distributed among the lines, with 53 % of the accessions carrying the same allele as RAC875 (Fig. 6). We looked at the origins and years of release of the accessions in each allelic group to identify a potential pattern of selection (Supplementary Table 1). The RAC875 allele was over-represented in accessions originating from CIMMYT in Mexico and in Australia compared to the Kukri allele which was more abundant in the germplasm originating from the Middle-East, South America and Asia (Fig. 6). Both alleles were evenly represented in European and African wheats. The Kukri allele was present in historical wheat varieties such as Du Toits, Federation, Ward’s Prolific and Hudson Early Purple Straw, released before 1900, and also in a large proportion of landraces (Supplementary Table 1). The RAC875 allele appeared later in the panel’s chronology, with most of the lines released after 1950, and seemed to follow the migration of germplasm released by CIMMYT during the green revolution of the 1960s.

**Fig. 6.**
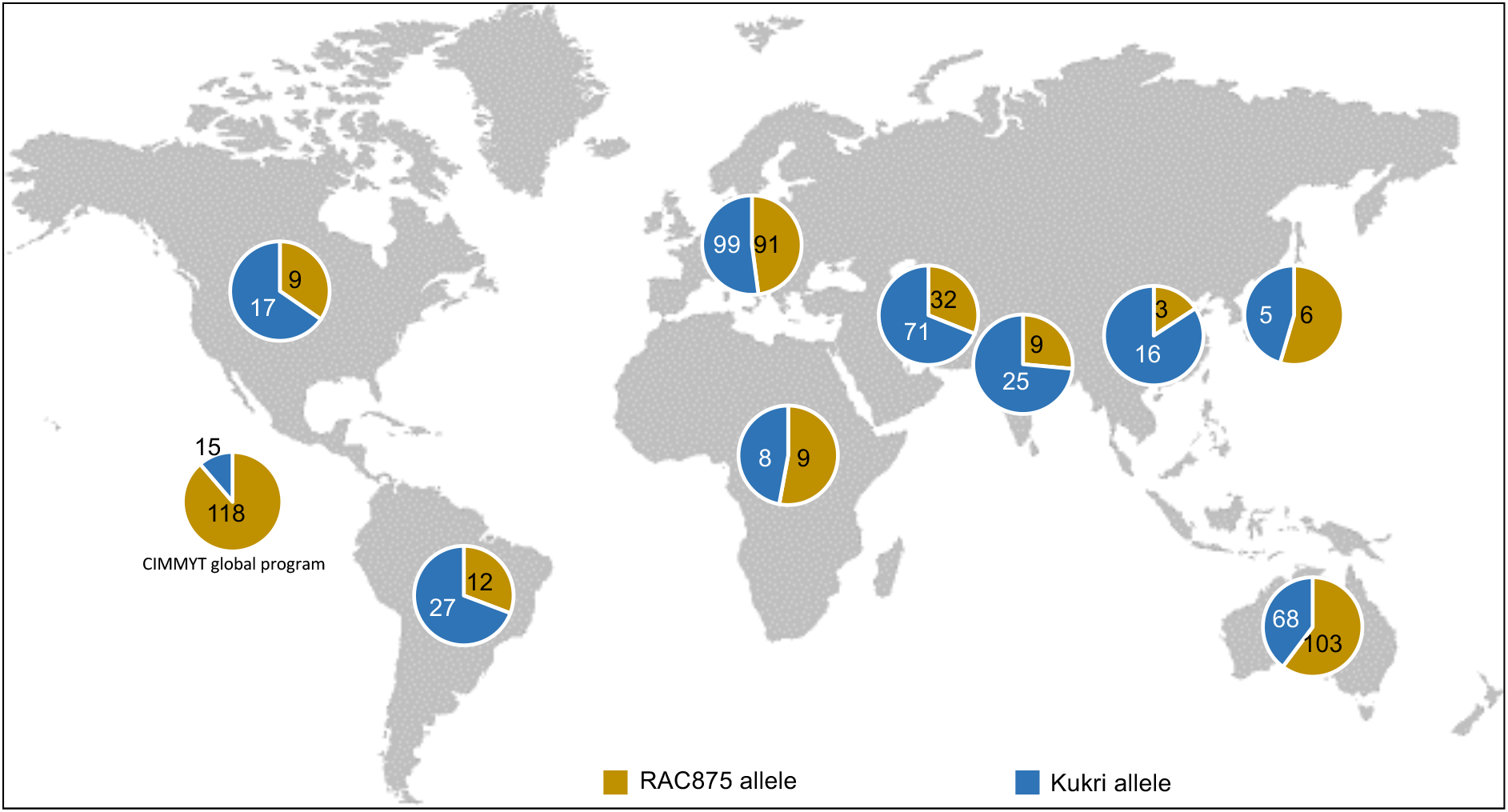
Allelic distribution at *TaSINA* in a diversity panel of 743 *T. aestivum* accessions genotyped with KASP markers ADW594 and ADW595.

## Discussion

The RAC875 allele at *qYDH.3BL* increases early vigour, seminal root number, plant height, spike length, above ground biomass, stem and spike biomass, spike number, number of spikelets per spike and total grain number in deep soil. The association of *qYDH.3BL* with early vigour suggests that the mechanisms underlying the QTL are effective at an early stage of plant development, before abiotic stress is encountered. Early vigour is an important trait associated with water uptake, preventing water loss from soil evapotranspiration to the profit of transpiration^40^. We hypothesize that this increased early growth in plants carrying the RAC875 allele had no significant effect under optimal conditions, as observed in southern Australian field trials in 2009^29^, but was beneficial to the plant when temperatures increased.

As *qYDH.3BL* had been previously associated with differences in canopy temperature^11^, a surrogate for plant evaporative cooling, we used sap flow sensors to evaluate the transpiration rates of whole plants^35^. We observed that sap flow responded to daily temperature variations, but this response was abolished in lines carrying the RAC875 allele during heat stress late in development. These results are consistent with the observations of Izanloo et al.^28^ on the water-management behaviour of RAC875 and Kukri parental lines under cyclic-drought. They concluded that RAC875 reduced its water consumption at early stages of drought stress for later consumption when the stress become more severe. Here, the RAC875 allele increased plant biomass and water use at early, vegetative stages of development and reduced transpiration in response to heat late in development without any negative impact on yield components.

By fine mapping the QTL and by expression analysis of eight genes in the QTL interval, we found that only the *TaSINA* gene showed consistent variation between lines carrying contrasting alleles at the QTL. Beyond those identified in the reference assembly, no other gene was present in the interval when we locally assembled RAC875 genomic sequence. Therefore, *TaSINA* is a strong candidate for the QTL effect.

*SINA* genes encode for an E3 ubiquitin ligase protein that forms a complex with the E1 ubiquitin-activating enzyme and the E2 ubiquitin-conjugating enzyme which ubiquitinate target proteins for degradation^41^. *SINA* genes were first reported in Drosophila associated with the development of photoreceptor cell, R7, localized in the Drosophila eye^42^. Analyses of the gene family in plants identified several copies of *SINA* in *Arabidopsis thaliana*^43^, *Populus trichocarpa*^44^, *Oryza sativa*^43^, *Zea mays*^43^, *Physcomitrella patens*^43^, *Medicago truncatula*^44^ and *Solanum lycopersicum*^45^ with roles in plant development and abiotic stress tolerance^46,47^. No *SINA* gene has previously been reported in bread wheat. In Arabidopsis, *SINAT5* ubiquitinates a *NAC1* transcription factor to regulate the growth signalling hormone auxin^48^. Arabidopsis plants overexpressing *SINAT5* had fewer lateral roots compared to the wild type, whereas expression of a dominant-negative mutant (substitution Cys49-Ser) induced more lateral roots^48^. The ectopic expression of this dominant negative form of *SINAT5* in *M. truncatula* reduced root nodulation, but both root and shoot growth were more vigorous in young (*in vitro*) plants after 20 days and in older plants, after eight weeks’ growth^44^. We observed similar results in our experiments when Kukri had fewer primary and seminal roots compared to RAC875 (Table 1): *TaSINA* was more strongly expressed in Kukri compared to RAC875 and early vigour and biomass were increased in wheat plants when *TaSINA* was not expressed (Table 1, Fig. 4). No change in *TaSINA* expression was observed for RAC875 and expression was reduced in Kukri only after prolonged cyclic drought (25d) (Fig. 3b). This suggests that *TaSINA* expression is relatively insensitive to stress and that heat tolerance in wheat is more likely conferred by increases in growth and water use when *TaSINA* expression is reduced.

The promoter sequence of RAC875 *TaSINA* contained a *cis*-acting element, SBOXATRCS (CACCTCCA) also called S box, absent in the promoter of Kukri *TaSINA* (Fig. 5). This motif is a putative ABSCISIC ACID INSENSITIVE-4 (ABI4) -binding site which negatively regulates the *Conserved Modular Arrangement 5* genes, *CMA5*, in Arabidopsis in response to sugar and ABA signals^49^. ABA is a phytohormone that plays a central role in different physiological processes of plants, mediates gene expression in response to environmental stimuli and regulates ABA-dependent responses in heat stress^50^. Sugar signalling also plays an important role in mediating plant responses to environmental stimuli and mediates gene expression associated to photosynthesis, carbon and nitrogen metabolism and secondary metabolisms^51^. We hypothesize that the presence of the S box element and ABI4 binding site, in the promoter sequence of *TaSINA*, contributes to the down-regulation of its expression in RAC875 and might affect ABA and sugar signalling. In Kukri, the absence of the motif would prevent the regulation of *TaSINA* and lead to gene expression and, consequently, the degradation of its target proteins.

Following genotyping of 743 worldwide wheat accessions, we found that the RAC875 allele was over-represented in the CIMMYT germplasm suggesting that the allele had been selected through breeding for high yield under heat stress. The RAC875 allele was unexploited for wheat breeding until the 1950s when it appears to have been recruited by the CIMMYT wheat programme in Mexico and into Australia with the introduction of CIMMYT material. It is surprising that RAC875 allele is not more prevalent in India and other Asian countries given that CIMMYT germplasm, either as parents or directly as cultivars, has been estimated to account for over 65% of production in India. This allele has not yet been exploited in other regions where heat was not considered a threat until recently. With the increasing occurrence of heat stress events, this allele could be beneficial in other regions with deep soil such as in Europe and in the Punjab in India.

## Materials and Methods

### Plant materials

A set of 160 recombinant inbred lines (RILs) of the cross between RAC875 (RAC-655//SR21/4*Lance/3/4*Bayonet) and Kukri (Madden/6*RAC-177//Grajo/76-ECN-44) were used for the fine mapping of *qYDH.3BL*. A set of 70 RAC875 x Kukri RILs were selected for their contrasting yield in the 2011^29^ and 2012 field trials conducted in Mexico (Supplementary Table 2) and based on their recombination point in the QTL interval.

Four near isogenic families derived from QTL-specific residual heterozygous lines from RAC875 x Kukri RIL were developed to further study the mechanism underlying *qYDH.3BL*. A second set of 44 RILs of the cross between Drysdale (Hartog*3/Quarrion) and Gladius (RAC875/Krichauff//Excalibur/Kukri/3/RAC875/Krichauff/4/RAC875//Excalibur/Kukri) (DG) was also studied. The lines were selected based on their recombination breakpoint in the QTL interval and their extreme yield values during the 2009 and 2010 Mexican and New South Wales field trials^52^. The four parental lines RAC875, Kukri, Drysdale and Gladius were included in each trial.

Nulli-tetrasomic lines cv. Chinese Spring (LV-Szechuan) of the chromosome group 3 were used to test the specificity of chromosome-specific primers. The method used to obtain nulli-tetrasomic lines was previously described^53^.

A diversity panel combining 743 landraces and modern varieties of hexaploid wheat was used to identify the distribution of alleles at *qYDH.3BL*. 544 accessions were spring wheat types originating from world-wide locations with more than one quarter from Australia. We also used the INRA core collection which contains 372 worldwide accessions of winter wheat mostly from Europe^54^. Pedigree, year of release and geographical origin of the lines were retrieved from the wheat pedigree portal, www.wheatpedigree.net, and personal communications (Peter Langridge, Margaret Pallotta).

### Genotyping and genetic map construction

DNA was extracted as described previously^55^. Gene-based markers were designed using the whole genome sequencing (WGS) data of the parental lines (RAC875, Kukri, Drysdale and Gladius) generated by BioPlatforms Australia using Illumina HiSeq sequencing technology^33^. Sequence reads (100 bp) were aligned against the reference sequence IWGSC RefSeq v1.0 of Chinese Spring (https://www.wheatgenome.org/) using DAWN. DAWN is a web interface that integrates multiple datasets including IWGSC RefSeq v1.0, RNA sequencing data of five tissues at three developmental stages^39^ and WGS data of the parental lines. SNPs within *qYDH.3BL* interval were identified with a minimum of five reads per line at a SNP position. As regions with a high density of reads are more likely to be repetitive elements, SNPs with a coverage higher than 50 reads were discarded.

Kompetitive Allele Specific PCR (KASP) markers were designed with the Kraken software (LGC genomics, Middlesex, UK) on sequences of at least 100 bp spanning a SNP. Two allele-specific forward primers were designed for each SNP in combination with a common reverse primer. Each forward primer had a tail attached at the 5’end specific to a fluorophore. The hybridization of one specific forward primer to the target sequence allowed the pairing of the fluorophore present in the KASP mix to the primer tail releasing the quencher allowing emission of fluorescence^56^. KASP assays were performed using the KASPline (https://www.lgcgroup.com). KASP markers were prefixed AWG- and ADW- (Supplementary Table 3).

The genotyping data of the KASP markers were added to RAC875 x Kukri and Drysdale x Gladius RILs genetic maps previously generated by Bonneau et al.^29^. Genetic maps were constructed using ASMap package^57^ available in the software R. Genetic distance between each marker was calculated using the Kosambi mapping function^58^ which is based on a new algorithm named MSTmap^59^. This algorithm can determine the markers order efficiently by establishing a minimum spanning tree. Two markers, AWG594 and AWG595, were used for studying the allelic distribution in the wheat diversity panel.

### Deep soil phenotyping platform

Wheelie bins (100 × 57.5 × 51 cm) were filled with a mixed medium (1/3 coco peat, 1/3 sand and 1/3 clay). Twenty-five plants were grown per bin, in a 5 rows × 5 columns grid with 10 cm space between each plant. Trials were conducted at the Waite Campus, Urrbrae, Australia, in a tunnel with a polyurethane cover. Meteorological data, air temperature and relative humidity were recorded with a data logger (Kongin KG100, China). Gypsum blocks (MEA, Magill, SA, Australia) were placed in the bins at two depths (10 and 40 cm) to record soil water potential (Supplementary Fig. 3).

In 2014 and 2015, RILs and parental lines were grown in a resolved latinized incomplete block using CycDesigN^60^, plus one filler plant placed in the centre of the bin. The design took into account the position of the lines according to a west/east axis and split the bins between an inner and outer layers. Two trials were conducted in both 2014 and 2015. In 2014, RAC875 x Kukri RILs were sown later than normal planting season (44 RILs sown on 11^th^ of August and 20 RILs sown on 8^th^ of September, instead of May) to phenotype plants under combined drought and heat treatment. Plants were well-watered during the first month for both treatments; watering was stopped at booting stage for stress treatments. In 2015, both trials were conducted under dry and hot conditions: 30 RAC875 x Kukri RILs were sown on the 10^th^ of July, and 44 Drysdale x Gladius RILs were sown on the 18^th^ of August.

In 2017, the NILs were grown in a single plant plot design with 15 replicates each for the parental lines RAC875 and Kukri and 54 replicates of each pair of NIL. The design was developed using the function “prDiGGer” in the DiGGer package in R for partially replicated designs (Supplementary Fig. 6). Replicates were distributed within the bins so that the same genotype was not present twice in the same row or column. In 2017, the plants were sown on 31^st^ of July, watering was maintained until late booting in the beginning of October and then stopped.

### Phenotypic evaluation in the deep soil-mimic system

Plant developmental stages were scored using the Zadoks’ scale around anthesis^61^. Early vigour was scored using two methods. In 2014, during the first month after planting, photographs were taken for each bin every week and early vigour were scored using a 1 (less vigorous) to 5 (more vigorous) comparative scale. In 2015, early vigour was monitored by measuring the total leaf area when a plant reached the four-leaves stage. The lengths and widths of each leaf on the main tiller were measured to determine the individual leaf area. The total leaf area was calculated as the sum of the leaf area (leaf width x leaf length x 0.8) of the second and third leaves^62^. Chlorophyll content of the flag leaf was estimated with a SPAD-502L meter (Ozaka, Japan) at three stages: booting, anthesis and grain filling. Three measurements per flag leaf at each stage were taken to estimate the average value. Flag leaf area was measured from booting to flowering time using the LI-3000C portable leaf area meter (LI-COR Inc., USA) in 2014. In 2015 flag leaf length and width were measured at flowering stage with a ruler. Stomatal density was measured by taking a leaf imprint of the flag leaf adaxial face at anthesis using translucent nail polish on a glass slide^63^. The slides were then analysed by microscopy using the Leica AS-LMD Laser Microdissection Microscope (Wetzlar, Germany). Three pictures per sample were taken and the guard cells number was counted for each.

Tiller number was recorded at four-leaf stage and before harvesting. Tiller abortion was calculated at the end of the trial by subtracting the spike number from the tiller number. After harvest, spikes were manually counted to evaluate the number of spikes per plant. Spikes and stem biomass were weighed separately, and harvest index was determined by dividing grain biomass by total above-ground biomass. Spike length was measured using a ruler before counting the number of spikelets per spike for each spike. Seed number was measured using a seed counter (Pfueffer GmBH, Germany) and then weighed to determine single grain weight. In 2014, carbon and nitrogen isotopes discrimination of mature grains were measured on five grains per plant. Grains were dried for one week at 60°C and finely ground to 1 to 4 mg of powder stored in tin capsules pressed (Sercon, gateway Crewe, UK). The samples were sent to the University of California, Davis Stable Isotope Facility, University of California, for analysis using continuous flow Isotope Ratio Mass Spectrometer (IRMS).

Root traits were also evaluated in the 2014 experiment in deep-soil bins by digging out the roots at 10 cm depth after harvesting. The number of both seminal and nodal roots were measured. Pictures of the root system were taken for each plant and analysed using the Image J software^64^. The root growth angle of the angle formed by seminal roots and soil surface was measured.

### Measurement of plant water use using sap flow sensors

In 2017, SF-4/5 Micro Stem Sap Flow sensors (Edaphic Scientific, Port Macquarie, NSW, Australia) were used in the wheelie bins on the NILs to measure ascending sap flow through the main stem and calculate plant water use^35^. The measurements are non-destructive and continuous. Each sensor contains a heater located between two temperature probes. Data were automatically collected every 15 min after the initial warm up of the stem by the heater for five min. The output voltage was proportional to the difference of temperatures between the two probes. Sensors were installed on plants from mid-October when watering was stopped until harvest and placed between the first and second node. Data were recorded from the beginning of grain filling. Sensors were isolated with aluminium foil to avoid temperature fluctuations. Sensors were calibrated by measuring stomatal conductance (g_s_, mmol.m^−2^.s^−1^) of plants every two hours from pre-dawn until after sunset with an AP4 leaf porometer (Delta-T devices, Cambridge, UK). Collected raw sap flow data were normalized by setting a constant zero as the average of readings for five hours during each preceding night and then averaged for every hour to calculate the mean average of the sap flow data for each allele within a NIL pair. The data were then plotted with the daily mean temperature. We also calculated the total transpiration rate per day and compared it within each NIL pair between the lines carrying the RAC875 allele and those with the Kukri allele.

### Early vigour imaging and water use analysis

Uniformly sized seeds of the NIL3 and NIL4 allelic pairs (n=6) were sown in 150 mm pots (one seed per pot) in cocopeat compost and grown in a climate-controlled glasshouse containing an automated gravimetric watering and imaging platform previously described^65^ with day/night temperatures of 22/15°C. The plants were arranged in two lanes × 18 positions using a randomised complete-block design. After 25 d, images of each plant from each NIL and allelic pair were captured daily for the succeeding 25 d using the LemnaTec-Scanalyser 3D platform. Projected shoot area (PSA kpixel) was estimated as the sum of the areas from three RGB camera views comprising two side views at 80° angles and one view from above. Water use was computed using the smoothed values of watering amounts (mL) and water use index for a given interval was the ratio in change between PSA and total water use in that interval, with higher values corresponding to higher water use efficiency.

### QTL analysis

The multi-environment QTL analysis of the RAC875 x Kukri RILs combined four field trials conducted in Mexico, Ciudad de Obregon, in 2011^29^ and 2012^52^ and was performed as described previously^29^. For the single-marker analysis, we generated the best predicted unbiased estimates (BLUEs) for each trait using the model (1) and the asremlPlus package^66^. The normal distribution of the BLUEs was then evaluated using the Shapiro-Wilk test^67^. Single-marker analysis at each marker position was performed to test association with the studied traits by a one-way ANOVA in R. For the trait which did not follow a normal distribution, a Kruskal-Wallis test was used^68^.

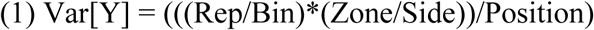

### Anchoring of *qYDH.3BL* onto the wheat reference sequence

Flanking markers at the QTL were used to find sequence similarity with the reference sequence RefSeq v1.0 of Chinese Spring by BLASTN (e-value −10; # hit return =100). The 1.5 Mbp sequence delimited by AWG43_1 and AWG38 was extracted using Fetch-seq, an in-house sequence retriever that allows the extraction of sub-sequences from the IWGSC RefSeq v1.0 using coordinates. Gene annotations were retrieved using the coordinated interval in the browser available on the URGI website (https://wheat-urgi.versailles.inra.fr/). Genomic sequences of the annotated genes were used to find similarities with *Brachypodium distachyon* v3.1 and the *Oryza sativa* v7 reference genomes to identify putative gene function by BLASTN (default settings) in phytozome (https://phytozome.jgi.doe.gov/).

### Targeted assembly of the region in RAC875

Initial, stringent alignments of WGS reads of RAC875^33^ indicated that approximately 150 kbp region in Chinese Spring RefSeq 1.0 (from adw477 to adw594) may either be absent or highly divergent in RAC875. Further, more relaxed alignments of RAC875 paired-end (PE) and mate-pair (MP) data to RefSeq v1.0 failed to produce any evidence for the region being deleted. Reads and their mates which remained unaligned and those which aligned within the region of interests were k-merized, k=64 using KMC2^69^. We then assembled k-mers occurring three or more times in the input data into unitigs i.e. contigs unambiguously supported by k-mers, using yakat kextend^70^. Unitigs were scaffolded with the complete MP and PE datasets using sspace^71^. The resulting scaffolds were aligned to RefSeq v1.0 using BLASTn to exclude those derived from other regions of the genome and to identify a subset likely to be derived from the region of interest. The total length of 80 scaffolds putatively identified as coming from the region of interest adds up to ∼200 Kbp and the longest scaffold covers 36 Kbp. Scaffolds with a minimum length of 10 Kbp were then annotated using TriAnnot^72^.

### Gene sequencing and promoter analysis

*TaSINA* chromosome specific primers were designed with Primer 3 available in Geneious (Biomatters, Auckland, New Zealand) (Supplementary Table 3). Nulli-tetrasomic lines^53^ specific to the group 3 chromosome were used to test the specificity of the primers through amplification of the amplicon fragment. PCR samples were Sanger sequenced at the Australian Genome Research Facility Ltd (http://www.agrf.org.au/). Oligo sequences were conserved in RAC875, Kukri and Chinese Spring. Promoter sequences of the RAC875 and Kukri *TaSINA* were annotated using the Plant Cis-acting Regulatory DNA Elements (PLACE) database which includes sequence motifs reported as cis-acting regulatory DNA elements^73^. Protein domains, residues and motifs of *TaSINA* were identified using Prosite^74^ and Pfam (http://beta.supfam.org/).

### Gene expression analysis

The wheat-drought experiment cDNA series of RAC875 and Kukri was previously described^39^. Additionally, we used two-week-old seedlings of the NIL pairs for gene expression analysis. Seedlings were grown in an air-conditioned glasshouse in pot trays filled with a potting mix (coco-peat). Total RNA for both root and shoot tissues was extracted using the Spectrum™ Plant Total RNA Kit (Sigma-Alridch, Carlsbad, California, USA) using the manufacturer’s recommended protocol. cDNAs were synthetised using the SuperScript IV First-Strand Synthesis System (Invitrogen, Sigma-Alridch, Carlsbad, California, USA) as per the manufacturer’s instructions and used for qPCR. qPCR primers targeting 70-200 bp amplicon sequences were designed to target the chromosome specific gene copy (Supplementary Table 3). Specificity was tested with nulli-tetrasomic lines. qPCR assays were performed using the Kapa SYBR Fast Universal 2X qPCR Master Mix (Geneworks, Thebarton, South Australia, Australia) on the QuantStudio 6 Flex (Applied Biosystems, Foster City, CA, USA) following these steps: 95 °C for 3 min; followed by 40 cycles at 95 °C for 20 sec, 63 °C for 20 sec, 72 °C for 15 sec. The initial run was then followed by a melting curve at 95 °C for 15 sec, 60 °C for 1 min increasing by 0.05 °C per cycle up to 95 °C. Three technical replicates per sample and per gene and four reference genes including *TaActin, TaCyclophilin, TaGAPDH* (glyceraldehyde-3-phosphate dehydrogenase) and elongation factor (*TaEFα*), were included in the analysis for normalization of gene expression.

## Acknowledgements

This work was supported by the Australian Research Council Industrial Transformation Research Hub for wheat in a hot and dry climate (IH130200027), the Grains Research and Development Corporation, the Government of South Australia and DuPont-Pioneer, USA. PaT would like to thank the University of Adelaide for the Beacon of Enlightenment PhD scholarship. We also acknowledge the use of the facilities and scientific and technical assistance of the Australian Plant Phenomics Facility, which is supported by the Australian Government’s National Collaborative Research Infrastructure Strategy (NCRIS). We would also like to acknowledge BioPlatforms Australia for funding the sequencing of RAC875, Kukri, Drysdale and Gladius wheat lines. We thank Dr Fabio Arsego, Dr Yuan Li, Dr Matteo Riboni, Niharika Sharma, Dimitri Sanchez and Kate Dowling for technical assistance. We acknowledge and thank Florian Veillet who developed some of the markers used, Julian Taylor for preliminary running of the genetic mapping models and Paul Eckermann for the sap flow data normalization script.

## Author contributions

PaT and PK performed the genotyping, QPCR and plant phenotyping experiments, developed the NIL and PaT analysed data. JB developed molecular markers, ran the trial on RIL in Ciudad de Obregon and generated the RIL fine genetic map. CB developed the experimental design of the deep soil-mimic and imaging experiments and wrote the R scripts to analyse the data. RS did the local assembly of RAC875 genomic sequence. UB supervised the DAWN analysis and the annotation of the genomic sequences performed by PaT. PT analysed the sap flow data. PaT, JB, PL, PT, and DF designed the experiments, interpreted the results and co-wrote the paper.

