## Supplementary figures and tables for "The wheat *Seven in Absentia* gene is associated with increases in biomass and yield in hot climates" (from another version of this preprint: version 2, 2019-08-13)

**a**

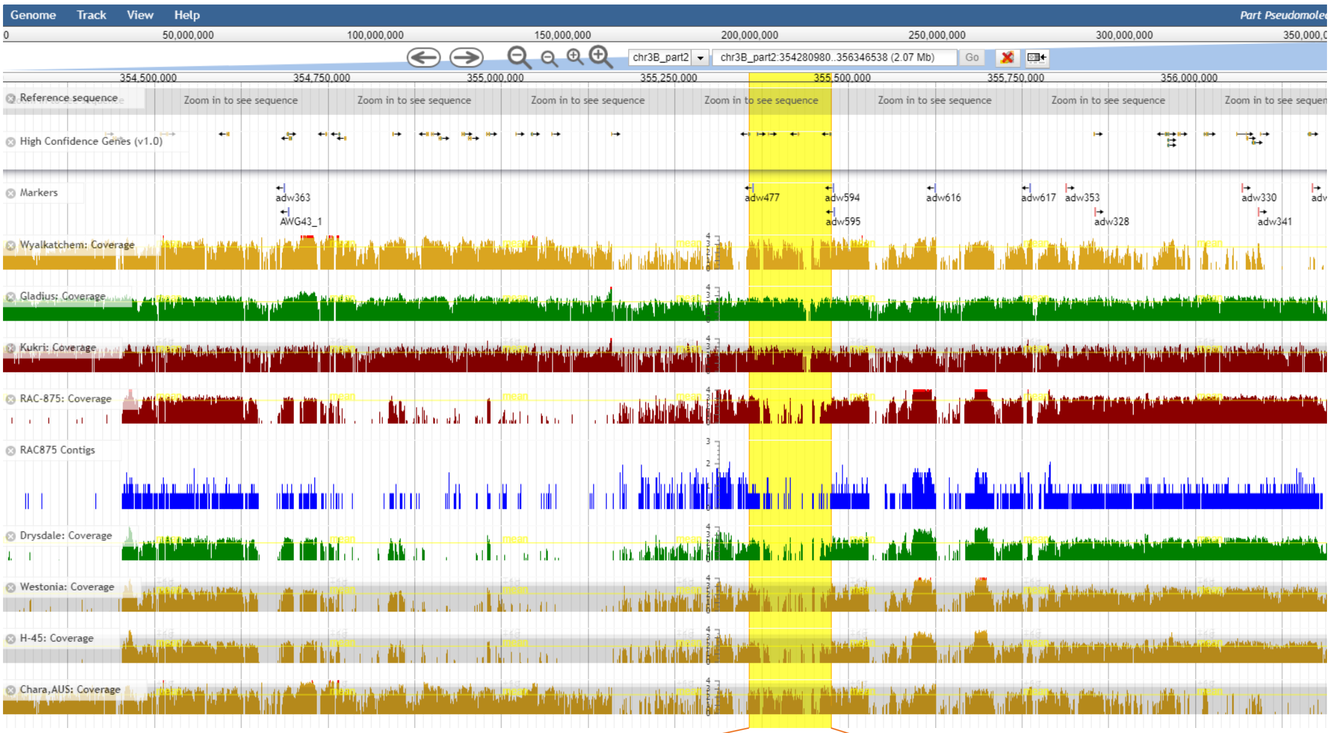


**b**

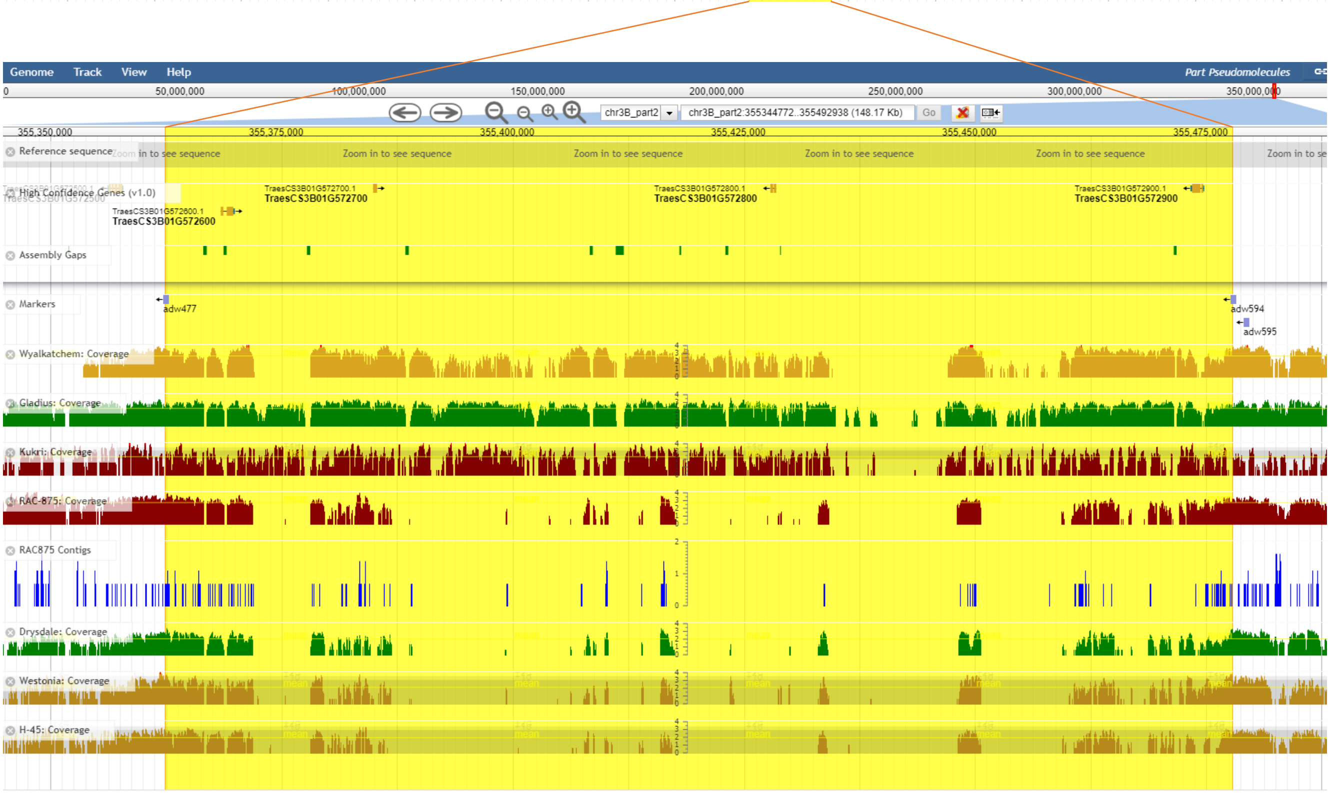


**Supplementary Fig. 1 | Genetic diversity around the *qYDH.3BL* QTL visualised in DAWN**[75]. (**a**) The QTL is located between markers adw477 and adw594 and highlighted in yellow. Read coverage is displayed for eight wheat accessions with Kukri and RAC875 in dark red, Gladius and Drysdale in green. The track in blue represents the minimap2 alignment of assembled RAC875 contigs to the reference genome. (**b**) Close-up of the QTL showing the sparsity of reads from RAC875 and also RAC875 derived contigs. A track showing the location and relative size of assembly gaps has been added (3rd from top).

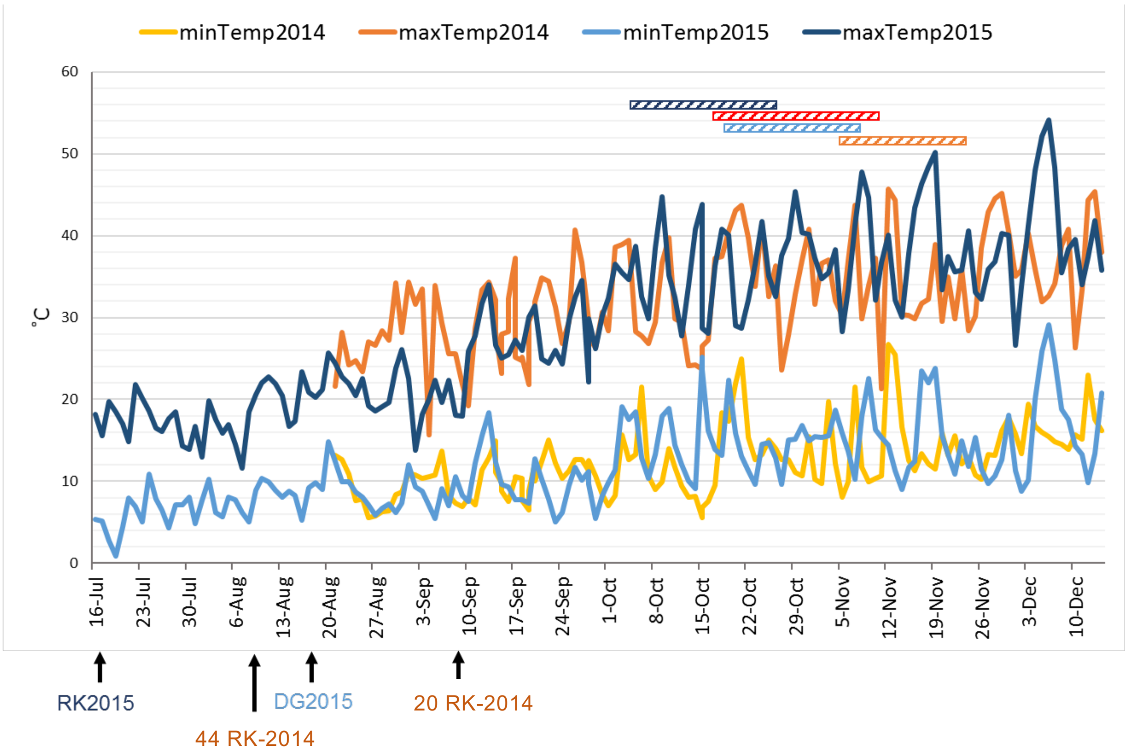


**Supplementary Fig. 2 | Air temperatures during 2014 and 2015 experiments using the RAC875 x Kukri and Drysdale x Gladius RILs in the deep soil platform in a polytunnel.** Black arrows indicate the sowing date for each experiment and lines. In 2014, 44 RAC875 x Kukri RIL (RK) were planted in August and 20 RK in September. In 2015, 30 RK lines were planted in July and 20 Drysdale x Gladius RIL (DG) in August. Flowering periods have been highlighted for each experiment: red for Aug-2014; orange for Sept-2014; light blue for DG2015; dark blue for RK2015.

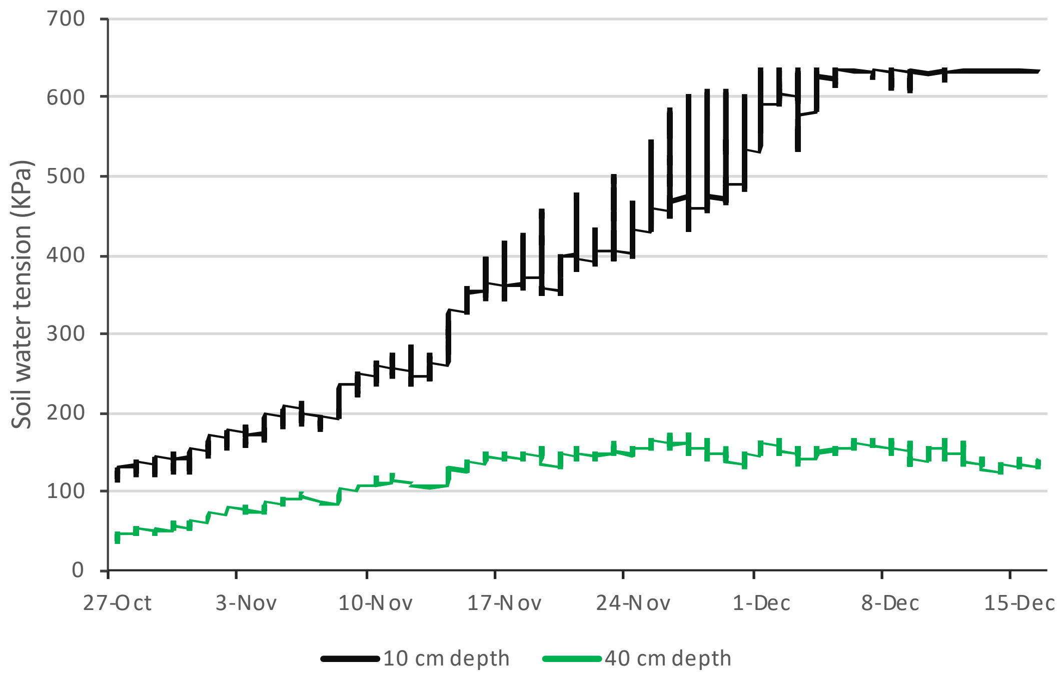


**Supplementary Fig. 3 | Average soil water tension during the grain-filling period when drought was imposed in the deep soil platform.** RAC875 x Kukri RILs and parental lines were planted in August in three bins.

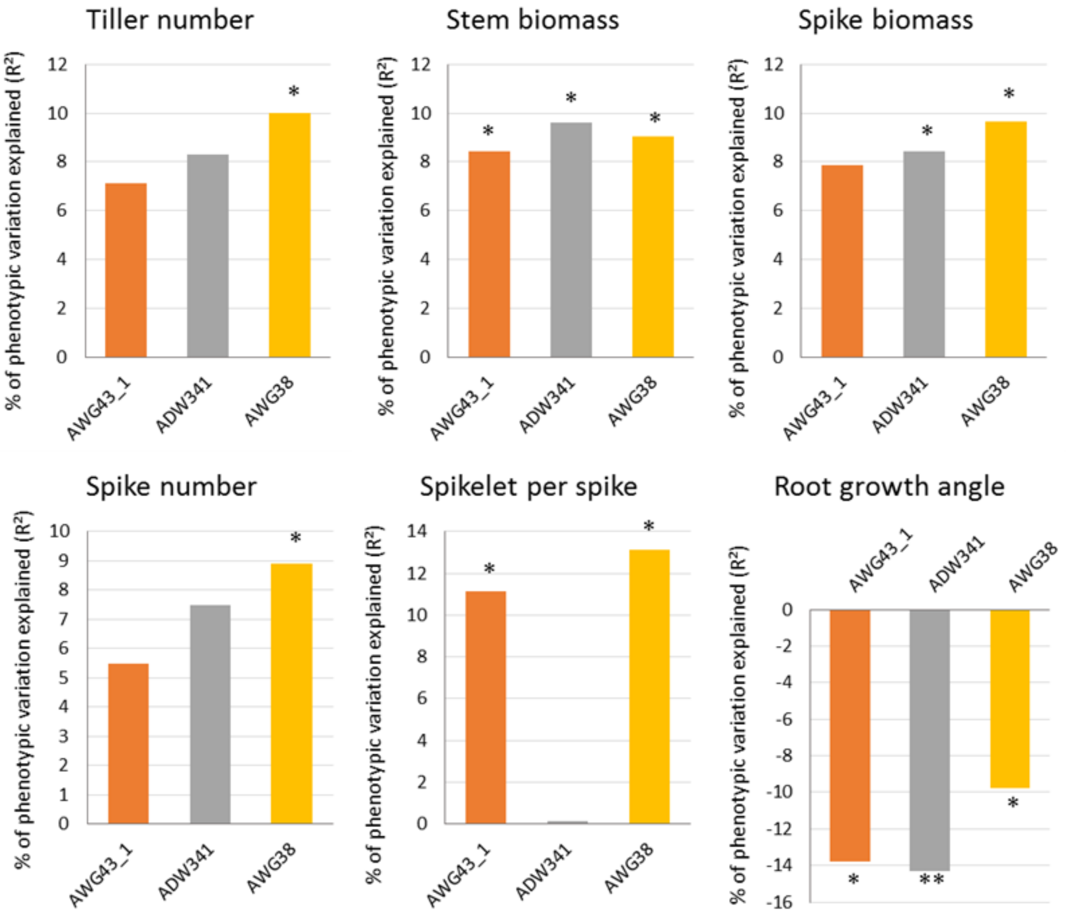


**Supplementary Fig. 4 | Single marker analysis of 44 RAC875 x Kukri RILs phenotyped in the deep soil platform sown in August 2014.** Positive effect indicates that the RAC875 allele is associated with the trait while a negative effect indicates that the Kukri allele is associated with the trait.

* : p value < 0.05; ** : p value < 0.01;

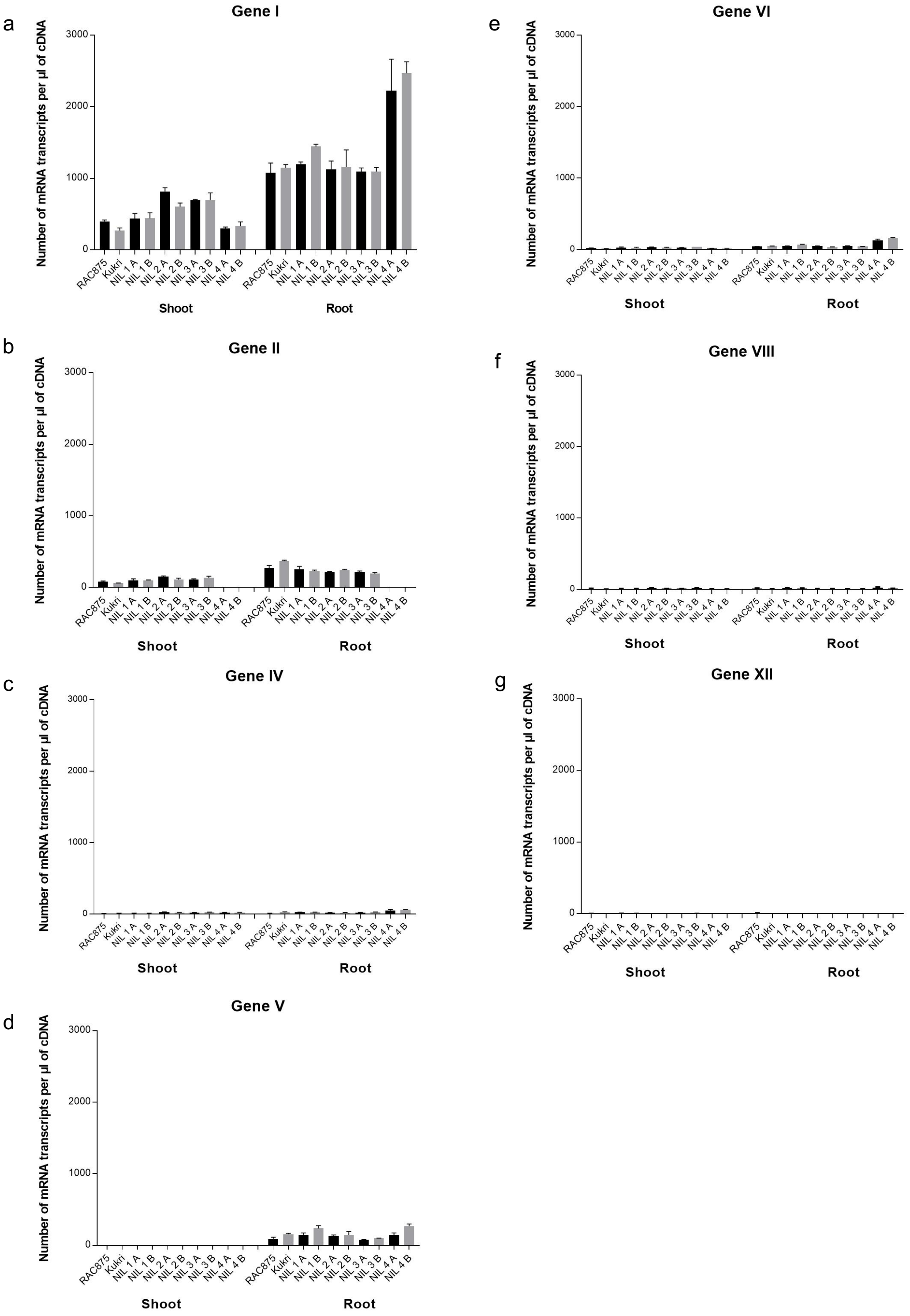


**Supplementary Fig. 5 | Expression of seven genes annotated in the interval delimited by ADW594 and ADW577.** No significant differences in the expression profile of these genes could be found between parents and among the NILs. “A”: NIL contains the RAC875 allele; “B”: NIL contains the Kukri allele.

| | | | | |
|---|---|---|---|---|
| NIL 5A | NIL 2A | NIL 3B | NIL 3A | NIL 5B |
| NIL 1B | NIL 5B | NIL 5A | NIL 1A | NIL 4B |
| NIL 3B | NIL 2B | NIL 2A | Kukri | NIL 4A |
| NIL 4A | NIL 3A | NIL 2B | NIL 4B | NIL 5A |
| RAC875 | NIL 1A | NIL 1B | NIL 4A | Kukri |

**Supplementary Fig. 6 | Example of the randomized design developed for the deep-soil mimic wheelie bin experiments showing one of the bins in 2017 with parental lines RAC875 (allele Aa) and Kukri (allele B) and NIL containing either allele.**

**Supplementary Table 2.** Descriptions of the environment in Ciudad de Obregon (Mexico) where the RAC875 x Kukri RILs field trials were conducted, showing water supply from rainfall and/or irrigation and RAC875 x Kukri DH and RI lines (from Bonneau, 2012[52]).

| Environ-ment [a] | Latitude | Longitude | Altitude (M) | Lines tested [b] | Sowing density seed $m^{-2}$ | Rainfall and/or irrigation mm | Mean temperature around flowering time [c] | |
|---|---|---|---|---|---|---|---|---|
| | | | | | | | **Sept** | **Oct** |
| 11_DI_CS | 27˚28' N | 109˚56' W | 38 | 34 DH/77 RI | 200 | 150 | 24.5 | 29.3 |
| 12_DI_CS | 27˚28' N | 109˚56' W | 38 | 27 DH/ 40 RI | 200 | 100 | 24.7 | 26.6 |
| | | | | | | | **April** | **May** |
| 11_FI_LS | 27˚28' N | 109˚56' W | 38 | 34 DH/ 109 RI | 200 | 1050 | 30.7 | 34.8 |
| 12_FI_LS | 27˚28' N | 109˚56' W | 38 | 27 DH/ 40 RI | 200 | 900 | 30.5 | 37.8 |

[a] Mex Mexico, DI drip irrigation, FI for flooding irrigation, LS for late sowing, CS for conventional sowing date.
[b] DH doubled haploid lines, RI recombinant inbred lines.
[c] The temperature data for Mexico were collected at www.agroson.com.mx
