## Supplementary Table 1 for "The wheat *Seven in Absentia* gene is associated with increases in biomass and yield in hot climates" (from another version of this preprint: version 2, 2019-08-13)

| ID | accession name | ADW594 | ADW595 | Origin |
| --- | --- | --- | --- | --- |
| lw248 | TOUZELLE BLANCHE BARBUE | - | A:A | FRANCE |
| lw318 | AXONA | - | A:A | NETHERLAND |
| lw369 | Landrace | - | A:A | TURKMENISTAN |
| lw378 | Tom-Thumb | - | A:A | USA |
| DPG_513 | H 1546 | A:A | A:A | AFGHANISTAN |
| DPG_497 | H 1409 | A:A | A:A | AFGHANISTAN |
| DPG_488 | H 1287 | A:A | A:A | AFGHANISTAN |
| DPG_492 | H 1328A | A:A | A:A | AFGHANISTAN |
| DPG_515 | H 1582 | A:A | A:A | AFGHANISTAN |
| DPG_517 | H 1594 | A:A | A:A | AFGHANISTAN |
| DPG_525 | H 1655 | A:A | A:A | AFGHANISTAN |
| DPG_530 | H 1685 | A:A | A:A | AFGHANISTAN |
| DPG_1263 | H 1288 | A:A | A:A | AFGHANISTAN |
| DPG_491 | H 1314 | A:A | A:A | AFGHANISTAN |
| DPG_512 | H 1542 | A:A | A:A | AFGHANISTAN |
| DPG_535 | H 339 | A:A | A:A | AFGHANISTAN |
| DPG_1266 | H 1326 | A:A | A:A | AFGHANISTAN |
| DPG_1267 | H 1329 | A:A | A:A | AFGHANISTAN |
| DPG_489 | H 1302 | A:A | A:A | AFGHANISTAN |
| lw298 | BAROOTA WONDER | A:A | A:A | AUSTRALIA |
| DPG_442 | Ford | A:A | A:A | AUSTRALIA |
| DPG_399 | DIRK | A:A | A:A | AUSTRALIA |
| DPG_626 | Koda | A:A | A:A | AUSTRALIA |
| DPG_453 | GALA | A:A | A:A | AUSTRALIA |
| DPG_455 | GAMENYA | A:A | A:A | AUSTRALIA |
| DPG_830 | Raven | A:A | A:A | AUSTRALIA |
| DPG_438 | Festiguay | A:A | A:A | AUSTRALIA |
| DPG_849 | SCOUT | A:A | A:A | AUSTRALIA |
| DPG_672 | Madden | A:A | A:A | AUSTRALIA |
| DPG_864 | Songlen | A:A | A:A | AUSTRALIA |
| DPG_315 | COOK | A:A | A:A | AUSTRALIA |
| DPG_856 | Shortim | A:A | A:A | AUSTRALIA |
| DPG_152 | BANKS | A:A | A:A | AUSTRALIA |
| DPG_126 | AROONA | A:A | A:A | AUSTRALIA |
| DPG_875 | SUNECA | A:A | A:A | AUSTRALIA |
| DPG_678 | Matong | A:A | A:A | AUSTRALIA |
| DPG_542 | HARTOG | A:A | A:A | AUSTRALIA |
| DPG_166 | BODALLIN | A:A | A:A | AUSTRALIA |
| DPG_428 | ERADU | A:A | A:A | AUSTRALIA |
| DPG_440 | Flinders | A:A | A:A | AUSTRALIA |
| DPG_157 | Bass | A:A | A:A | AUSTRALIA |
| DPG_961 | WYUNA | A:A | A:A | AUSTRALIA |
| DPG_197 | CD87 | A:A | A:A | AUSTRALIA |
| DPG_308 | Cocamba | A:A | A:A | AUSTRALIA |
| DPG_700 | Meering | A:A | A:A | AUSTRALIA |
| DPG_836 | ROSELLA | A:A | A:A | AUSTRALIA |
| DPG_327 | CRANBROOK | A:A | A:A | AUSTRALIA |
| DPG_671 | MACHETE | A:A | A:A | AUSTRALIA |
| DPG_635 | KULIN | A:A | A:A | AUSTRALIA |
| DPG_874 | SUNCO | A:A | A:A | AUSTRALIA |
| DPG_900 | Tatiara | A:A | A:A | AUSTRALIA |

| | | | | |
|---|---|---|---|---|
| DPG_949 | WILGOYNE | A:A | A:A | AUSTRALIA |
| DPG_603 | JANZ | A:A | A:A | AUSTRALIA |
| DPG_640 | LANG | A:A | A:A | AUSTRALIA |
| DPG_832 | REEVES | A:A | A:A | AUSTRALIA |
| DPG_433 | EXCALIBUR | A:A | A:A | AUSTRALIA |
| DPG_342 | CUNNINGHAM | A:A | A:A | AUSTRALIA |
| DPG_642 | Lark | A:A | A:A | AUSTRALIA |
| DPG_872 | SUNBRI | A:A | A:A | AUSTRALIA |
| DPG_158 | BATAVIA | A:A | A:A | AUSTRALIA |
| DPG_086 | Babax | A:A | A:A | AUSTRALIA |
| DPG_182 | CADOUX | A:A | A:A | AUSTRALIA |
| DPG_837 | Rowan | A:A | A:A | AUSTRALIA |
| DPG_879 | SUNSTATE | A:A | A:A | AUSTRALIA |
| DPG_768 | PELSART | A:A | A:A | AUSTRALIA |
| DPG_194 | CASCADES | A:A | A:A | AUSTRALIA |
| DPG_880 | SUNVALE | A:A | A:A | AUSTRALIA |
| DPG_897 | TAMMIN | A:A | A:A | AUSTRALIA |
| DPG_873 | SUNBROOK | A:A | A:A | AUSTRALIA |
| DPG_192 | CARNAMAH | A:A | A:A | AUSTRALIA |
| DPG_177 | BUCKLEY | A:A | A:A | AUSTRALIA |
| DPG_947 | WESTONIA | A:A | A:A | AUSTRALIA |
| DPG_397 | DIAMONDBIRD | A:A | A:A | AUSTRALIA |
| DPG_183 | CALINGIRI | A:A | A:A | AUSTRALIA |
| DPG_948 | Whistler | A:A | A:A | AUSTRALIA |
| DPG_159 | BAXTER | A:A | A:A | AUSTRALIA |
| DPG_617 | KENNEDY | A:A | A:A | AUSTRALIA |
| DPG_631 | KRICHAUFF | A:A | A:A | AUSTRALIA |
| DPG_537 | H45 | A:A | A:A | AUSTRALIA |
| DPG_859 | Silverstar | A:A | A:A | AUSTRALIA |
| DPG_818 | QAL 2000 | A:A | A:A | AUSTRALIA |
| DPG_474 | GILES | A:A | A:A | AUSTRALIA |
| DPG_960 | WYLAH | A:A | A:A | AUSTRALIA |
| DPG_404 | DRYSDALE | A:A | A:A | AUSTRALIA |
| DPG_306 | CLEARFIELD JNZ | A:A | A:A | AUSTRALIA |
| DPG_863 | SOKOLL | A:A | A:A | AUSTRALIA |
| DPG_421 | EGA_WEDGETAIL | A:A | A:A | AUSTRALIA |
| DPG_425 | ELLISON | A:A | A:A | AUSTRALIA |
| DPG_416 | EGA BLANCO | A:A | A:A | AUSTRALIA |
| DPG_920 | VENTURA | A:A | A:A | AUSTRALIA |
| DPG_418 | EGA CASTLE ROCK | A:A | A:A | AUSTRALIA |
| DPG_462 | GBA HUNTER | A:A | A:A | AUSTRALIA |
| DPG_968 | YOUNG | A:A | A:A | AUSTRALIA |
| DPG_896 | TAMMARIN ROCK | A:A | A:A | AUSTRALIA |
| DPG_395 | DERRIMUT | A:A | A:A | AUSTRALIA |
| DPG_149 | BARHAM | A:A | A:A | AUSTRALIA |
| DPG_420 | EGA_EAGLEHAWK | A:A | A:A | AUSTRALIA |
| DPG_966 | YENDA | A:A | A:A | AUSTRALIA |
| DPG_167 | BOLAC | A:A | A:A | AUSTRALIA |
| DPG_145 | AXE | A:A | A:A | AUSTRALIA |
| DPG_483 | GUARDIAN | A:A | A:A | AUSTRALIA |
| DPG_196 | CATALINA | A:A | A:A | AUSTRALIA |
| DPG_647 | LINCOLN | A:A | A:A | AUSTRALIA |

| | | | | |
|---|---|---|---|---|
| DPG_427 | ENDURE | A:A | A:A | AUSTRALIA |
| DPG_973 | ZEBU | A:A | A:A | AUSTRALIA |
| DPG_329 | CRUSADER | A:A | A:A | AUSTRALIA |
| DPG_435 | FANG | A:A | A:A | AUSTRALIA |
| DPG_707 | MERINDA | A:A | A:A | AUSTRALIA |
| DPG_881 | SUNVEX | A:A | A:A | AUSTRALIA |
| DPG_976 | ZIPPY | A:A | A:A | AUSTRALIA |
| DPG_940 | WAAGAN | A:A | A:A | AUSTRALIA |
| DPG_552 | HTWT_004 | A:A | A:A | AUSTRALIA |
| DPG_141 | AUS29529 | A:A | A:A | AUSTRALIA |
| DPG_460 | GASCOIGNE | A:A | A:A | AUSTRALIA |
| DPG_825 | RAC655 | A:A | A:A | AUSTRALIA |
| DPG_826 | RAC875 | A:A | A:A | AUSTRALIA |
| lw037 | AURORE | A:A | A:A | AUSTRALIA |
| DPG_452 | Gabo | A:A | A:A | AUSTRALIA |
| DPG_456 | GAMMENYA | A:A | A:A | AUSTRALIA |
| DPG_703 | MENDOS | A:A | A:A | AUSTRALIA |
| DPG_819 | QT7475 | A:A | A:A | AUSTRALIA |
| DPG_944 | WAWHT2750 | A:A | A:A | AUSTRALIA |
| DPG_956 | WW1842 | A:A | A:A | AUSTRALIA |
| DPG_958 | WW80 | A:A | A:A | AUSTRALIA |
| lw223 | P4523-80 | A:A | A:A | AUSTRIA |
| lw295 | PRINCE LEOPOLD | A:A | A:A | BELGIUM |
| lw151 | IAS 1 | A:A | A:A | BRAZIL |
| lw092 | COTIPORA | A:A | A:A | BRAZIL |
| lw289 | LE2092 | A:A | A:A | BRAZIL |
| lw010 | 3716-1 | A:A | A:A | BULGARIA |
| lw383 | OGOSTA | A:A | A:A | BULGARIA |
| DPG_478 | GP260 | A:A | A:A | CANADA |
| DPG_704 | Menflo | A:A | A:A | CHILE |
| DPG_965 | YECORA 70 | A:A | A:A | CHILE |
| lw384 | CHAGUAL | A:A | A:A | CHILE |
| DPG_649 | LIVINGSTON | A:A | A:A | CHINA |
| lw311 | TAI911 | A:A | A:A | CHINA |
| DPG_097 | ACADEMIE DE PEKIN | A:A | A:A | CHINA |
| lw113 | DRAVA | A:A | A:A | CROATIA |
| lw319 | ROKYCANSKA_SAMETKA | A:A | A:A | CZECH REPUBLIC |
| lw320 | HANA | A:A | A:A | CZECH REPUBLIC |
| lw109 | DIANA II | A:A | A:A | CZECH REPUBLIC |
| lw097 | Danubia | A:A | A:A | CZECH REPUBLIC |
| lw296 | Solaris | A:A | A:A | CZECH REPUBLIC |
| lw168 | KRAKA | A:A | A:A | DENMARK |
| DPG_084 | 37627 | A:A | A:A | EGYPT |
| lw012 | 6-1-3 | A:A | A:A | EGYPT |
| DPG_656 | LW337 | A:A | A:A | ETHIOPIA |
| DPG_189 | Capelle Desprez | A:A | A:A | FRANCE |
| lw302 | FONDARD CRESPIN | A:A | A:A | FRANCE |
| lw190 | MICHE | A:A | A:A | FRANCE |
| lw033 | ARROMANCHES | A:A | A:A | FRANCE |
| lw304 | BEHERT | A:A | A:A | FRANCE |
| DPG_813 | PROVINS | A:A | A:A | FRANCE |
| lw009 | DI37-12-2 | A:A | A:A | FRANCE |

| | | | | |
|---|---|---|---|---|
| lw075 | CEREALOR | A:A | A:A | FRANCE |
| lw034 | ARTOIS-DESPREZ | A:A | A:A | FRANCE |
| lw107 | DI7215-100 | A:A | A:A | FRANCE |
| lw112 | DRAGON-FRA | A:A | A:A | FRANCE |
| lw141 | GRENIER | A:A | A:A | FRANCE |
| lw154 | JANGO | A:A | A:A | FRANCE |
| lw247 | TF6 | A:A | A:A | FRANCE |
| lw259 | VALDOR | A:A | A:A | FRANCE |
| lw261 | WATTINES | A:A | A:A | FRANCE |
| lw375 | (95-13*BEZOSTAIA)3-3 | A:A | A:A | FRANCE |
| lw376 | HIVERNAL | A:A | A:A | FRANCE |
| lw377 | RINGOT 2 | A:A | A:A | FRANCE |
| lw381 | DI182-9 | A:A | A:A | FRANCE |
| lw024 | AMIFORT | A:A | A:A | FRANCE |
| lw053 | BLASON | A:A | A:A | FRANCE |
| lw286 | (V*C)5-3 | A:A | A:A | FRANCE |
| lw287 | DC147U | A:A | A:A | FRANCE |
| lw297 | ANATOLIE2 | A:A | A:A | FRANCE |
| lw299 | CONCURRENT | A:A | A:A | FRANCE |
| lw305 | ORNICAR | A:A | A:A | FRANCE |
| lw307 | APACHE | A:A | A:A | FRANCE |
| lw308 | BELLOVAC | A:A | A:A | FRANCE |
| lw266 | BORDEAUX 113 | A:A | A:A | FRANCE |
| lw046 | BEL ET BON | A:A | A:A | FRANCE |
| lw102 | DI6402-34-2-4 | A:A | A:A | FRANCE |
| lw104 | DI7005-113-3 | A:A | A:A | FRANCE |
| lw105 | DI7202-103 | A:A | A:A | FRANCE |
| lw227 | POILU DU TARN | A:A | A:A | FRANCE |
| lw233 | RENAN | A:A | A:A | FRANCE |
| lw005 | 25-AO-20/11-60 | A:A | A:A | FRANCE |
| lw269 | RALET | A:A | A:A | FRANCE |
| lw323 | BLE DE REDON BLANC 1/2 LACHE 1 1 | A:A | A:A | FRANCE |
| lw324 | BLE DE REDON GLUMES VELUES 1 | A:A | A:A | FRANCE |
| lw260 | VPM V1-1-2-4R2-3-8-3-2 | A:A | A:A | FRANCE |
| lw029 | ARCHE | A:A | A:A | FRANCE |
| lw326 | AS68VM4-3-2/TJB636 13 | A:A | A:A | FRANCE |
| lw327 | ASVM4/BEAUCHAMP 81B13 | A:A | A:A | FRANCE |
| lw331 | DI9234-11-15 | A:A | A:A | FRANCE |
| lw309 | PALIO | A:A | A:A | FRANCE |
| lw288 | TM7MB1-1 | A:A | A:A | FRANCE |
| lw149 | HYBRIDE 56 VILMORIN | A:A | A:A | FRANCE |
| lw031 | ARGENT | A:A | A:A | GREAT BRITAIN |
| DPG_546 | Hereward | A:A | A:A | GREAT BRITAIN |
| lw128 | G66257 | A:A | A:A | GREECE |
| lw138 | GODOLLOI 15 | A:A | A:A | HUNGARIA |
| lw136 | GK SZOKE | A:A | A:A | HUNGARIA |
| lw303 | VOLT | A:A | A:A | HUNGARIA |
| lw155 | JASZsAgI TF | A:A | A:A | HUNGARIA |
| DPG_611 | KALYANSONA | A:A | A:A | INDIA |
| DPG_574 | INDIA 107 | A:A | A:A | INDIA |
| DPG_581 | INDIA 231 | A:A | A:A | INDIA |
| DPG_584 | INDIA 325 | A:A | A:A | INDIA |

| | | | | |
|---|---|---|---|---|
| DPG_586 | Indian 6 | A:A | A:A | INDIA |
| lw203 | NAPHAL | A:A | A:A | INDIA |
| DPG_576 | INDIA 120 | A:A | A:A | INDIA |
| DPG_593 | IRAN 68 | A:A | A:A | IRAN |
| DPG_111 | AMC 157 | A:A | A:A | IRAN |
| DPG_598 | IRAQ 47 | A:A | A:A | IRAQ |
| DPG_539 | H742A | A:A | A:A | ISRAEL |
| DPG_726 | N67M2 | A:A | A:A | ISRAEL |
| DPG_926 | VOLCANI-BPA | A:A | A:A | ISRAEL |
| DPG_725 | N46 | A:A | A:A | ISRAEL |
| DPG_668 | M708//G25/N163 | A:A | A:A | ISRAEL |
| lw030 | Ardito | A:A | A:A | ITALY |
| DPG_736 | Norin 10 | A:A | A:A | JAPAN |
| DPG_739 | NORIN 64 | A:A | A:A | JAPAN |
| DPG_737 | NORIN 29 | A:A | A:A | JAPAN |
| DPG_449 | FUKUHOKOMUGI | A:A | A:A | JAPAN |
| lw148 | HOROSHIRI KOMUGI | A:A | A:A | JAPAN |
| DPG_738 | NORIN 60 | A:A | A:A | JAPAN |
| lw281 | NISHIKAZE KOMUGI | A:A | A:A | JAPAN |
| lw178 | MACHA | A:A | A:A | JAPAN |
| DPG_334 | Cul2 | A:A | A:A | KAZAKHSTAN |
| DPG_337 | Cul5 | A:A | A:A | KAZAKHSTAN |
| DPG_333 | Cul10 | A:A | A:A | KAZAKHSTAN |
| lw163 | KENYA 350 | A:A | A:A | KENYA |
| DPG_711 | Mexique 58 | A:A | A:A | MEXICO |
| DPG_803 | Pitic 62 | A:A | A:A | MEXICO |
| DPG_588 | INIA 66 | A:A | A:A | MEXICO |
| DPG_710 | MEXICO-120 | A:A | A:A | MEXICO |
| DPG_208 | CIANO 79 | A:A | A:A | MEXICO |
| DPG_307 | CLEOPATRA | A:A | A:A | MEXICO |
| DPG_463 | GENARO F81 | A:A | A:A | MEXICO |
| DPG_752 | OPATA85 | A:A | A:A | MEXICO |
| DPG_436 | Fang 60 | A:A | A:A | MEXICO |
| DPG_180 | BURRION | A:A | A:A | MEXICO |
| DPG_761 | PASTOR | A:A | A:A | MEXICO |
| DPG_154 | BERKUT | A:A | A:A | MEXICO |
| DPG_890 | Tacupeto 2001 | A:A | A:A | MEXICO |
| DPG_927 | VOROBEY | A:A | A:A | MEXICO |
| DPG_220 | cid279807sid86 | A:A | A:A | MEXICO |
| DPG_176 | 7HRWSN108 | A:A | A:A | MEXICO |
| DPG_213 | cid183961sid5 | A:A | A:A | MEXICO |
| DPG_214 | cid192646sid0 | A:A | A:A | MEXICO |
| DPG_216 | cid20569sid929 | A:A | A:A | MEXICO |
| DPG_217 | cid20570sid926 | A:A | A:A | MEXICO |
| DPG_219 | cid260142sid43 | A:A | A:A | MEXICO |
| DPG_222 | cid279826sid63 | A:A | A:A | MEXICO |
| DPG_223 | cid306547sid131 | A:A | A:A | MEXICO |
| DPG_225 | cid325362sid63 | A:A | A:A | MEXICO |
| DPG_226 | cid334938sid1119 | A:A | A:A | MEXICO |
| DPG_227 | cid335446sid92 | A:A | A:A | MEXICO |
| DPG_228 | cid373279sid61 | A:A | A:A | MEXICO |
| DPG_230 | cid385184sid26 | A:A | A:A | MEXICO |

| | | | | |
|---|---|---|---|---|
| DPG_231 | cid388327sid117 | A:A | A:A | MEXICO |
| DPG_233 | cid388408sid65 | A:A | A:A | MEXICO |
| DPG_234 | cid388412sid46 | A:A | A:A | MEXICO |
| DPG_235 | cid388436sid70 | A:A | A:A | MEXICO |
| DPG_236 | cid388444sid89 | A:A | A:A | MEXICO |
| DPG_237 | cid394062sid22 | A:A | A:A | MEXICO |
| DPG_238 | cid394070sid114 | A:A | A:A | MEXICO |
| DPG_239 | cid394087sid30 | A:A | A:A | MEXICO |
| DPG_240 | cid394715sid160 | A:A | A:A | MEXICO |
| DPG_242 | cid399013sid209 | A:A | A:A | MEXICO |
| DPG_243 | cid399062sid49 | A:A | A:A | MEXICO |
| DPG_244 | cid420707sid98 | A:A | A:A | MEXICO |
| DPG_245 | cid423295sid45 | A:A | A:A | MEXICO |
| DPG_246 | cid425852sid48 | A:A | A:A | MEXICO |
| DPG_248 | cid427658sid33 | A:A | A:A | MEXICO |
| DPG_252 | cid428589sid93 | A:A | A:A | MEXICO |
| DPG_253 | cid428592sid124 | A:A | A:A | MEXICO |
| DPG_254 | cid428603sid31 | A:A | A:A | MEXICO |
| DPG_255 | cid431763sid639 | A:A | A:A | MEXICO |
| DPG_256 | cid431798sid387 | A:A | A:A | MEXICO |
| DPG_268 | cid460334sid63 | A:A | A:A | MEXICO |
| DPG_269 | cid460342sid40 | A:A | A:A | MEXICO |
| DPG_270 | cid460344sid158 | A:A | A:A | MEXICO |
| DPG_272 | cid460354sid105 | A:A | A:A | MEXICO |
| DPG_273 | cid460356sid168 | A:A | A:A | MEXICO |
| DPG_275 | cid460360sid38 | A:A | A:A | MEXICO |
| DPG_276 | cid460396sid87 | A:A | A:A | MEXICO |
| DPG_277 | cid463491sid196 | A:A | A:A | MEXICO |
| DPG_278 | cid463508sid200 | A:A | A:A | MEXICO |
| DPG_279 | cid472868sid38 | A:A | A:A | MEXICO |
| DPG_280 | cid472918sid54 | A:A | A:A | MEXICO |
| DPG_281 | cid472923sid71 | A:A | A:A | MEXICO |
| DPG_282 | cid472924sid40 | A:A | A:A | MEXICO |
| DPG_283 | cid472927sid53 | A:A | A:A | MEXICO |
| DPG_285 | cid473243sid61 | A:A | A:A | MEXICO |
| DPG_286 | cid473245sid56 | A:A | A:A | MEXICO |
| DPG_287 | cid473247sid51 | A:A | A:A | MEXICO |
| DPG_289 | cid473251sid35 | A:A | A:A | MEXICO |
| DPG_290 | cid473259sid69 | A:A | A:A | MEXICO |
| DPG_291 | cid473269sid30 | A:A | A:A | MEXICO |
| DPG_293 | cid480829sid70 | A:A | A:A | MEXICO |
| DPG_294 | cid486216sid209 | A:A | A:A | MEXICO |
| DPG_296 | cid69587sid58 | A:A | A:A | MEXICO |
| DPG_299 | cid7506sid110 | A:A | A:A | MEXICO |
| DPG_301 | cid82996sid44 | A:A | A:A | MEXICO |
| DPG_302 | cid8626sid502 | A:A | A:A | MEXICO |
| DPG_303 | cid91078sid341 | A:A | A:A | MEXICO |
| DPG_549 | HTWT_001 | A:A | A:A | MEXICO |
| DPG_551 | HTWT_003 | A:A | A:A | MEXICO |
| DPG_556 | HTWT_008 | A:A | A:A | MEXICO |
| DPG_557 | HTWT_009 | A:A | A:A | MEXICO |
| DPG_558 | HTWT_010 | A:A | A:A | MEXICO |

| | | | | |
|---|---|---|---|---|
| DPG_559 | HTWT_012 | A:A | A:A | MEXICO |
| DPG_560 | HTWT_013 | A:A | A:A | MEXICO |
| DPG_563 | HTWT_030 | A:A | A:A | MEXICO |
| DPG_564 | HTWT-011 | A:A | A:A | MEXICO |
| DPG_565 | HTWT-014 | A:A | A:A | MEXICO |
| DPG_567 | HTWT-031 | A:A | A:A | MEXICO |
| DPG_589 | INIA-19 | A:A | A:A | MEXICO |
| DPG_607 | JUPATECO 73 | A:A | A:A | MEXICO |
| DPG_1491 | VOROBEY-5 | A:A | A:A | MEXICO |
| DPG_1497 | cid183961sid5 | A:A | A:A | MEXICO |
| DPG_1498 | cid428603sid31 | A:A | A:A | MEXICO |
| DPG_1499 | cid394087sid30 | A:A | A:A | MEXICO |
| DPG_1500 | cid473259sid69 | A:A | A:A | MEXICO |
| DPG_1501 | cid473259sid70 | A:A | A:A | MEXICO |
| DPG_1502 | cid460354sid132 | A:A | A:A | MEXICO |
| DPG_257 | cid431801sid446 | A:A | A:A | MEXICO |
| DPG_260 | cid435275sid65 | A:A | A:A | MEXICO |
| DPG_261 | cid435303sid61 | A:A | A:A | MEXICO |
| DPG_262 | cid437242sid88 | A:A | A:A | MEXICO |
| DPG_266 | cid452470sid102 | A:A | A:A | MEXICO |
| DPG_858 | SIETE CERROS | A:A | A:A | MEXICO |
| lw079 | CHANATE | A:A | A:A | MEXICO |
| DPG_218 | cid257170sid21 | A:A | A:A | MEXICO |
| DPG_241 | cid394799sid20 | A:A | A:A | MEXICO |
| DPG_295 | cid6831sid90 | A:A | A:A | MEXICO |
| DPG_304 | cid94795sid11 | A:A | A:A | MEXICO |
| DPG_471 | GH 'S' | A:A | A:A | MEXICO |
| DPG_232 | cid388384sid40 | A:A | A:A | MEXICO |
| DPG_886 | SYNTHETIC W7984 | A:A | A:A | MEXICO |
| DPG_250 | cid427898sid67 | A:A | A:A | MEXICO |
| DPG_249 | cid427887sid0 | A:A | A:A | MEXICO |
| DPG_297 | cid72726sid102 | A:A | A:A | MEXICO |
| DPG_325 | CPI133859 | A:A | A:A | - |
| DPG_1013 | synthetic bread wheat with AE.SQUAR | A:A | A:A | - |
| DPG_561 | HTWT_015 | A:A | A:A | - |
| DPG_988 | synthetic bread wheat with AE.SQUAR | A:A | A:A | - |
| DPG_989 | synthetic bread wheat with AE.SQUAR | A:A | A:A | - |
| DPG_991 | synthetic bread wheat with AE.SQUAR | A:A | A:A | - |
| DPG_998 | synthetic bread wheat with AE.SQUAR | A:A | A:A | - |
| DPG_1002 | synthetic bread wheat with AE.SQUAR | A:A | A:A | - |
| DPG_1003 | synthetic bread wheat with AE.SQUAR | A:A | A:A | - |
| DPG_1005 | synthetic bread wheat with AE.SQUAR | A:A | A:A | - |
| DPG_1014 | synthetic bread wheat with AE.SQUAR | A:A | A:A | - |
| DPG_1015 | synthetic bread wheat with AE.SQUAR | A:A | A:A | - |
| DPG_1016 | synthetic bread wheat with AE.SQUAR | A:A | A:A | - |
| DPG_980 | synthetic bread wheat with AE.SQUAR | A:A | A:A | - |
| DPG_981 | synthetic bread wheat with AE.SQUAR | A:A | A:A | - |
| DPG_992 | synthetic bread wheat with AE.SQUAR | A:A | A:A | - |
| DPG_1495 | AUS-17838-4 | A:A | A:A | - |
| DPG_562 | HTWT_028 | A:A | A:A | - |
| DPG_566 | HTWT-029 | A:A | A:A | - |
| DPG_085 | 6HRWSN98 | A:A | A:A | - |

| | | | | |
|---|---|---|---|---|
| DPG_982 | synthetic bread wheat with AE.SQUAR | A:A | A:A | - |
| DPG_984 | synthetic bread wheat with AE.SQUAR | A:A | A:A | - |
| DPG_986 | synthetic bread wheat with AE.SQUAR | A:A | A:A | - |
| DPG_1008 | synthetic bread wheat with AE.SQUAR | A:A | A:A | - |
| DPG_323 | CPI133814 | A:A | A:A | - |
| DPG_1009 | synthetic bread wheat with AE.SQUAR | A:A | A:A | - |
| DPG_1011 | synthetic bread wheat with AE.SQUAR | A:A | A:A | - |
| DPG_987 | synthetic bread wheat with AE.SQUAR | A:A | A:A | - |
| DPG_1007 | synthetic bread wheat with AE.SQUAR | A:A | A:A | - |
| DPG_1010 | synthetic bread wheat with AE.SQUAR | A:A | A:A | - |
| DPG_995 | synthetic bread wheat with AE.SQUAR | A:A | A:A | - |
| DPG_999 | synthetic bread wheat with AE.SQUAR | A:A | A:A | - |
| DPG_1001 | synthetic bread wheat with AE.SQUAR | A:A | A:A | - |
| lw332 | HAMAC | A:A | A:A | NETHERLAND |
| DPG_808 | PRESTON | A:A | A:A | NEW-ZELAND |
| lw028 | ARAWA | A:A | A:A | NEW-ZELAND |
| lw263 | RONGOTEA | A:A | A:A | NZL |
| DPG_936 | W48B | A:A | A:A | PAKISTAN |
| DPG_929 | W100 | A:A | A:A | PAKISTAN |
| lw042 | BARANI 70 | A:A | A:A | PAKISTAN |
| lw301 | SANDOMIRKA | A:A | A:A | POLAND |
| DPG_717 | MOCHO DE ESPIGA BRANCA | A:A | A:A | PORTUGAL |
| lw228 | PRECOCE | A:A | A:A | PORTUGAL |
| DPG_745 | ODESSA EXPSTA21821 | A:A | A:A | PORTUGAL |
| lw216 | ODESSA EXPSTA20722 | A:A | A:A | PORTUGAL |
| DPG_401 | DOMINAT: 2793 | A:A | A:A | ROMANIA |
| lw221 | OULIANOWSKA | A:A | A:A | RUSSIA |
| lw121 | FERRUGINEUM | A:A | A:A | RUSSIA |
| lw140 | GRANIT | A:A | A:A | RUSSIA |
| DPG_587 | Indis | A:A | A:A | SOUTH-AFRICA |
| lw278 | WS-13 CARDENO 34/45 | A:A | A:A | SPAIN |
| lw072 | CANDEAL DE SORIA | A:A | A:A | SPAIN |
| DPG_701 | MEIRA | A:A | A:A | SPAIN |
| lw253 | TRIGO DE MONTE | A:A | A:A | SPAIN |
| lw242 | SOANDRES LARACHA | A:A | A:A | SPAIN |
| lw222 | P. DE BROLLON | A:A | A:A | SPAIN |
| lw052 | BLANC PRECOCE | A:A | A:A | SWITZERLAND |
| lw237 | ROUGE DE MARCHISSY | A:A | A:A | SWITZERLAND |
| lw280 | SANUNU | A:A | A:A | SYRIA |
| DPG_659 | LW348 | A:A | A:A | TAJIKISTAN |
| DPG_162 | BT2277 | A:A | A:A | TUNISIA |
| lw328 | Mirleben | A:A | A:A | UKRAINE |
| lw110 | DNEPROVSKAIA | A:A | A:A | UKRAINE |
| DPG_431 | ESTANZUELA DORADO | A:A | A:A | URUGUAY |
| DPG_853 | SENTINEL | A:A | A:A | USA |
| DPG_429 | ERNIE | A:A | A:A | USA |
| DPG_107 | ALSEN | A:A | A:A | USA |
| lw002 | 11IWSWSN14 | A:A | A:A | USA |
| DPG_392 | DAKOTA | A:A | A:A | USA |
| DPG_740 | Norin10/Brevor | A:A | A:A | USA |
| lw191 | MINTURK | A:A | A:A | USA |
| lw041 | Balkan | A:A | A:A | YUGOSLAVIA |

| | | | | |
|---|---|---|---|---|
| DPG_840 | S975-A4-A1 | A:A | A:A | ZIMBABWE |
| lw290 | LIMPOPO | A:A | A:A | ZIMBABWE |
| DPG_499 | H 1440 | G:G | - | AFGHANISTAN |
| DPG_511 | H 1540 | G:G | - | AFGHANISTAN |
| lw100 | DETENICKA CERVENA | G:G | - | CZECH REPUBLIC |
| lw157 | JUBING | G:G | - | NEPAL |
| DPG_534 | H 1703 | G:G | T:T | AFGHANISTAN |
| DPG_493 | H 1357 | G:G | T:T | AFGHANISTAN |
| DPG_487 | H 11 | G:G | T:T | AFGHANISTAN |
| DPG_518 | H 1601 | G:G | T:T | AFGHANISTAN |
| DPG_533 | H 1695 | G:G | T:T | AFGHANISTAN |
| DPG_1265 | H 1311 | G:G | T:T | AFGHANISTAN |
| DPG_496 | H 1398 | G:G | T:T | AFGHANISTAN |
| DPG_498 | H 1411 | G:G | T:T | AFGHANISTAN |
| DPG_500 | H 1445 | G:G | T:T | AFGHANISTAN |
| DPG_501 | H 1494 | G:G | T:T | AFGHANISTAN |
| DPG_502 | H 1496 | G:G | T:T | AFGHANISTAN |
| DPG_503 | H 1509 | G:G | T:T | AFGHANISTAN |
| DPG_504 | H 1511 | G:G | T:T | AFGHANISTAN |
| DPG_506 | H 1525 | G:G | T:T | AFGHANISTAN |
| DPG_507 | H 1527 | G:G | T:T | AFGHANISTAN |
| DPG_508 | H 1531 | G:G | T:T | AFGHANISTAN |
| DPG_509 | H 1534 | G:G | T:T | AFGHANISTAN |
| DPG_510 | H 1537 | G:G | T:T | AFGHANISTAN |
| DPG_514 | H 1550 | G:G | T:T | AFGHANISTAN |
| DPG_516 | H 1584 | G:G | T:T | AFGHANISTAN |
| DPG_520 | H 1621 | G:G | T:T | AFGHANISTAN |
| DPG_521 | H 1627 | G:G | T:T | AFGHANISTAN |
| DPG_522 | H 1632 | G:G | T:T | AFGHANISTAN |
| DPG_523 | H 1644 | G:G | T:T | AFGHANISTAN |
| DPG_524 | H 1647 | G:G | T:T | AFGHANISTAN |
| DPG_526 | H 1660 | G:G | T:T | AFGHANISTAN |
| DPG_528 | H 1673 | G:G | T:T | AFGHANISTAN |
| DPG_529 | H 1681A | G:G | T:T | AFGHANISTAN |
| DPG_531 | H 1686B | G:G | T:T | AFGHANISTAN |
| DPG_532 | H 1691 | G:G | T:T | AFGHANISTAN |
| DPG_536 | H 344 | G:G | T:T | AFGHANISTAN |
| DPG_1264 | H 1305 | G:G | T:T | AFGHANISTAN |
| DPG_1268 | H 1359 | G:G | T:T | AFGHANISTAN |
| DPG_1269 | H 1369 | G:G | T:T | AFGHANISTAN |
| DPG_1271 | H 1399 | G:G | T:T | AFGHANISTAN |
| lw016 | A4 | G:G | T:T | AFGHANISTAN |
| DPG_151 | BAHATANE 87 | G:G | T:T | ALGERIA |
| DPG_175 | BUCK_ATLANTICO | G:G | T:T | ARGENTINA |
| DPG_423 | EGYPCIO208 | G:G | T:T | ARGENTINA |
| DPG_652 | LW333 | G:G | T:T | ARMENIA |
| DPG_405 | Du Toits | G:G | T:T | AUSTRALIA |
| DPG_568 | Hudsons Early Purple Straw | G:G | T:T | AUSTRALIA |
| DPG_942 | Ward's Prolific | G:G | T:T | AUSTRALIA |
| DPG_437 | Federation | G:G | T:T | AUSTRALIA |
| DPG_441 | Florence | G:G | T:T | AUSTRALIA |
| lw086 | COLLEGE ECLIPSE | G:G | T:T | AUSTRALIA |

| | | | | |
|---|---|---|---|---|
| DPG_343 | Currawa | G:G | T:T | AUSTRALIA |
| DPG_623 | KINGSWHITE | G:G | T:T | AUSTRALIA |
| DPG_941 | Waratah | G:G | T:T | AUSTRALIA |
| DPG_472 | Ghurka | G:G | T:T | AUSTRALIA |
| DPG_828 | Ranee | G:G | T:T | AUSTRALIA |
| DPG_445 | Free Gallipoli | G:G | T:T | AUSTRALIA |
| DPG_407 | Dundee | G:G | T:T | AUSTRALIA |
| DPG_179 | Bungulla | G:G | T:T | AUSTRALIA |
| DPG_161 | Bencubbin | G:G | T:T | AUSTRALIA |
| DPG_477 | GLUYAS EARLY | G:G | T:T | AUSTRALIA |
| DPG_821 | Quadrat | G:G | T:T | AUSTRALIA |
| DPG_802 | Pinnacle | G:G | T:T | AUSTRALIA |
| DPG_476 | Glenwari | G:G | T:T | AUSTRALIA |
| DPG_547 | Heron | G:G | T:T | AUSTRALIA |
| DPG_434 | Falcon | G:G | T:T | AUSTRALIA |
| DPG_461 | Gatcher | G:G | T:T | AUSTRALIA |
| DPG_312 | Combination III | G:G | T:T | AUSTRALIA |
| DPG_540 | HALBERD | G:G | T:T | AUSTRALIA |
| DPG_422 | EGRET | G:G | T:T | AUSTRALIA |
| DPG_625 | KITE | G:G | T:T | AUSTRALIA |
| DPG_619 | Kewell | G:G | T:T | AUSTRALIA |
| DPG_905 | TINCURRIN | G:G | T:T | AUSTRALIA |
| DPG_943 | Warrigal | G:G | T:T | AUSTRALIA |
| DPG_143 | AVOCET | G:G | T:T | AUSTRALIA |
| DPG_715 | MILLEWA | G:G | T:T | AUSTRALIA |
| DPG_486 | GUTHA | G:G | T:T | AUSTRALIA |
| DPG_541 | Harrier | G:G | T:T | AUSTRALIA |
| DPG_391 | DAGGER | G:G | T:T | AUSTRALIA |
| DPG_867 | SPEAR | G:G | T:T | AUSTRALIA |
| DPG_878 | Sunstar | G:G | T:T | AUSTRALIA |
| DPG_876 | SUNELG | G:G | T:T | AUSTRALIA |
| DPG_919 | Vasco | G:G | T:T | AUSTRALIA |
| DPG_928 | VULCAN | G:G | T:T | AUSTRALIA |
| lw087 | COMET | G:G | T:T | AUSTRALIA |
| DPG_719 | MOLINEUX | G:G | T:T | AUSTRALIA |
| DPG_908 | TRIDENT | G:G | T:T | AUSTRALIA |
| DPG_344 | CURRAWONG | G:G | T:T | AUSTRALIA |
| DPG_967 | YIPTI | G:G | T:T | AUSTRALIA |
| DPG_444 | FRAME | G:G | T:T | AUSTRALIA |
| DPG_902 | TENNANT | G:G | T:T | AUSTRALIA |
| DPG_105 | AJANA | G:G | T:T | AUSTRALIA |
| DPG_633 | KUKRI | G:G | T:T | AUSTRALIA |
| DPG_417 | EGA BONNIE ROCK | G:G | T:T | AUSTRALIA |
| DPG_419 | EGA JITARNING | G:G | T:T | AUSTRALIA |
| DPG_415 | EGA 2248 | G:G | T:T | AUSTRALIA |
| DPG_475 | GLADIUS | G:G | T:T | AUSTRALIA |
| DPG_673 | MAGENTA | G:G | T:T | AUSTRALIA |
| DPG_766 | PEAKE | G:G | T:T | AUSTRALIA |
| DPG_318 | CORRELL | G:G | T:T | AUSTRALIA |
| DPG_430 | ESPADA | G:G | T:T | AUSTRALIA |
| DPG_669 | MACE | G:G | T:T | AUSTRALIA |
| DPG_622 | KING ROCK | G:G | T:T | AUSTRALIA |

| | | | | |
|---|---|---|---|---|
| lw143 | GULAR | G:G | T:T | AUSTRALIA |
| DPG_138 | AUS1490 | G:G | T:T | AUSTRALIA |
| DPG_636 | KUNJIN | G:G | T:T | AUSTRALIA |
| DPG_817 | Purple Straw AUS885 | G:G | T:T | AUSTRALIA |
| DPG_957 | WW2449 | G:G | T:T | AUSTRALIA |
| lw161 | KATYIL | G:G | T:T | AUSTRALIA |
| DPG_871 | SUN325B | G:G | T:T | AUSTRALIA |
| lw038 | AUSTRO_BANKUT | G:G | T:T | AUSTRIA |
| lw294 | NON_PLUS_EXTRA | G:G | T:T | AUSTRIA |
| lw018 | ADMONTER | G:G | T:T | AUSTRIA |
| lw070 | 114/62 | G:G | T:T | AUSTRIA |
| lw036 | ATUT II | G:G | T:T | AUSTRIA |
| lw371 | Landrace | G:G | T:T | AZERBAIJAN |
| lw372 | Arazbugdasi | G:G | T:T | AZERBAIJAN |
| lw271 | ZANDA | G:G | T:T | BELGIUM |
| lw158 | JUFY II | G:G | T:T | BELGIUM |
| DPG_191 | CARAZINHO | G:G | T:T | BRAZIL |
| DPG_446 | FRONTANA 3671 | G:G | T:T | BRAZIL |
| DPG_674 | Maringa (Rht1) | G:G | T:T | BRAZIL |
| lw015 | 664-258-18 | G:G | T:T | BULGARIA |
| lw007 | 2838-39 | G:G | T:T | BULGARIA |
| lw160 | K1898-9/L200-6 | G:G | T:T | BULGARIA |
| lw232 | REDMAN | G:G | T:T | CANADA |
| lw310 | AGATHA | G:G | T:T | CANADA |
| lw073 | CANUCK | G:G | T:T | CANADA |
| DPG_614 | KATEPWA | G:G | T:T | CANADA |
| DPG_833 | Roblin | G:G | T:T | CANADA |
| DPG_092 | AC DOMAIN | G:G | T:T | CANADA |
| DPG_091 | AC BARRIE | G:G | T:T | CANADA |
| lw137 | GLENLEA | G:G | T:T | CANADA |
| lw272 | Coronation | G:G | T:T | CANADA |
| lw273 | KITCHENER | G:G | T:T | CANADA |
| lw358 | Krasnaya | G:G | T:T | CANADA |
| DPG_729 | NANKING NO25 | G:G | T:T | CHINA |
| DPG_206 | CHUAN_MAI_18 (CM18) | G:G | T:T | CHINA |
| DPG_615 | KE-HAN-NO8 | G:G | T:T | CHINA |
| DPG_204 | **Chinese Spring** | G:G | T:T | CHINA |
| lw206 | NONG DA NO141 | G:G | T:T | CHINA |
| DPG_484 | Guisuiskaya Syao-bai-mai | G:G | T:T | CHINA |
| DPG_634 | Kuldzhinskaya | G:G | T:T | CHINA |
| lw111 | DONG-FANG-HONG-NO3 | G:G | T:T | CHINA |
| lw021 | AIFENG NO4 | G:G | T:T | CHINA |
| DPG_962 | XIAOYAN54 | G:G | T:T | CHINA |
| lw239 | SENTUNG | G:G | T:T | CHINA |
| lw341 | Landrace | G:G | T:T | CHINA |
| lw342 | Landrace | G:G | T:T | CHINA |
| lw344 | R 1 | G:G | T:T | CHINA |
| DPG_913 | U-Man-Syao-Mai | G:G | T:T | CHINA |
| lw025 | ANDES 56 | G:G | T:T | COLOMBIA |
| lw257 | US(62)P66 | G:G | T:T | COLOMBIA |
| lw337 | Landrace | G:G | T:T | ETHIOPIA |
| lw293 | Hopea | G:G | T:T | FINLAND |

| | | | | |
|---|---|---|---|---|
| DPG_604 | JO3045 | G:G | T:T | FINLAND |
| DPG_912 | Ulla | G:G | T:T | FINLAND |
| lw050 | BERZATACA | G:G | T:T | FINLAND |
| lw175 | LONTOI | G:G | T:T | FINLAND |
| lw246 | TAHTI | G:G | T:T | FINLAND |
| lw093 | COURTOT | G:G | T:T | FRANCE |
| DPG_201 | CHAMPTAL | G:G | T:T | FRANCE |
| lw091 | CORSODOR | G:G | T:T | FRANCE |
| lw098 | DAVIDOC | G:G | T:T | FRANCE |
| lw134 | GH126 | G:G | T:T | FRANCE |
| lw004 | DI185 | G:G | T:T | FRANCE |
| lw054 | BLE D'OR | G:G | T:T | FRANCE |
| lw210 | NOUGAT | G:G | T:T | FRANCE |
| lw382 | DI6404-19-15 | G:G | T:T | FRANCE |
| lw011 | DI50-12 | G:G | T:T | FRANCE |
| lw114 | E108 | G:G | T:T | FRANCE |
| DPG_891 | TAFERSTAT | G:G | T:T | FRANCE |
| DPG_676 | MARS DE SUEDE ROUGE BARBU | G:G | T:T | FRANCE |
| lw103 | DI7003-1-12 | G:G | T:T | FRANCE |
| lw106 | DI7210-15-11 | G:G | T:T | FRANCE |
| lw129 | G7 | G:G | T:T | FRANCE |
| lw003 | DI15 | G:G | T:T | FRANCE |
| lw006 | DI276 | G:G | T:T | FRANCE |
| lw008 | DI330 | G:G | T:T | FRANCE |
| lw043 | BARBU DU FINISTERE | G:G | T:T | FRANCE |
| lw044 | BARBU DU TRONCHET | G:G | T:T | FRANCE |
| lw056 | BLE DE HAIE | G:G | T:T | FRANCE |
| lw076 | CF3003-2-7-4-4-3 | G:G | T:T | FRANCE |
| lw094 | CP4 | G:G | T:T | FRANCE |
| lw077 | CF4563-1-5-3-2-5 | G:G | T:T | FRANCE |
| lw169 | KRELOF 3 | G:G | T:T | FRANCE |
| lw195 | MOTTIN | G:G | T:T | FRANCE |
| lw234 | RICHELLE | G:G | T:T | FRANCE |
| lw236 | ROUGE-D'ALTKIRCH | G:G | T:T | FRANCE |
| lw250 | TRAMI~PUY DE DOME | G:G | T:T | FRANCE |
| lw264 | BLANC DE CHENAN | G:G | T:T | FRANCE |
| lw265 | BLE DANOIS | G:G | T:T | FRANCE |
| lw270 | BLE BARBU DE MUROL | G:G | T:T | FRANCE |
| lw292 | GENESIS | G:G | T:T | FRANCE |
| lw306 | TALDOR | G:G | T:T | FRANCE |
| lw321 | BLE DE REDON BLANC BARBU 1 1 | G:G | T:T | FRANCE |
| lw325 | BLE DE REDON ST-JUST Q21 | G:G | T:T | FRANCE |
| lw385 | SPONSOR | G:G | T:T | FRANCE |
| lw386 | TALISMAN | G:G | T:T | FRANCE |
| lw268 | INSTITUT 1802 | G:G | T:T | FRANCE |
| lw350 | Landrace | G:G | T:T | GEORGIA |
| lw352 | Landrace | G:G | T:T | GEORGIA |
| lw364 | Dolis Puri | G:G | T:T | GEORGIA |
| lw243 | Strubes Dickkopf | G:G | T:T | GERMANY |
| lw219 | Opal | G:G | T:T | GERMANY |
| lw032 | Arkas | G:G | T:T | GERMANY |
| lw300 | FRUH-WEIZEN | G:G | T:T | GERMANY |

| | | | | |
|---|---|---|---|---|
| lw019 | Adular | G:G | T:T | GERMANY |
| lw026 | APEX | G:G | T:T | GREAT BRITAIN |
| lw182 | MASTER | G:G | T:T | GREAT BRITAIN |
| DPG_451 | G72300 | G:G | T:T | GREECE |
| lw047 | BELLIEI 590 | G:G | T:T | HUNGARIA |
| lw183 | MATRADERECSKEITF | G:G | T:T | HUNGARIA |
| lw196 | MV MA | G:G | T:T | HUNGARIA |
| DPG_741 | NP120 | G:G | T:T | INDIA |
| DPG_870 | Sujata | G:G | T:T | INDIA |
| DPG_765 | PBW343 | G:G | T:T | INDIA |
| DPG_572 | INDIA 105 | G:G | T:T | INDIA |
| DPG_575 | INDIA 108 | G:G | T:T | INDIA |
| DPG_580 | INDIA 223 | G:G | T:T | INDIA |
| DPG_582 | INDIA 322 | G:G | T:T | INDIA |
| DPG_583 | INDIA 323 | G:G | T:T | INDIA |
| DPG_585 | INDIA 39 | G:G | T:T | INDIA |
| DPG_814 | Punjab Tipe 15 | G:G | T:T | INDIA |
| DPG_815 | Punjab Tipe 18 | G:G | T:T | INDIA |
| DPG_816 | Punjab Tipe 24 | G:G | T:T | INDIA |
| DPG_930 | W195 | G:G | T:T | INDIA |
| DPG_573 | INDIA 106 | G:G | T:T | INDIA |
| DPG_775 | PERSIA 21 | G:G | T:T | IRAN |
| DPG_772 | PERSIA 111 | G:G | T:T | IRAN |
| DPG_778 | PERSIA 4 | G:G | T:T | IRAN |
| DPG_110 | AMC 142 | G:G | T:T | IRAN |
| DPG_771 | PERSIA 11 | G:G | T:T | IRAN |
| DPG_774 | PERSIA 17 | G:G | T:T | IRAN |
| DPG_776 | PERSIA 28 | G:G | T:T | IRAN |
| DPG_777 | PERSIA 29 | G:G | T:T | IRAN |
| DPG_779 | PERSIA 7 | G:G | T:T | IRAN |
| DPG_595 | IRAQ 40 | G:G | T:T | IRAQ |
| DPG_596 | IRAQ 42 | G:G | T:T | IRAQ |
| DPG_597 | Iraq 43 | G:G | T:T | IRAQ |
| DPG_1493 | AUS-4885-6 | G:G | T:T | IRAQ |
| lw220 | Orlandi | G:G | T:T | ITALY |
| DPG_479 | Granarolo | G:G | T:T | ITALY |
| DPG_698 | Mec | G:G | T:T | ITALY |
| DPG_767 | Pegaso | G:G | T:T | ITALY |
| DPG_705 | Mentana | G:G | T:T | ITALY |
| lw040 | BAIONETTE I | G:G | T:T | ITALY |
| DPG_807 | PRECOCE-DU-JAPON | G:G | T:T | JAPAN |
| lw022 | AKADARUMA | G:G | T:T | JAPAN |
| lw035 | ATAKEDA KOMUGI | G:G | T:T | JAPAN |
| lw241 | SIRAZAIAIBARIGI 2 | G:G | T:T | JAPAN |
| lw212 | NYU BAY | G:G | T:T | JAPAN |
| DPG_332 | Cul1 | G:G | T:T | KAZAKHSTAN |
| DPG_340 | Cul8 | G:G | T:T | KAZAKHSTAN |
| DPG_341 | Cul9 | G:G | T:T | KAZAKHSTAN |
| lw020 | AFRICA MAYO | G:G | T:T | KENYA |
| lw240 | SEU SEUN 27 | G:G | T:T | KOREA |
| lw336 | Landrace | G:G | T:T | MAROCCO |
| DPG_865 | Sonora 64 | G:G | T:T | MEXICO |

| | | | | |
|---|---|---|---|---|
| DPG_805 | Potam 70 | G:G | T:T | MEXICO |
| DPG_426 | EMU'S' | G:G | T:T | MEXICO |
| DPG_251 | cid428587sid159 | G:G | T:T | MEXICO |
| DPG_298 | cid7401sid122 | G:G | T:T | MEXICO |
| DPG_209 | cid181178sid3 | G:G | T:T | MEXICO |
| DPG_210 | cid183298sid1 | G:G | T:T | MEXICO |
| DPG_211 | cid183300sid3 | G:G | T:T | MEXICO |
| DPG_212 | cid183347sid7 | G:G | T:T | MEXICO |
| DPG_215 | cid195095sid3 | G:G | T:T | MEXICO |
| DPG_221 | cid279818sid83 | G:G | T:T | MEXICO |
| DPG_300 | cid8090sid414 | G:G | T:T | MEXICO |
| DPG_720 | Morocco 426 | G:G | T:T | MEXICO |
| DPG_712 | Mexique 9 | G:G | T:T | MEXICO |
| DPG_330 | CS(Hope3B) | G:G | T:T | - |
| DPG_137 | AUS1408 | G:G | T:T | - |
| DPG_139 | AUS19123 (ASIA MINOR 7 1)=AMC 71 | G:G | T:T | - |
| DPG_796 | PI624933-1 | G:G | T:T | - |
| DPG_797 | PI625123-3 | G:G | T:T | - |
| DPG_799 | PI626580-4 | G:G | T:T | - |
| DPG_1494 | AUS-28080-4 | G:G | T:T | - |
| DPG_1496 | AUS-14828-6 | G:G | T:T | - |
| DPG_1012 | synthetic bread wheat with AE.SQUAR | G:G | T:T | - |
| DPG_922 | Vigour 18 | G:G | T:T | - |
| DPG_798 | PI625983-1 | G:G | T:T | - |
| lw099 | DAYKU | G:G | T:T | NEPAL |
| lw142 | GUDEL | G:G | T:T | NEPAL |
| lw147 | HONGGAOAN | G:G | T:T | NEPAL |
| lw048 | BENCHUNG | G:G | T:T | NEPAL |
| lw069 | BUNG EPI BLANC | G:G | T:T | NEPAL |
| lw082 | CHITLANG | G:G | T:T | NEPAL |
| lw084 | CHYAKSILA EPI NON VELU | G:G | T:T | NEPAL |
| lw085 | CHYAMTANG | G:G | T:T | NEPAL |
| lw159 | JUNBESI | G:G | T:T | NEPAL |
| lw170 | KULUNG | G:G | T:T | NEPAL |
| lw194 | MONJO | G:G | T:T | NEPAL |
| lw199 | NABU EPI BLANC | G:G | T:T | NEPAL |
| lw204 | NEPAL 84 | G:G | T:T | NEPAL |
| lw205 | NEPAL 89 | G:G | T:T | NEPAL |
| lw225 | PEYKRU INERME NON VELU | G:G | T:T | NEPAL |
| lw235 | RITAK | G:G | T:T | NEPAL |
| DPG_728 | NACHIPUNDO | G:G | T:T | NEPAL |
| lw132 | GAU | G:G | T:T | NEPAL |
| lw200 | NACHIPUNDO | G:G | T:T | NEPAL |
| DPG_812 | PROMESSA | G:G | T:T | NETHERLAND |
| lw013 | 60293 | G:G | T:T | NETHERLAND |
| DPG_702 | MELCHIOR | G:G | T:T | NETHERLAND |
| lw213 | NZ(81)P43 | G:G | T:T | NEW ZELAND |
| lw282 | HOUMANA | G:G | T:T | NIGERIA |
| DPG_666 | LYALLPUR-73 | G:G | T:T | PAKISTAN |
| lw351 | Soor Ghanum | G:G | T:T | PAKISTAN |
| DPG_081 | 117-var.12/564 | G:G | T:T | PAKISTAN |
| DPG_577 | INDIA 134 | G:G | T:T | PAKISTAN |

| | | | | |
|---|---|---|---|---|
| DPG_931 | W21A | G:G | T:T | PAKISTAN |
| DPG_932 | W21B | G:G | T:T | PAKISTAN |
| DPG_937 | W61 | G:G | T:T | PAKISTAN |
| DPG_938 | W93 | G:G | T:T | PAKISTAN |
| DPG_939 | W96 | G:G | T:T | PAKISTAN |
| lw363 | Spin, 121-var.12/536 | G:G | T:T | PAKISTAN |
| DPG_904 | Thori, 212-var.8/1 | G:G | T:T | PAKISTAN |
| DPG_758 | Palestinskaya | G:G | T:T | PALESTINE |
| lw368 | Landrace | G:G | T:T | PALESTINE |
| lw108 | DIANA | G:G | T:T | POLAND |
| lw060 | BLONDYNKA | G:G | T:T | POLAND |
| lw095 | D130-63 | G:G | T:T | POLAND |
| lw173 | LESZYNSKA WCZESNA | G:G | T:T | POLAND |
| lw184 | MEDRA/AURORA//HOHENTURM/DINA | G:G | T:T | POLAND |
| lw074 | CENAD 512 | G:G | T:T | ROMANIA |
| lw361 | Sary-bugda, Gulgeri | G:G | T:T | RUSSIA |
| DPG_393 | Dalnevostochnaya 10 | G:G | T:T | RUSSIA |
| lw343 | Sarrubra | G:G | T:T | RUSSIA |
| lw353 | Landrace | G:G | T:T | RUSSIA |
| lw365 | Tau-bugda | G:G | T:T | RUSSIA |
| lw366 | Albidum 12 | G:G | T:T | RUSSIA |
| lw096 | DAERAAD | G:G | T:T | SOUTH-AFRICA |
| lw071 | CANDEAL DE AREVALO | G:G | T:T | SPAIN |
| lw201 | NANG NANO 140 | G:G | T:T | SPAIN |
| lw115 | EBRO | G:G | T:T | SPAIN |
| lw145 | H93-70 | G:G | T:T | SPAIN |
| lw277 | NAVARRO150 | G:G | T:T | SPAIN |
| lw051 | BIRGITTA | G:G | T:T | SWEDEN |
| lw167 | KOLBEN 3 | G:G | T:T | SWEDEN |
| lw001 | CH01193 | G:G | T:T | SWITZERLAND |
| lw014 | CH62022 | G:G | T:T | SWITZERLAND |
| lw078 | CH73052 | G:G | T:T | SWITZERLAND |
| lw279 | NEELKANT | G:G | T:T | SYRIA |
| lw214 | ODESSA EXP.STA.17413 | G:G | T:T | SYRIA |
| lw152 | ICARDA079 | G:G | T:T | SYRIA |
| DPG_660 | LW349 | G:G | T:T | TAJIKISTAN |
| lw347 | Landrace | G:G | T:T | TAJIKISTAN |
| DPG_174 | BT2281 | G:G | T:T | TUNISIA |
| lw063 | BT2277 | G:G | T:T | TUNISIA |
| lw065 | BT2296 | G:G | T:T | TUNISIA |
| DPG_760 | PAMUKALE | G:G | T:T | TURKEY |
| lw334 | Landrace | G:G | T:T | TURKEY |
| lw335 | Landrace | G:G | T:T | TURKEY |
| DPG_316 | COPPADRA | G:G | T:T | TURKEY |
| DPG_975 | ZILVE | G:G | T:T | TURKEY |
| lw360 | Landrace | G:G | T:T | TURKEY |
| lw218 | ODESSKAIA 16 | G:G | T:T | UKRAINE |
| lw329 | FANTASIYA-ODESSKAYA | G:G | T:T | UKRAINE |
| lw359 | Stepnyachka | G:G | T:T | UKRAINE |
| DPG_868 | Spica | G:G | T:T | USA |
| lw125 | FRONTHATCH | G:G | T:T | USA |
| DPG_894 | TAM 107 | G:G | T:T | USA |

| | | | | |
|---|---|---|---|---|
| lw256 | US(59)34 | G:G | T:T | USA |
| lw123 | FLINT | G:G | T:T | USA |
| lw174 | Little Club | G:G | T:T | USA |
| lw088 | Compton | G:G | T:T | USA |
| lw049 | Benni | G:G | T:T | USA |
| lw061 | Blueboy | G:G | T:T | USA |
| lw255 | Tyler | G:G | T:T | USA |
| lw312 | CENTURK | G:G | T:T | USA |
| lw081 | CHIRKUNG | - | T:T | NEPAL |

| Year of release | Pedigree |
|---|---|
| | LV-FRA[39][1735] |
| 1983 | HPG-522-66/MARIS-DOVE[306][759][851][1343][1413][1790] |
| | - |
| | (S)LV-TIBET[39][2535] |
| 1989 | - |
| | - |
| | - |
| | - |
| | - |
| | - |
| | - |
| | - |
| | - |
| | - |
| | - |
| | - |
| | - |
| | - |
| | - |
| 1895 | (S)WARDS-PROLIFIC[56][39][1138][1451]; (S)BUDDS-EARLY[1322] |
| 1916 | FANCY/CROSSBRED-53[56][39][1138][1451] |
| 1951 | FORD/DUNDEE[39][1138] |
| 1955 | DUNDEE/KENYA-745-C-6042//BOBIN*2/GAZA[39][1138]; DUNDEE/GABO[56] |
| 1960 | LAWRENCE/GABO[56][39][1138] |
| 1960 | KENYA-117-A/2*GABO//MENTANA/6*GABO[56][39]; GABO/3/GABO*5/MENTANA//GABO |
| 1963 | MAYO-48/URUGUAY-1084//ORFED/3/4*DIRK-48[1281]; MAYO/4*DIRK-48[39] |
| 1963 | FESTIVAL/URUGUAY-C-10837[39][851][1138]; FESTIVAL/(URY)CI-10837[1335] |
| 1966 | SUNSTATE/QH-71-6//YITPI[4113] |
| 1973 | KHAPSTEIN*3/GABO//GAMENYA[39]; GAMENYA//3*GABO/KHAPSTEIN[113]; GAMENYA// |
| 1975 | LERMA-ROJO-64/SONORA-64-A//(SIB)TIMGALEN[113][1322] |
| 1977 | TIMGALEN/(SIB)CONDOR//CONDOR[760][1322]; TIMGALEN/2*CONDOR[000] |
| 1977 | SONORA-64/P-4160-3//3*TIMGALEN[760][851][113] |
| 1979 | PWTH/(SIB)CONDOR//2*CONDOR[113][760][851] |
| 1981 | WW-15/RAVEN[113][626][851] |
| 1982 | CIANO-67//SPICA/(AMBER-MUTANT)SONORA-64[113][176][760][851] |
| 1982 | KALYANSONA/OLYMPIC[176][760][113][2854] |
| 1982 | VICAM-71//CIANO-67(SIB)/SIETE-CERROS-66/3/KALYANSONA/BLUEBIRD[176][114][144][ |
| 1982 | BOKAL/SIETE-CERROS-66[113][851][626][760] |
| 1982 | CIANO-67/GAMENYA[626][760][113] |
| 1982 | PWTH/(SIB)CONDOR//2*CONDOR[113][760][851] |
| 1983 | FLINDERS(SIB)/2*COOK[176][976][851] |
| 1984 | SONORA-64/KNOTT*2//II-8156(DX-3-134)/3/OLYMPIC[113][851][976]; DS-3-134/OLYMP |
| 1984 | PAVON-76/CONDOR[2854][3756] |
| 1984 | AUS-10894/4*CONDOR[760][851][976] |
| 1984 | (S)CONDOR[760][976]; WW-80/2*WW-15[1281] |
| 1985 | FARRO-LUNGO/HERON//2*CONDOR/3/(SIB)QUARRION[760][1322]; FARRO-LUNGO/HERO |
| 1985 | WREN,MEX//CIANO-67(SIB)/NOROESTE-66/3/ZAMBEZI |
| 1985 | MEC-3/2*GABO(RAC-177)//MADDEN[626][113] |
| 1986 | BODALLIN(SIB)/(SIB)HYDEN[626][976][760][113] |
| 1986 | SUN-9-E-27*4/3-AG-14//WW-15/3/3*COOK[760][113] |
| 1988 | MEXICO-120/KODA//RAVEN/3/MENGAVI/SIETE-CERROS-66[708][760] |

| | |
|---|---|
| 1988 | CIANO-67/GALLO[626][708][760] |
| 1989 | 3-AG-3/4*CONDOR//COOK[723][760][1325] |
| 1989 | QT-3765/SUNCO[3596] |
| 1989 | BODALLIN//GAMENYA/INIA-66[626][760][113] |
| 1990 | RAC-177(Sr26)/UNICULM-492//RAC-311-S[1326] |
| 1990 | 3-AG-3/4*CONDOR//COOK[760][113] |
| 1990 | CANROCK-I/2*P-44//BANKS[755][760][113][707][1350]; CANROCK-I/2*CSP-44//BANKS[7 |
| 1990 | COOK*2/VPM-1//3*COOK[626][760][1322] |
| 1991 | BROCHIS(SIB)/BANKS[626][816][113] |
| 1992 | BOBWHITE/NACOZARI-76//VEERY/3/BLUEJAY/COCORAQUE-75[1281] |
| 1992 | CENTRIFEN/GAMENYA(F3)//GAMENYA/3/JACUP[113][4094] |
| 1993 | QT-2338/4*HARTOG[113][1345][816] |
| 1993 | HARTOG*4//COOK*5/VPM-1[3766] |
| 1994 | POTAM-70/4*COOK[113][1345] |
| 1994 | AROONA*3//(AUSEN-VII-95)TADORNA/INIA-66[1340][113][2854]; AROONA*3//AUSEN-VI |
| 1994 | COOK*2/VPM-1//3*COOK[1345][3766] |
| 1994 | BODALLIN//(74-W-07-47)ERADU/XBVT-233/3/(75-W-04-258)ATLAS-66/2*MADDEN[134 |
| 1995 | HARTOG*2/SUNECA[113]; HARTOG/SUNECA[2854] |
| 1996 | BOLSENA-1-CH(RAC-529)/77-W-660[3763]; RAC-529-911/77-W-660[3442] |
| 1997 | PITIC-62/FESTIGUAY//WARIMBA/3/MEXICO-120/QUADRAT/4/KLOKA/PITIC-62//BAYONET |
| 1997 | CO-1190-203/84-W-127-501[3767][4094] |
| 1997 | VICAM-71//CIANO-67/SIETE-CERROS-66/3/KALYANSONA/BLUEBIRD[1866]; VICAM-71/3/ |
| 1997 | CHINO/KULIN//REEVES[2854] |
| 1998 | KITE/2*OSPREY//HARTOG/OSPREY[2854] |
| 1998 | INIA-66/GAMUT//COOK/4/JUPATECO/3/LERMA-ROJO-64/SONORA-64-A//(SIB)TIMGALEN |
| 1998 | HARTOG/VEERY-5[1957][2854][3765] |
| 1998 | WARIQUAM//KLOKA/PITIC-62/3/WARIMEK/HALBERD/4/3-AG-3/AROONA[1866][2854]; |
| 1998 | KALYANSONA/BLUEBIRD//ANZA*3/WW-80/3/OLYMPIC*2/CIANO-67[2854]; B-1814//WV |
| 1998 | PAVON-76(SIB)/TM-56[2854][3764] |
| 1999 | LANCE*2/3/CONDOR*3//3-AG-14/CONDOR/4/4*TINCURRIN/5/TATIARA*3//COOK*5/VP |
| 1999 | VULKAN/3*JANZ[2854] |
| 1999 | M-3458/OSPREY[1349][3756] |
| 2001 | HARTOG*3/QUARRION[3775] |
| 2001 | JANZ*4/FIDEL[3758]; JANZ*4/FIDEL-FS-2[3758] |
| 2002 | PASTOR/3/ALTAR-84/AE.SQ(TR.TA)//OPATA-M-85[3692] |
| 2002 | M-3508/DOLLARBIRD[3777] |
| 2003 | VICAM-71/3*SUNECA//SUN-231-A[3596] |
| 2004 | BOBWHITE(SIB)/NARIS-HUNTSMAN//CRANBROOK/VICAM-71(83-Z-1288)/3/(IW-1266)PF |
| 2004 | SUNVALE/ROWAN[3596][3780] |
| 2004 | - |
| 2005 | ATTILA//ALTAR-84/ARAOS/3/ATTILA[3596] |
| 2005 | VPM-1/3*BEULAH//SILVERSTAR[3810] |
| 2005 | KALANNIE/81-Y-970 |
| 2006 | VN-150/VN-715[3799] |
| 2006 | BOWIE//BERSEE/3*BINDAWARRA-126937/3/BOWIE[3755] |
| 2006 | VPM/4*SUNBROOK[3791] |
| 2006 | BINDAWARRA/BOWIE//3-AG-3/WYUNA[3750] |
| 2006 | NESSER/2*VI-252[3751][3792] |
| 2007 | (DH)RAC-875//EXCALIBUR/KUKRI/3/RAC-875//EXCALIBUR/KUKRI[3794] |
| 2007 | VL-709/KRICHAUFF[3795] |
| 2007 | VI-184/SILVERSTAR[3795] |
| 2007 | LEVELHEAD//BLOCKHEAD/KINGS-JUBILEE[39][1138] |

| | |
|---|---|
| 2008 | VPM-WESTONIA/WESTONIA[3800] |
| 2008 | CGS-94-Y-00005-S/SUPER-KAUZ[3798] |
| 2008 | (DH)SUNBROOK/H-45[3596][3795] |
| 2008 | (DH)ANNUELLO/2*STYLET[3798] |
| 2008 | JANZ/SUN-129-A[3796] |
| 2008 | CNT-3/4*3765//2*CUNNINGHAM/3/2*SUNVALE[3875] |
| 2009 | KLASIC/84-Y-1426(96-Y-119)/(86-Y-204-13-23)PFAU/REEVES[3882] |
| 2009 | JANZ/CIMMYT-LINE[3875] |
| | - |
| | - |
| | - |
| | CHA/MENGAVI-8156//CIANO-67(SIB)/GALLO//BEZOSTAYA-2(CO-2224)/3/RAC-309-S[133 |
| | RAC-655//SR21/4*LANCE/3/4*BAYONET |
| | JACINTH(Colorado)/(RUS)LADOGA[37][38][39][1138][1446]; LADOGA/(AUS)JONATHAN[7 |
| | BOBIN(S)/(TR.DR)GAZA//(S)BOBIN[56][39][1173]; GULAR/(TR.DR)GAZA//GULAR[851]; BO |
| | - |
| | SPICA/KODA//GABO/3/(SIB)MENGAVI[39][117][1138] |
| | AUS-1408/3*JANZ//CUNNINGHAM[3936] |
| | WESTONIA/2*PERENJORI-26[4094] |
| | NI-601-95/(SUN-64-G)SUNDOR[1350] |
| | PENJAMO-62/4*GABO-56//TEZANOS-PINTOS-PRECOZ/NAINARI-60[279][1318] |
| | PROBSTDORFER-GIGANT/HP-35719//PROBSTDORFER-POKAL[2535] |
| | (S)TRESOR-18[39][1446] |
| | FRONTANA/KENYA-58[145][1765] |
| | VERANOPOLIS*2/EGYPTIAN-101[39][145][1765] |
| | PRELUDIO/L-10//ESTANZUELA-TARARIRAS[2965] |
| | - |
| | N-234,BGR/S-13,ITA//BEZOSTAYA-1[851][1041][1294] |
| | BLUEBIRD/INIA-66//2*TEZANOS-PINTOS-PRECOZ/3/3*ANDES-ENANO[299][606] |
| 1955 | MENTANA/FLORENCE[39][1322]; MENTANA/FLORENCE-AURORE[145][1765] |
| 1970 | CIANO-67(SIB)/3/SONORA-64/KLEIN-RENDIDOR//SIETE-CERROS-66[39]; CIANO-67//SONO |
| | II-21563/ANHINGA//FLAMINGO,MEX[145][115][1765]; JORI-C-69(SIB)/(SIB)ANHINGA//(S |
| 1927 | SUN-129-A/SUNVALE[3875]; |
| | - |
| | - |
| | ZLATNA-DOLINA/GALLINI[133] |
| 1899 | LV-CECHY[761]; (S)LV[851] |
| 1985 | NS-984-1//MIRONOVSKAYA-808/MOISSON[201][851][1592][1865] |
| | HODONINSKA-HOLICE/HEINES-KOGA/3/CARSTENS-V/HODONINSKA-OSINATKA//HEINES-K |
| | AVRORA/SO-985//PD-5517[40][1592][1865] |
| | SO-985/KAVKAZ[69][1592] |
| | - |
| | - |
| | - |
| | - |
| 1946 | VILMORIN-27/HYBRIDE-DU-JONCQUOIS[37][39][145][1790] |
| 1948 | SAISSETTE-54/SAISSETTE-D-ARLES[39][1790] |
| 1954 | PROVIDENCE/SZEKACS//K-3/ORO[39][1790] |
| 1961 | VILMORIN-27/HYBRIDE-DE-JONCQUOIS |
| 1994 | KAVKAZ/MARIS-DOVE//FIDEL[1790] |
| | COTE-D-OR/MARIVAL[39][1790][2331]; OSCAR-BENOIST/BON-FERMIER[2331] |
| | - |

K-3/HATIF-INVERSABLE/3/K-3/HATIF-INVERSABLE//LIBERATOR/4/PRECOCE-DE-JUILLET/ZA
BLE-DES-DOMES/PEVELE
-
MAGDALENA,FRA/CAPPELLE-DESPREZ[39][1790]
TARASQUE/MATCH//PURSANG/ARMINDA[1790]
-
-
PROVIDENCE/?[39][1322]; (DER)PROVIDENCE[000]
CHAMPLEIN/FD-2816-348[39][418][1531][1790]
-
VILMORIN-27/ILE-DE-FRANCE//SQUAREHEAD[39]; ILE-DE-FRANCE/VILMORIN-27//BLE-A-E
-
-
-
-
-
AE.VE/TR.DN//CAPPELLE-DESPREZ/61[1790]
-
KRANICH/CARIBO[667][851][1895]
RECITAL/FIDEL//CORDIAL[2770]
AXIAL/NRPB-84-4233[2530][4208]
-
-
BELLEVUE/CAPPELLE-DESPREZ
-
-
-
ALLIES/HATIF-INVERSABLE-BARBU//RIETI[39][1790]; RIETI/PARSEL[39][1446]
MIRONOVSKAYA-808/MARIS-HUNTSMAN//VPM-1/MOISSON/3/COURTOT[1790]; MIRONO
-
LV-DE-DORDOGNE[1790]
-
-
-
TRIBUTE,FRA/VS-73644-9-4-1
-
-
-
-
AE.VE/TR.TG//MOISSON/COURTOT[2535]
-
1979 MARIS-FREEMAN/MARIS-DURIN
NORMAN/DISPONENT[1070]; DISPONENT/NORMAN[1257][1790]; NORMAN(SIB)/DISPON
-
1957 -
TOBARI-66/II-8156-R//BEZOSTAYA-1[113][752];
LILLA/MV-8[1790]
-
1967 PENJAMO-62(SIB)/GABO-55[39][114][144][1324]/ Historical T or S
-
-
-

| | |
|---|---|
| | - |
| | LV-IND[113] |
| | - |
| | - |
| | - |
| | - |
| | - |
| | - |
| | V-761-28-J4-B2-N28 |
| | LEE//(SEL.29-LC)NORIN-10/BREVOR[39][2681] |
| | - |
| 1916 | IBRIDO-21/AKAKOMUGI |
| 1940 | DARUMA/FULTZ//TURKEY-RED[1281] |
| 1944 | NORIN-9/SHINCHUNAGA[667][841][851]; NORIN-9/SINAKANA-2-A[39][1322] |
| 1949 | WASEKOMUGI/NORIN-3[841][667][851][2822]; HAYAKOMUGI/NORIN-3[3918] |
| 1979 | USHIOKOMUGI/D.W.SEL.71//HIYOKUKOMUGI[840][841] |
| | HOKKEI-8/HOKKAI-240[39][40][840][841][851]; KITAKEI-8/HOKKAI-240[2847] |
| | FUKUOKAKOMUGI-18/SHINCHUNAGA[39][667][841][851] |
| | SAIKAI-120/USHIOKOMUGI[841][851] |
| | - |
| | - |
| | - |
| | - |
| 1951 | KENYA-223-F-1-A-3/KENYA-294-M-7-C-8[39]; KENYA-223-F/KENYA-294-M[113]; KENYA-22 |
| 1958 | - |
| 1962 | YAKTANA-54//(SEL.26-1-C)NORIN-10/BREVOR[39][954][145][114][144][1765] |
| 1966 | LERMA-ROJO-64/SONORA-64[39][953][145][114][144][1335] |
| 1970 | YAKTANA-54//NORIN-10/BREVOR[39][2331] |
| 1979 | BUCKY/(SIB)MAYA-74/4/BLUEBIRD//HD-832.5.5/OLESEN/3/CIANO-67/PENJAMO-62[114 |
| 1981 | BLUEBIRD/3/2*INIA-66/NAPO-63//II-20350/4/F2-UNKNOWN[114][144]; BLUEBIRD/3/IN |
| 1981 | KAVKAZ/(SIB)BUHO//KALYANSONA/BLUEBIRD[114][144]; KAVKAZ/KALYANSONA/4/INIA-6 |
| 1985 | BLUEJAY(SIB)/JUPATECO-73[144] |
| 1987 | PITIC-62/FRONDOSO/3/PITIC-62/MAZOE//MEXIPAK[1281]/ Reportedly boron efficient (g |
| 1988 | PEREGRINE/PF-70354/4/KALYANSONA/BLUEBIRD//ALONDRA/3/MARINGA[1281] |
| 1993 | PFAU/SERI-82(CID-93300//BOBWHITE[2964] |
| 2002 | IRENA/BAVIACORA-M-92//PASTOR[3692]; IRENE/BABAX//PASTOR[000] |
| 2002 | BABAX*2/9/KENTANA/BAGE//FRONTANA(FN)/(U)URQUIZA-O-GRAL.URQUIZA/3/BONZA/4/ |
| 2002 | CROC-1/AE.SQ(224)//OPATA-M-85/3/PASTOR[3692] |
| | VOROBEY |
| | - |
| | - |
| | FNBW74013-52 |
| | - |
| | BAW898 |
| | TACUPETO F2001 |
| | - |
| | - |
| | NAVOJOA (ATTILA/PASTOR) |
| | - |
| | CROC_1/AE.SQUARROSA (213)//PGO/3/BAV92 |
| | - |
| | - |

-
-
PASTOR//SRMA/TUI/3/SAAR
-
CROC_1/AE.SQUARROSA (224)//OPATA/3/BJY/COC//PRL/BOW/4/BJY/COC//PRL/BOW
-
JNRB.5/PIFED
ALTAR 84/AEGILOPS SQUARROSA (TAUS)//OPATA/3/ATTILA
CACUKE
-
-
-
PARUS/3/CHEN/AE.SQ//2*OPATA
CHIR3/4/SIREN//ALTAR 84/AE.SQUARROSA (205)/3/3*BUC/5/PFAU/WEAVER
-
-
SLVS/ATTILA//WBLL1
BAVIS #1 (SHARP/3/PRL/SARA//TSI/VEE#5/5/VEE/LIRA//BOW/3/BCN/4/KAUZ)
SUNCO/2*PASTOR
WORRAKATTA/2*PASTOR
W15.92/WBLL1
CROC_1/AE.SQUARROSA (224)//OPATA/3/WBLL1
FRTL//ATTILA/3*BCN
SOKOLL/WBLL1
PASTOR//HXL7573/2*BAU/3/WBLL1
-
SERI/BAV92//WBLL1
PRL/2*PASTOR/4/CHOIX/STAR/3/HE1/3*CNO79//2*SERI
-
SOKOLL/3/PASTOR//HXL7573/2*BAU
CROC_1/AE.SQUARROSA (224)//OPATA/3/SOKOLL
FRTL/SOKOLL
PUB94.15.1.12/FRTL
MEX94.2.19/PUB94.15.1.12
PASTOR//HXL7573/2*BAU/3/MEX94.2.19//ATTILA/3*BCN
MEX94.2.19//SOKOLL/WBLL1
-
-
WBLL4//OAX93.24.35/WBLL1
PUB94.15.1.12/3/FRTL//ATTILA/3*BCN
WHEAR//2*PRL/2*PASTOR
-
ATTILA/3*BCN
GRANERO INTA
-
-
-
CIMMYT HTWYT (High temperature wheat yeald trial) line
CIMMYT HTWYT (High temperature wheat yeald trial) line
CIMMYT HTWYT (High temperature wheat yeald trial) line
CIMMYT HTWYT (High temperature wheat yeald trial) line
CIMMYT HTWYT (High temperature wheat yeald trial) line

CIMMYT HTWYT (High temperature wheat yeald trial) line
CIMMYT HTWYT (High temperature wheat yeald trial) line
CIMMYT HTWYT (High temperature wheat yeald trial) line
CIMMYT HTWYT (High temperature wheat yeald trial) line
CIMMYT HTWYT (High temperature wheat yeald trial) line
CIMMYT HTWYT (High temperature wheat yeald trial) line
-
II-12300//LERMA-ROJO-64/(II-8156)SIETE-CERROS-66/3/NORTENO-67
CROC-1/AE.SQ(224)//OPATA-M-85/3/PASTOR[3692]
-
-
-
-
-
-
-
-
-
-
PASTOR//HXL7573/2*BAU/3/CMH82.575/CMH82.801
Penjamo-62/Gabo-55
CIANO-67(SIB)/PENJAMO-62//CIANO-67(SIB)/SIETE-CERROS-66
CHIBIA/PASTOR//CHIBIA
-
PJ62/GB55
OLIMPIA2/SLM
-
KS940935.7.1.2/2*PASTOR
ALTAR-84(TR.DR)/(TR.TA)CI-18[1540]
D67.2/P66.270//AE.SQUARROSA (220)/3/PRL/SARA//TSI/VEE#5/4/METSO
SCOT/MEXI_1//AE.SQUARROSA(314)/6/TZPP*2/ANE//LR64/SN64/3/CNO67/JAR//KVZ/4/[illegible]
-
-
GAN/AE.SQUARROSA (658)
-
CPI/GEDIZ/3/GOO//JO69/CRA/4/AE.SQUARROSA (208)/5/HAHN/2*WEAVER
CROC_1/AE.SQUARROSA (210)//EXCALIBUR/3/RAC-702
CROC_1/AE.SQUARROSA (213)//PGO/3/CMH81.38/2*KAUZ
ACO89/AE.SQUARROSA (309)//RAC710
-
SRN/AE.SQUARROSA (358)
CPI/GEDIZ/3/GOO//JO69/CRA/4/AE.SQUARROSA (409)
68.111/RGB-U//WARD/3/FGO/4/RABI/5/AE.SQUARROSA (878)/6/2*M5660
OPATA//CETA/AE.SQ(895)
TRAP#1/BOW//PFAU/3/VEE/TRAP#1/5/CNDO/R143//ENTE/MEXI_2/3/AEGILOPS SQUARR[illegible]
DOY1/AE.SQUARROSA (1026)//WARIGAL 5R/3/SUNCO
CETA/AE.SQUARROSA (1027)
DVERD_2/AE.SQUARROSA (214)//2*BCN/3/2*RAC710
-
CIMMYT HTWYT (High temperature wheat yeald trial) line
CIMMYT HTWYT (High temperature wheat yeald trial) line
-

| | |
|---|---|
| | AE.SQUARROSA (1029)/DVERD_2 |
| | 68.111/RGB-U//WARD RESEL/3/STIL/4/AE.SQUARROSA (188) |
| | - |
| | YAV79//DACK/RABI/3/SNIPE/4/AE.SQUARROSA (460)/5/EXCALIBUR/6/SUNCO |
| | - |
| | STY-US/CELTA//PALS/3/SRN_5/4/AE.SQUARROSA (502) |
| | SCA/AE.SQUARROSA (518) |
| | OCI//TJB368.251/BUC/3/CROC_1/AE.SQUARROSA (205)//KAUZ/4/NL 682 |
| | GREEN/AE.SQUARROSA (458) |
| | CROC_1/AE.SQUARROSA (507) |
| | ALTAR 84/AE.SQUARROSA (221)//PASTOR/3/PASTOR |
| | SCOT/MEXI_1//AE.SQUARROSA (314) |
| | CETA/AE.SQUARROSA (327)//2*JANZ |
| 2002 | - |
| 2008 | BRIMSTONE*2/HARTOG//3424.11.04.1[3798] |
| | INDIA-9/2*SOLID-STRAW-TUSCAN(F8)//DREADNOUGHT/3/CROSS-7//FEDERATION/KENYA |
| 1979 | RAVEN/(MEX)1966-ISWRN-430[851][1348][2833] |
| | - |
| | - |
| | PITIC-62/GABO-55//C-271 |
| | LV-Ivanovo,Yaroslavl[12][519]; INTROD.FROM_POL[2189] |
| 1928 | LV[39] |
| 1955 | LV-PRT[39] |
| | - |
| | - |
| | - |
| | - |
| | - |
| | - |
| 1990 | INIA-66*3/AG.DI[3599] |
| | - |
| | LV-Soria |
| | - |
| | - |
| | - |
| | - |
| | LV-FRA,CHE[39]; (S)BLANC-DE-SAVOIE |
| | (S)LV[39] |
| | - |
| | - |
| | PITIC-62/II-53-526//SONORA-64[2535] |
| 1993 | TAW-23833-73/3/GAINES/6*MIRONOVSKAYA-808//ALCEDO[1056][1369][1724] |
| | - |
| | ESTANZUELA-TARARIRAS/3/TOBARI-66//KLEIN-PETISO/RAFAELA[145][1765] |
| 1973 | SCOUT/5/KENYA-58/NEWTHATCH/3/CHEYENNE/TENMARQ//MEDITERRANEAN/HOPE/4/P |
| 1994 | PIKE/MO-9965[1519]; PIKE/MO-9965/3/STODDARD/BLUEBOY//STODDARD/D-1707[317 |
| 2000 | ND-674/ND-2710//ND-688[3227][3328] |
| | - |
| | (S)HAYNES-BLUESTEM[39][1110]; (S)BLUESTEM[5] |
| | NORIN-10/BREVOR[1281] |
| | ODESSA/TURKEY[39][1110][1451] |
| | BACKA/BEZOSTAYA-1//MIRONOVSKAYA-808/3/NS-433/4/SKOROSPELKA-35 |

| | |
|---|---|
| 1963 | F-430/S-595[37][38] |
| 1971 | F-1518(MEX)/TOKWE[39][416][1322]; SONORA-64-A//TEZANOS-PINTOS-PRECOZ/NAINAR |
| | - |
| | - |
| | (S)CESKA-CERVENKA[1592] |
| | - |
| 1964 | - |
| | - |
| | - |
| | - |
| | - |
| | - |
| | - |
| | - |
| | - |
| | - |
| | - |
| | - |
| | - |
| | - |
| | - |
| | - |
| | - |
| | - |
| | - |
| | - |
| | - |
| | - |
| | - |
| | - |
| | - |
| | - |
| | - |
| | - |
| | - |
| | - |
| | - |
| | - |
| | - |
| | - |
| | - |
| | - |
| 1987 | LV-Alger[2965] |
| 1956 | KANRED/38-MA//LIN-CALEL-MA/3/BUCK-QUEQUEN[39][145] |
| | - |
| | - |
| 1876 | (S)INTRODUCTION-FROM-FRA[851] |
| 1890 | (S)PURPLE-STRAW[39][1138] |
| 1890 | (S)DU-TOIT[37][39] |
| 1900 | YANDILLA/PURPLE-STRAW[113]; PURPLE-STRAW/YANDILLA[56][39][37][1110][1446][14 |
| 1906 | WHITE-NAPLES//FIFE/WHITE-NAPLES/3/FIFE/EDEN,AUS[56][76][39][1138]; WHITE-NAPLE |
| 1906 | (S)CARMICHAELS-ECLIPSE[39][1138][1451] |

| | |
|---|---|
| 1912 | NORTHERN-CHAMPION/CRETAN//LITTLE-CLUB[56][37][39][1138][1451]; |
| 1920 | (S)KINGS-EARLY[56][76][39][1138][1451] |
| 1921 | HUDSONS-EARLY-PURPLE-STRAW/GLUYAS[56][39][1138] |
| 1924 | GALLIPOLI/3/CURRAWA//INDIAN-4-E/FEDERATION[39][37][1138]; INDIAN-H/FEDERATION |
| 1924 | INDIAN-F/FEDERATION[56][39][1138][1451] |
| 1926 | GALLIPOLI/YANDILLA-KING[39]; (S)GALLIPOLI[56][113][1451]; CLUB-WHEAT/YANDILLA-K |
| 1927 | HARD-FEDERATION/CLEVELAND//SANDS[56][39][1138][1451] |
| 1929 | (S)BENCUBBIN[56][39][1138] |
| 1929 | NABAWA/GLUYAS-EARLY[56][39][76][1138]; GLUYAS-EARLY/NABAWA |
| 1933 | (S)GLUYAS[1451]; (S)WARDS-PROLIFIC[39][56][1138] |
| 1941 | GHURKA/3/CURRAWA//MAJOR/GALLIPOLI[56][37][39][1138] |
| 1946 | (S)PINDAR[56][39][1138] |
| 1948 | NABAWA//RIVERINA/HOPE[56][39][40][1138] |
| 1959 | R.D.R./4*INSIGNIA-49[38][56][1138][1335] |
| 1960 | BENCUBBIN//DUNDEE/GULAR/3/GULAR[56][39]; GULAR//DUNDEE/GULAR/3/BENCUBBIN |
| 1969 | THATCHER/SANTA-CATALINA//MAYO-54/3/GABO*3/CHARTER[39][117][1308][1345]; CHA |
| 1969 | VERNSTEIN/CI-12632[117][591][215] |
| 1969 | SCIMITAR/KENYA-C-6042//BOBIN/3/INSIGNIA-49[39][626] |
| 1973 | HERON/2*WW-15[39][113][2854] |
| 1973 | NORIN-10/BREVOR(SEL.14)//4*EUREKA-2/3/T-A/3*FALCON/4/THATCHER/AG.EL(T-A)//4*F |
| 1977 | PEANUT-OIL-OLYMPIC-MUTANT-132-A/SOUTH-AFRICAN-184-P-2-AIF//OLYMPIC[760][113 |
| 1978 | GLUCLUB/3/CHILE-1-B//INSIGNIA/FALCON[626][760][851] |
| 1978 | WW-15/RAVEN[760][851][1322]; South Australia |
| 1979 | WW-119/WW-15//EGRET,AUS[113][760][851] |
| 1979 | SONORA-64/YAQUI-50-ENANO//GABOTO/II-8156[626][144][760] |
| 1982 | GAMENYA//GABO*3/KHAPSTEIN(M-146)/3/(66-W-02)FALCON*3/CHILE-1-B[113][176][76 |
| 1982 | NORIN-10/BREVOR(SEL.14)//(SIB)KITE/3/KITE[690][676][1484] |
| 1983 | RAC-111/INSIGNIA[626][760][976][113][176] |
| 1983 | RAC-111/INSIGNIA[626][760][976][113][176] |
| 1983 | CONDOR,AUS/4/2*WW-15/3/STEINWEDEL/YAROSLAV-EMMER//LA-PREVISION[113][176] |
| 1984 | DARF*4/3-AG-14[760][976][113] |
| 1985 | 3-AG-14/4*CONDOR//5*OXLEY[851][1345][113] |
| 1985 | CONDOR/PITIC-62//(SIB)CONDOR[760] |
| 1986 | (F1)CONDOR//CIANO-67/2*OLYMPIC(A-LINE)(MS)/4/(R-LINE)KITE/3/DIRK-R-LINE//DEKALB |
| 1988 | PITIC-62/FESTIGUAY//2*WARIGAL[708][760] |
| 1993 | VPM-1/5*COOK//4*SPEAR[113][2755] |
| 1994 | VEERY(SIB)/QUARRION[1340][1349][2854] |
| 1996 | - |
| 1997 | MOLINEUX/3*DAGGER[1866][2854] |
| 1998 | HORNET/B-65[2854][3768] |
| 1998 | KULIN/2*BLADE[2854]; BLADE/2*KULIN[3768] |
| 1999 | MADDEN/6*RAC-177//GRAJO/76-ECN-44[2964]; SR13*3//76-ECN-44/76-ECN-36[000]; C |
| 2002 | Sr9e/3*WARIGAL//3*AROONA[3752]; 83-Z-1048/82-Z-1097[3752][3781]; 3-AG-3/4*CON |
| 2004 | 86-Z-522-62/83-Z-1175[3793] |
| 2004 | 79-W-793/83-W-1087[3781] |
| 2007 | (DH)RAC-875/KRICHAUFF//EXCALIBUR/KUKRI/3/RAC-875/KRICHAUFF/4/RAC-875//EXCA |
| 2007 | CARNAMAH/TAMMIN-18[3800] |
| 2007 | VN-150/VN-715[3799] |
| 2008 | RAC-875/YITPI[3792] |
| 2008 | (DH)RAC-875/KRICHAUFF//EXCALIBUR/KUKRI/3/RAC-875/KRICHAUFF/4/RAC-875//EXCA |
| 2008 | WYALKATCHEM/STYLET//WYALKATCHEM |
| 2009 | (DER)EGA-BONNIE-ROCK[3879] |

| | |
|---|---|
| | WAGGA-13/(S)MARSHALLS-3[39][1138] |
| | COWAN-SELECTION/DUNDEE[2619]; Australia |
| | - |
| | (S)IMPORTED-CULTIVAR[113] |
| | QT-3604/WW-725[1350] |
| | AUS-10894/4*OLYMPIC[760][851][113] |
| | HARTOG*2//VASCO/AUS-1408[2964]; Australia |
| 1948 | (S)BANKUTI-1201[1781]; BANKUTI/MARQUIS[85]; THEISS/MARQUIS[39][1592]; (S)BANKU |
| | - |
| | THATCHER/KRONEN,SWE |
| | THEISS/KANADISCHE-DICKKOPF |
| | STAMM-S-64/STAMM-MR-14-5//NS-1563[39]; STAMM-264/STAMM-14-5//NS-1563 |
| | - |
| | - |
| | TRESOR,FRA/STAAL[39] |
| | JUBILE/FYLGIA[39]; JUBILEGEM/FYLGIA[2946] |
| 1957 | COLONISTA/FRONTANA[37][39][145][1765] |
| | FRONTEIRA/MENTANA |
| | - |
| | - |
| | - |
| | - |
| 1946 | REGENT,CAN/CANUS[37][39][221][1032][1111][1323] |
| 1965 | AGRUS/6*THATCHER[47]; THATCHER*5/AGRUS(M)//THATCHER[117] |
| 1974 | CANTHATCH//(4351-331)CT-609/RESCUE[39][489][480][748] |
| 1981 | NEEPAWA*6/RL-2938/3/NEEPAWA*6//CI-81542/2*FROCOR[590][1323] |
| 1986 | RL-4302/RL-4356//RL-4359/RL-4353[1126][851]; BW-15/BW-38//BW-40/RL-4353[393 |
| 1993 | BW-83/ND-585[3894][3936] |
| | NEEPAWA/COLUMBUS(CID-188385)//BW-90[1323][1577][1792] |
| | UM-530/(MEX)CB-100[39][1323][2331]; |
| | PENTAD/MARQUIS[39][40][74]; H-44-24/MARQUIS[1323] |
| | (S)MARQUIS[10][39][74][1110] |
| | - |
| 1937 | LV-Jiagsu |
| 1965 | 69-1776/663[3499] |
| 1975 | KEZHEN/KEHONG[1334][3492] |
| | - |
| | NONGDA-183/VIRGILIO//NONGDA-183-916[2535] |
| | - |
| | - |
| | (S)NONGDA-45[1334][3492] |
| | XIANNONG-39/58-18-2//FENGCHAN-3 |
| | (S)XIAOYAN-6[3939]; ST-2422-464/XIAOYAN-86[000] |
| | - |
| | - |
| | - |
| | - |
| | - |
| 1956 | KENTANA-48/FRONTANA//MAYO-48 |
| | SIB/MARTIN//M/2*RED-WAVE[2535] |
| | - |
| 1936 | RUSKEA/MARQUIS[39] |

| | |
|---|---|
| 1975 | ERTUS/VAKKA[221][1252][1786] |
| 1975 | TAMMI/TA-c-4431[39][69][221][667][1335][1786][1930] |
| | - |
| | - |
| | KARN-I/JO-0172[38][39][1786]; KARN-II/JO-0172[221] |
| 1974 | MEXIQUE-50/B-21-VERSAILLES[418][39] |
| 1986 | MARIS-HUNTSMAN/R-267[1790]; MARIS-HUNTSMAN/(LINE_FROM_BENOIST)H-76267[13 |
| 1994 | HOBBIT/1073.32//ADAM[1347][1790] |
| | MARIS-HUNTSMAN/CASSIUS//CASSIUS[1346][667]; MARIS-HUNTSMAN/2*CASSIUS[1790] |
| | - |
| | - |
| | Benoist 40*Professeur Delos |
| | MARIS-HUNTSMAN/3/022453/R-3669//POPPY[1790]; MARIS-HUNTSMAN/022-453/R-36 |
| | - |
| | - |
| | - |
| | LV-Sahara[39] |
| | LV-FRA[1790] |
| | - |
| | - |
| | - |
| | - |
| | - |
| | - |
| | LV-FRA |
| | (S)MC-XXII |
| | LV-NORFOLK[39]; LV-FRA |
| | - |
| | - |
| | - |
| | EPI-CARRE/KRELOF//HATIF-INVERSABLE[2770] |
| | LV-HAUTE-SAVOY[39][1790] |
| | LV-FRA[1790] |
| | (S)ROUGE-D-ALSACE[1790] |
| | (S)FLORENCE-AURORE[1790] |
| | LV-FRA[1790] |
| | - |
| | LV-FRA[1790] |
| | ARMINDA/MOULIN[1413][1790]; ARMINDA/TJB-363[1346] |
| | GA-600/LOGOR//SOISSONS[2770] |
| | - |
| | - |
| | - |
| | RIETI/GROS-BLEU[39][1446][1790] |
| | - |
| | - |
| | - |
| | LV-GEO[37][9][11][39][1448] |
| 1895 | (S)ENGLISH-SQUAREHEAD[6][946][1322]; (S)SQUAREHEAD[233][1592] |
| 1959 | TRIESDORFER-21-40/RUMKERS-ERLI[1102][40][851][667][1643] |
| | LICHTIS-FRUH/MERLIN,DEU//OPAL/3/FIRLBECK-I/HALLE-MH-6706-47[39]; LICHTIS-FRUH, |
| | - |

| | |
|---|---|
| | IBIS/CARIBO//MARIS-HUNTSMAN |
| 1972 | MINISTER,BEL/HYBRID-46,GBR |
| | MARIS-RANGER/MAYO-64//CHILI-8923/MARIS-RANGER/3/MARIS-BEACON[40][667] |
| | - |
| | - |
| | - |
| | - |
| 1925 | NP-52/FEDERATION |
| 1965 | REGENT/CZECH REPUBLICCHOSLOVAKIA-3//2*C-591/3/C-217/NIPHAD-4//C-281[1324]; F |
| 1995 | NORD-DESPREZ/VG-1944//KALYANSONA//BLUEBIRD/3/YACO(SIB)/4/VEERY-5[2848][3557 |
| | - |
| | - |
| | - |
| | PASHMAK |
| | - |
| | - |
| | LV-IND |
| | LV-Punjab |
| | LV-IND |
| | - |
| | - |
| | - |
| | - |
| | - |
| | - |
| | - |
| | - |
| | - |
| | - |
| | - |
| | - |
| | - |
| | - |
| | - |
| 1947 | SAITAMA-27//INALLETTABILE-95/ARDITO[39][2494] |
| 1973 | CAMPODORE//FUNO/SAN-PASTORE[39][851] |
| 1974 | MARZOTTO/COMBINE[1067][1624][1619][1656][1764][2676] |
| 1987 | Z-1,ITA/RUBIN[1066][1558][1657]; Z-1,ITA/RUBIN,BGR[000] |
| | RIETI/WILHELMINA//AKAKOMUGI[145][39][1446] |
| | RIETI/PRINCE-ALBERT |
| 1868 | LV-JPN[1790] |
| | (S)DARUMA[851][113]; LV-KANAGAWA |
| | - |
| | - |
| | LV-IBARAGI[39] |
| | - |
| | - |
| | - |
| 1960 | AFRICA/MAYO-48 |
| 1936 | SUWON-90/SHIROBORO[879][851]; SHIROBORO/SUWON-90[1054]; SUWON-90/TOCHIG |
| | - |
| 1964 | YAKTANA-54//NORIN-10/BREVOR/3/2*YAQUI-54 |

1970 INIA-66(SIB)/NAPO-63[254][39][114][144]
1979 TOBARI-66(SIB)/NAPO-63//NOROESTE-66/ERA/3/BLUEBIRD/GALLO[144]; TOBARI-66(SIB)/
-
-
MEX94.27.1.20
HGO94.9.1.3
HGO94.9.1.5
HGO94.9.1.52
-
BJY/COC//PRL/BOW/3/URES/JUN//KAUZ
SITELLA
(S)LV[541]
-
1979 Intervarietal substitution line
LV-Transvaal[2678][3916]; ZAF
-
-
-
-
-
-
68.111/RGB-U//WARD/3/FGO/4/RABI/5/AE.SQUARROSA (629)/6/2*FRAME
-
-
-
-
-
-
-
-
-
-
-
-
-
-
-
-
-
-
-
-
-
1991 EMOU/MELCHIOR[1413][2536][2650]; EMU/MELCHIOR[1790]
-
MINISTER/PEKO//HEINES-13037[39][40]
-
-
1973 BLUEBIRD/NORTENO-67[613][114][144][39]
LV-PAK
-
-

| | |
|---|---|
| | - |
| | - |
| | - |
| | - |
| | - |
| | - |
| | (S)THORI[541] |
| | LV-PALESTINE[000] |
| | - |
| | CARSTENS-V/HODONINSKA-OSINATKA//KOGA[39] |
| | OSTKA-GRODKOWICKA/SQUAREHEAD-GRODKOWICKA |
| | MALGORZATKA-UDYCKA/PETIT-QUINQUIN[39][946] |
| | - |
| | - |
| | KOOPERATORKA/BANKUTI-1205[39][2834] |
| 1928 | LV-Dagestan[000] |
| 1985 | MONAKINKA/ACADIA[28][1156][1369] |
| | BELOTURKA/POLTAVKA[143][11] |
| | - |
| | LV-Dagestan[000] |
| | (M)PPG-870[1353][2836][2913] |
| 1958 | UNIE-52-A/KRUGER[39] |
| | - |
| | - |
| | NEBRED/ARAGON-0-3 |
| | H-1-1(TR.DR)/(AE.VE)AP-1//(TR.AE)ALMATENSE-H-10-15[788] |
| | - |
| | EXTRA-SQUAREHEAD-II/TYSTOFTE-SMAAH |
| | - |
| | - |
| | - |
| | - |
| | - |
| | - |
| | - |
| | - |
| | - |
| 1972 | TEZANOS-PINTOS-PRECOZ/SONORA-64//LERMA-ROJO-64/SONORA-64/3/SONORA-64-A//S |
| | PITIC-62/II-53-526//SONORA-64 |
| | SONORA-64/KLEIN-RENDIDOR |
| | - |
| | - |
| | - |
| | - |
| | - |
| | - |
| | - |
| | - |
| | (S)BANATKA,UKR[1448]; (S)BANATKA-KHERSONSKAYA[6][2706] |
| 1952 | MACVICAR/PB-1-87-WW-149[2965] |
| 1963 | FRONTANA/KENYA-58//NEWTHATCH[39] |
| 1984 | TAM-105*4/AMIGO[293][607][982][1078] |

-
LV-EASTERN-USA[1110]
LV-CHL[1318]
FULHIO/PURKOF/3/TRUMBULL//HOPE/HUSSAR,USA/5/FAIRFIELD/4/PI-94587/3/HUNGA
BENHUR(SIB)/FORLANI[186][483][683][851][2331]
NORIN-10/BREVOR(SEL.14)//ANDERSON/COKER-55-9
BLUEBOY//THORNE*5/199-4/3/VA-68-24-42[330][685][186][1318]
KENYA-58/NEWTHATCH//HOPE/2*TURKEY/3/CHEYENNE/4/PARKER[273][161][39][867][2
-

*2/KENYA-117-A[626]; GABO/4/GABO*5/MENTANA//2*GABO/3/KENYA-117-A[1138]

GABO*3/KHAPSTEIN[000]

760]

PIC[760]

ON(19-FM-74)//3*CONDOR/TA-3-PNB-3-P/WW-33-G/CONDOR*2/WW-33-B[1349]

'55][760]

I-95/QUALSET-601-20[2770]

)][3762]

[[1866]

CIANO-67/SIETE-CERROS-66//KALYANSONA/BLUEBIRD[2854]/ mapping population parent

[2854][3766]; QT-2327/COOK//QT-2804[3596]

VARIQUAM//KLOKA/PITIC-62/3/HALBERD/4/3-AG-3/AROONA[1832] - mapping population parent
V-15/QT-7605[3770]

M-1[2854]; TINCURRIN*4/3/LANCE*2/CONDOR*4/3-AG-14-14/TATIARA*3//COOK*5/VPM-1[3596][3761]

AU[3793]

5]

4]
3IN*2/GAZA[113]; BOBIN-W-39//BOBIN-W-39/GAZA[1281];

DRA-64/KLEIN-RENDIDOR/3/II-8156[145][114][144][1765]
IB)FLAMINGO,MEX[851]

OGA[1592]; HODONINSKA-HOLICE/KOGA//(S)DIANA-I[39][2370]

NDA[1790]; K-3/HATIF-INVERSABLE//K-3/HATIF-INVERSABLE[39]; K-3/HATIF-INVERSABLE//K-3/HATIF-INVERS

PI-CARRE-TYPE-HEINES-VII[40][1052][1790]

OVSKAYA-808/MARIS-HUNTSMAN/3/VPM-1/MOISSON//COURTOT[1790]; MIRONOVSKAYA-808/MARIS-HUNT

ENT[1346][1413][1478]

:3/KENYA-294[517]; KENYA-223*4/KENYA-SETTLER[2331]

][144]
IA-66(SIB)/NAPO-63*2//II-20350[39]
6//LERMA-ROJO-64/BLUEBIRD/3/2*BLUEBIRD[USLOVNO]

ood fertility under bow B conditions)

/TORIM-73/5/ALDAN/6/SERI-82/7/YECORA-70/8/OPATA-85[3589]

MN72252/5/SHI4414/CROW/7/METSO

OSA (TAUS)/4/WEAVER

-744

AWNEE/CHEYENNE[39][803][1322]
1]

I-60/3/TOKWE[2331]

51]; YANDILLA-KING/PURPLE-STRAW[74]; PURPLESTRAW-14-A/YANDILLA[1322];
S*2/IMPROVED-FIFE//IMPROVED-FIFE/EDEN,AUS[145][1111][1765][1451]

N//CURRAWA[56][1451]; ZAFF/YANDILLA-KING[39];

ING[1322]

N[76][1138]; DUNDEE/GULAR//BENCUBBIN(F4)/3/GULAR[113];
ARTER/3*GABO/3/SANTA-CATALINA/THATCHER//MAYO-54[2331]

FALCON/5/THATCHER/AG.EL(T-A)//5*FALCON[39][113][1322]
]

50][851][1335]

][760][851][976]

3-436-R-LINE/PRIMEPI[591][113]

CO-1213/RAC-549[3758]; DRP((FRONTANA/KENYA-58//N10B/G55)NAINARI-60)/(TOBARI-66/CIANO-67//TOB
NDOR//3*MILLEWA/3/BODALLIN[3752];

LIBUR/KUKRI[3794]

LIBUR/KUKRI[3794]

TI-5[000]

2]

47][1642]

569/POGGY[1642]; MARIS-HUNTSMAN/3/022453/R-3669//CORIN[000]

//OPAL/SOLO[1252]; LICHTIS-FRUH/HEINE-STAMM/OPAL/SOLO[1531];

REGENT-1974/3*CZECH REPUBLICCHOSLOVAKIA//2*C-591/3/819/C-281[39][1322]; REGENT-1974/3*CZECH
7]

ISHIROBORO[2831]

/NAPO-63//NOROESTE-66/ERA,USA/3/BLUEBIRD/GALLO,MEX[000];

SELKIRK*3/ANDES[39]

RIAN//FULTZ/HUNGARIAN/8/LAPORTE/7/MINHARDI/WABASH/5/FULTZ/HUNGARIAN//IL-1/3/WABASH/4/F

331]

SABLE/3/LIBERATOR//PRECOCE-DE-JUILLET/ZANDA[1052]

TSMAN/3/VPM-1/MOISSON//9*COURTOT[2839]

[illegible]ARI-66/8156/3/CALIDAD//BLUEBIRD/CIANO-67)/2/MADDEN/6*RAC-177[3758];

H REPUBLICCHOSLOVAKIA//C-591/3/P-1-19/C-281[543]; REGENT-1974/CH-23//2*C-591/3/P-19/C-281[793

AIRFIELD/6/REDCOAT//WISC-245/AFGHANISTAN/9/KNOX*4/PD-5127/10/KNOX*2/FREX//(SIB)RILEY[283]

3];
