## Supplementary Table 3 for "The wheat *Seven in Absentia* gene is associated with increases in biomass and yield in hot climates" (from another version of this preprint: version 2, 2019-08-13)

| method | Marker/gene model | Allele 1 primer sequences |
|---|---|---|
| KASP | AWG61 | GAAGGTGACCAAGTTCATGCTACCCCGAGACAGGCGAGG |
| KASP | AWG13 | GAAGGTGACCAAGTTCATGCTACGCTATCAACTTCCCGATCCAA |
| KASP | AWG66 | GAAGGTGACCAAGTTCATGCTGGCAGGACCCAGAGACACTG |
| KASP | AWG10 | GAAGGTGACCAAGTTCATGCTGTGGAGCAGCCTACGGCCAA |
| KASP | AWG31_1 | GAAGGTGACCAAGTTCATGCTGGCTTGGCTCTTTCCAGATTGATT |
| KASP | AWG41 | GAAGGTGACCAAGTTCATGCTCGCCGGCGAGAGGT |
| KASP | AWG84 | GAAGGTGACCAAGTTCATGCTGTTTGCCACTAGTTCTGACTGGAT |
| KASP | AWG27 | GAAGGTGACCAAGTTCATGCTTCGTGTTTGCTGCGGTCTGCT |
| KASP | AWG57 | GAAGGTGACCAAGTTCATGCTGTTTGGACGAAAGCCCTATTTTCTG |
| KASP | AWG47_2 | GAAGGTGACCAAGTTCATGCTAATAATGGTGTCCATGCACCGTTTC |
| KASP | AWG62 | GAAGGTGACCAAGTTCATGCTCCTCGGCGGCGGCTGTAG |
| KASP | AWG21 | GAAGGTGACCAAGTTCATGCTCCAAACAGCCACTAAGTGAGGAG |
| KASP | AWG43_1 | GAAGGTGACCAAGTTCATGCTGATGTTGACACTTTGTGTTGTTGCC |
| KASP | AWG69_1 | GAAGGTGACCAAGTTCATGCTCGCTAGGTTTCTGCGACCGC |
| KASP | AWG75 | GAAGGTGACCAAGTTCATGCTGGTTCAAGATCAGGTTCATGATCAG |
| KASP | AWG19_1 | GAAGGTGACCAAGTTCATGCTCCTGGTGTCCATCCATCGTCTT |
| KASP | AWG67_1 | GAAGGTGACCAAGTTCATGCTGGGGAGTGGCCTCTGCCTT |
| KASP | AWG38 | GAAGGTGACCAAGTTCATGCTACTGTGAATTGGCCATTCTTCTCTG |
| KASP | AWG82 | GAAGGTGACCAAGTTCATGCTCGGCAGCAGATTGAGCGTCG |
| KASP | AWG25 | GAAGGTGACCAAGTTCATGCTGTTGTTGAGGATGCGGGTT |
| KASP | ADW313 | GAAGGTGACCAAGTTCATGCTCCATGTTAAGATATAGTCATCTAACTGC |
| KASP | ADW314 | GAAGGTGACCAAGTTCATGCTTTTTCTTTTGATGGCTACTATGGTGC |
| KASP | ADW316 | GAAGGTGACCAAGTTCATGCTTGGTCTGTGCCTCTGTGATCA |
| KASP | ADW317 | GAAGGTGACCAAGTTCATGCTCGATGCTCACGCCAGACAACTAA |
| KASP | ADW318 | GAAGGTGACCAAGTTCATGCTCCCCTCGTGCTACATCTCGA |
| KASP | ADW319 | GAAGGTGACCAAGTTCATGCTAATCTATGATCACCGGTGCCCA |
| KASP | ADW320 | GAAGGTGACCAAGTTCATGCTAATAATTAAATGGAAGAGGCCTATGAGATA |
| KASP | ADW322 | GAAGGTGACCAAGTTCATGCTGAAGCAACATCATGAATTCATGCGG |
| KASP | ADW323 | GAAGGTGACCAAGTTCATGCTAATCCCAGTGGCATCATTTACAGC |
| KASP | ADW324 | GAAGGTGACCAAGTTCATGCTCCGCCAAACGACCCAG |
| KASP | ADW328 | GAAGGTGACCAAGTTCATGCTCAGCCATGCTGCTTGTAGCGG |
| KASP | ADW334 | GAAGGTGACCAAGTTCATGCTGTTGTACCCGGTCTAGGATCGT |
| KASP | ADW330 | GAAGGTGACCAAGTTCATGCTCGGTTAGGTATTGTGAGGCAACG |
| KASP | ADW332 | GAAGGTGACCAAGTTCATGCTGGTTTGGGCCTGTCCAACCG |
| KASP | ADW315 | GAAGGTGACCAAGTTCATGCTCAATATCTTGAATTCCAAGGCCCCA |
| KASP | ADW337 | GAAGGTGACCAAGTTCATGCTCGCATATCCTCCAATTTCGCTTTCT |

| | | |
|---|---|---|
| KASP | ADW338 | GAAGGTGACCAAGTTCATGCTGTCTCCATGCATGAGAAATCCAG |
| KASP | ADW341 | GAAGGTGACCAAGTTCATGCTCAAGATGTCGACACCCACCCTT |
| KASP | ADW342 | GAAGGTGACCAAGTTCATGCTCATCATCGAAACAGGGGCTAGGA |
| KASP | ADW343 | GAAGGTGACCAAGTTCATGCTGGGGACCCCATAAAGTTCATCTG |
| KASP | ADW345 | GAAGGTGACCAAGTTCATGCTGGCCGGCGCTTGGCTCG |
| KASP | ADW347 | GAAGGTGACCAAGTTCATGCTGTGTATGCGATCGGTCCGCTG |
| KASP | ADW348 | GAAGGTGACCAAGTTCATGCTGTCGGCCAAGCTAGATCGCTC |
| KASP | ADW349 | GAAGGTGACCAAGTTCATGCTAGCGAAGTGGGCCGTCCAG |
| KASP | ADW351 | GAAGGTGACCAAGTTCATGCTCATTTAATTTTCTGCATGTATCAAATTTGTAGA |
| KASP | ADW353 | GAAGGTGACCAAGTTCATGCTCCACATGCCACTTCCTCTTTTTTTTTA |
| KASP | ADW363 | GAAGGTGACCAAGTTCATGCTCGAAGTCACCATAGTCCACAGAC |
| KASP | ADW364 | GAAGGTGACCAAGTTCATGCTCCTCATCCCACATCTGCTGAG |
| KASP | ADW371 | GAAGGTGACCAAGTTCATGCTAACTTATTGGGAAGGAGCGCCATT |
| KASP | ADW373 | GAAGGTGACCAAGTTCATGCTGAGGTGGGTTGCTTCCAAACG |
| KASP | ADW374 | GAAGGTGACCAAGTTCATGCTCGTGCGTCCGTTTCACGTAAGA |
| KASP | ADW431 | GAAGGTGACCAAGTTCATGCTCCTCTATGAACGCATGCACACAC |
| KASP | ADW432 | GAAGGTGACCAAGTTCATGCTGAGCGTTTGTGCGTGAACTTTTTTTTT |
| KASP | ADW433 | GAAGGTGACCAAGTTCATGCTATGACGTGGAACCCAATATTCACATTT |
| KASP | ADW469 | GAAGGTGACCAAGTTCATGCTACTCGACATGACGGTCCACTT |
| KASP | ADW477 | GAAGGTGACCAAGTTCATGCTCACAGTATAGCAGTTCATTCGGACAT |
| KASP | ADW576 | GAAGGTGACCAAGTTCATGCTACACAGATCCTCCATAAAACAGAATTTTTA |
| KASP | ADW577 | GAAGGTGACCAAGTTCATGCTGTAGTCTCACTGGAGCTCTCCA |
| KASP | ADW594 | GAAGGTGACCAAGTTCATGCTGATGATGTCCTTAGACGTGGGC |
| KASP | ADW595 | GAAGGTGACCAAGTTCATGCTTTGGGATAATCTACAGTATGTTTAACTCA |
| KASP | ADW616 | GAAGGTGACCAAGTTCATGCTGCCACGGGGTGTTCGTGGAAA |
| KASP | ADW617 | GAAGGTGACCAAGTTCATGCTTCAAGCACTCTAATTATCATCCTTC |
| KASP | ADW657 | GAAGGTGACCAAGTTCATGCTAAAAAATGCAGACCACAATTGTCTGTC |
| KASP | ADW659 | GAAGGTGACCAAGTTCATGCTATACCAGTGTCATCCATGCACTAG |
| PCR | Indel-1 | TCTGAACCTCTGCCATTGCT |
| PCR | Indel-2 | CGGTGAAACGTCAATTGGGC |
| PCR | Indel-3 | ACAGGAACGTCACTTCCTCTTC |
| PCR | Indel-4 | GAACCTCTGCCATTGCTTGC |
| PCR | Indel-5 | ACGTCTACCCCTACTTCGCT |
| PCR | Indel-6 | GACATAGCTGCCAGTGGGTT |
| PCR | Indel-7 | ACTTTTTCTTAGCCGATGCA |
| PCR | Indel-8 | GTCAAATGCATGAGCTTTAC |
| QPCR | TraesCS3B01G570900 | ATCCATACTTCCTCGATGATC |

| | | |
|---|---|---|
| QPCR | TraesCS3B01G571000 | ATTCTTTGGCCCGGTGGA |
| QPCR | TraesCS3B01G572500 | CTTGACATTAGCCAATCCAAA |
| QPCR | TraesCS3B01G572600 | CAATGAATATACCATGATGTTGG |
| QPCR | TraesCS3B01G572700 | GTACACTGACATCAAGGGCTT |
| QPCR | TraesCS3B01G572800 | GGATAAGGAGAAGGATGTGCA |
| QPCR | TraesCS3B01G572900 (TaSINA) | ATCGAGAAGCTCCCGGTT |
| QPCR | TraesCS3B01G573000 | TCCTTCTGACACTACTCCTCCTG |
| Sanger sequencing of TaSINA (- 4,742 to - 2,780 bp) | ADW611 | GTTTATATGGTCTAGATGGTTGAAC |
| Sanger sequencing of TaSINA (- 3,088 to - 1,911 bp) | ADW612 | AAACTCGAATTCCTCCTAT |
| Sanger sequencing of TaSINA (-2,394 to -293 bp) | ADW613 | CTAAAAGTGAGTCTCTCTGG |
| Sanger sequencing of TaSINA (- 1,052 bp to +273 bp) | ADW664 | TCAAACCAAAGACGGAACCGAG |
| Sanger sequencing of TaSINA (+ 42 to + 1,167 bp) | ADW615 | GCCGTGGTGCCCATGGAG |
| QPCR control gene | TaActin | GACAATGGAACCGGAATGGTC |
| QPCR control gene | TaCyclophilin 2*2 | CAAGCCGCTGCACTACAAGG |
| QPCR control gene | TaGAPdH 2*2 | TTCAACATCATTCCAAGGAGCA |
| QPCR control gene | TaEFA2*2 | CAGATTGGCAACGGCTACG |

| Allele 2 primer sequences | Common reverse |
|---|---|
| GAAGGTCGGAGTCAACGGATTGACCCCGAGACAGGCGAGA | ACGGCAGGAGGGCCAACTGTAA |
| GAAGGTCGGAGTCAACGGATTCGCTATCAACTTCCCGATCCAG | CGATGCTCTGTGCCTGCAGGAA |
| GAAGGTCGGAGTCAACGGATTGGCAGGACCCAGAGACACTA | GGACATGCACAGTGACCACGTTAAA |
| GAAGGTCGGAGTCAACGGATTGGAGCAGCCTACGGCCAC | TTTTGCTCCTTCAACGCGACAGCTA |
| GAAGGTCGGAGTCAACGGATTGGCTTGGCTCTTTCCAGATTGATA | GTAGCTGTGGGAGTGAGTGATGAA |
| GAAGGTCGGAGTCAACGGATTGCTCGCCGGCGAGAGGC | CGTGTGCTGGAAGAAGGCGTCAT |
| GAAGGTCGGAGTCAACGGATTTGCCACTAGTTCTGACTGGAC | GTAACCAGCCACATCAGCATGACAA |
| GAAGGTCGGAGTCAACGGATTCGTGTTTGCTGCGGTCTGCG | AAAACAAAAAGAAGAGGAAGCCCAAGCTA |
| GAAGGTCGGAGTCAACGGATTAGTTTGGACGAAAGCCCTATTTTCTA | ACGAGAGCCGCTCACCCACAAT |
| GAAGGTCGGAGTCAACGGATTCAATAATGGTGTCCATGCACCGTTTT | CTCCTGACAACCATGCCTTCAACAA |
| GAAGGTCGGAGTCAACGGATTCCTCGGCGGCGGCTGTAT | CAAACGTTGTCGCCCGGCCAT |
| GAAGGTCGGAGTCAACGGATTCCAAACAGCCACTAAGTGAGGAA | AGCGATCCCCCCTTTGACCGAA |
| GAAGGTCGGAGTCAACGGATTGATGTTGACACTTTGTGTTGTTGCG | AGCTGCTACAACAATCAAACATTGCAGAT |
| GAAGGTCGGAGTCAACGGATTCCGCTAGGTTTCTGCGACCGT | CGCAGCAGCCCAGGCGCAA |
| GAAGGTCGGAGTCAACGGATTGGTTCAAGATCAGGTTCATGATCAC | AATTGTACTTGCATGGTAAGCTCTGGAAT |
| GAAGGTCGGAGTCAACGGATTCTGGTGTCCATCCATCGTCTC | GATTTATGAGCGCGCTGAATGGGTT |
| GAAGGTCGGAGTCAACGGATTGGGAGTGGCCTCTGCCTC | CACCACCACGAGTGCTGCGAT |
| GAAGGTCGGAGTCAACGGATTAAACTGTGAATTGGCCATTCTTCTCTA | TTTTGGGGATGATTTCATTGCCTGGAAA |
| GAAGGTCGGAGTCAACGGATTCCGGCAGCAGATTGAGCGTCA | GACTCTCCGTCGCCTCTGGAT |
| GAAGGTCGGAGTCAACGGATTCTGTTGTTGAGGATGCGGGTG | TGTTGCTCCTTGAATTCACTTATGCAGAT |
| GAAGGTCGGAGTCAACGGATTCCATGTTAAGATATAGTCATCTAACTGT | TCTTCCTATCAAATCGGTCTTTTGGGATA |
| GAAGGTCGGAGTCAACGGATTTTTCTTTTGATGGCTACTATGGTGG | TGACACAGACGCCCGTGACCTA |
| GAAGGTCGGAGTCAACGGATTGGTCTGTGCCTCTGTGATCG | GGCGACGCTTGGATCTCTGGTT |
| GAAGGTCGGAGTCAACGGATTGATGCTCACGCCAGACAACTAC | GTCGAGGATCGCTCTGCCTTGAT |
| GAAGGTCGGAGTCAACGGATTCCCCTCGTGCTACATCTCGC | TCGGCATGACAGGCAGCGACAA |
| GAAGGTCGGAGTCAACGGATTAATCTATGATCACCGGTGCCCG | GCAGAGTTGGAGAAGTCGAAGCTTT |
| GAAGGTCGGAGTCAACGGATTAATTAAATGGAAGAGGCCTATGAGATG | GATGGCATCCACATTTAGCTTTTACCAT |
| GAAGGTCGGAGTCAACGGATTAAGAAGCAACATCATGAATTCATGCGT | GGCAGTCTGATAGCCCAAAGAACTT |
| GAAGGTCGGAGTCAACGGATTGAATCCCAGTGGCATCATTTACAGT | GCTTGCTGTACAAAAGCATGGTACCAT |
| GAAGGTCGGAGTCAACGGATTTGCTCCGCCAAACGACCCAT | TATGGCCTTGTCTACCCTGGAGTT |
| GAAGGTCGGAGTCAACGGATTCAGCCATGCTGCTTGTAGCGA | CTCAGTCGAGTGACGTTAGCATCTT |
| GAAGGTCGGAGTCAACGGATTGTACCCGGTCTAGGATCGC | CCTACGGTGCGTTCAAGTTGATGTA |
| GAAGGTCGGAGTCAACGGATTACGGTTAGGTATTGTGAGGCAACA | CGCCCACTGCTGCCGCCAT |
| GAAGGTCGGAGTCAACGGATTCGGTTTGGGCCTGTCCAACCA | GTCCTTATATAGGAGGTCAGAAACCAA |
| GAAGGTCGGAGTCAACGGATTAATATCTTGAATTCCAAGGCCCCC | GGTGGAGTGTATTATGCCGACCTAA |
| GAAGGTCGGAGTCAACGGATTGCATATCCTCCAATTTCGCTTTCG | CACCGAGAATCTCATTGATGAGGAAATAT |

| | |
|---|---|
| GAAGGTCGGAGTCAACGGATTACGTCTCCATGCATGAGAAATCCAA | CGGGGCGAAAGCCTTTTTCGATAAT |
| GAAGGTCGGAGTCAACGGATTCAAGATGTCGACACCCACCCTA | TCATCGGCCGGTGGCACACATA |
| GAAGGTCGGAGTCAACGGATTATCATCGAAACAGGGGCTAGGC | GGCGTCTCCCTGAAGATGGAATAAA |
| GAAGGTCGGAGTCAACGGATTGGGGACCCCATAAAGTTCATCTC | GTTCTTGCGCAGGGTGTTGACGAT |
| GAAGGTCGGAGTCAACGGATTATGGCCGGCGCTTGGCTCA | CTCGTCTTCCGAGTCCATTAAACGAA |
| GAAGGTCGGAGTCAACGGATTGTGTATGCGATCGGTCCGCTT | CTGCTCTGGGCACCTAAAGATATCTT |
| GAAGGTCGGAGTCAACGGATTGTCGGCCAAGCTAGATCGCTT | CGTAATGTCGACGTTTGCAGCGAT |
| GAAGGTCGGAGTCAACGGATTGAGCGAAGTGGGCCGTCCAA | AACCGGAAACAGAAAGCCTTGCGTA |
| GAAGGTCGGAGTCAACGGATTCATTTAATTTTCTGCATGTATCAAATTTGTAGT | GAGCCTAAATTAGTATGAGTTCTACCACAT |
| GAAGGTCGGAGTCAACGGATTCCACATGCCACTTCCTCTTTTTTTTTT | GACATGATCTACTTTGCATACACAATGTAT |
| GAAGGTCGGAGTCAACGGATTACGAAGTCACCATAGTCCACAGAT | TGTGAGGAGACATTGCCGTCGATTA |
| GAAGGTCGGAGTCAACGGATTGCCTCATCCCACATCTGCTGAT | GAAATGGGTGGATAGGGAACTCGAT |
| GAAGGTCGGAGTCAACGGATTCTTATTGGGAAGGAGCGCCATC | GGTCGTGTGTTGGAGTTGTCCTA |
| GAAGGTCGGAGTCAACGGATTGGAGGTGGGTTGCTTCCAAACA | CATCAATTGGTGGCTGCTCCAAGTA |
| GAAGGTCGGAGTCAACGGATTGTGCGTCCGTTTCACGTAAGG | GGGTGCAGCCCAGTAAACAAAAGAA |
| GAAGGTCGGAGTCAACGGATTACCTCTATGAACGCATGCACACAT | GAGAAGCTCGCAGAGACGGCAT |
| GAAGGTCGGAGTCAACGGATTAGCGTTTGTGCGTGAACTTTTTTTTC | GAGAGCTCATCCAACTGGTTTTCCAA |
| GAAGGTCGGAGTCAACGGATTGACGTGGAACCCAATATTCACATTG | GACATGTGGGCCCTACTCGACAT |
| GAAGGTCGGAGTCAACGGATTACTCGACATGACGGTCCACTC | CATTGTAAATGATGACGTGGAACCCAATA |
| GAAGGTCGGAGTCAACGGATTACAGTATAGCAGTTCATTCGGACAG | CGCCAATATTTGGCAATTAGATGAACCAT |
| GAAGGTCGGAGTCAACGGATTACACAGATCCTCCATAAAACAGAATTTTTT | TATTATTATTAGGGCACATTTTCCGAATTT |
| GAAGGTCGGAGTCAACGGATTAGTCTCACTGGAGCTCTCCG | CAAGCTTGTTGGACATCATAGGAAAGTT |
| GAAGGTCGGAGTCAACGGATTGATGATGTCCTTAGACGTGGGT | TATTATGGTAGACCCTTCTTTCATTAAGAT |
| GAAGGTCGGAGTCAACGGATTTGGGATAATCTACAGTATGTTTAACTCT | CTTTATTCGGCATGGATGCTATTCCATA |
| GAAGGTCGGAGTCAACGGATTCCACGGGGTGTTCGTGGAAC | CTCTCATGCCCTAAGCCAACCTAA |
| GAAGGTCGGAGTCAACGGATTCTTCAAGCACTCTAATTATCATCCTTG | GCCCAGATGGTTTTCCCGCATATTT |
| GAAGGTCGGAGTCAACGGATTAAAAAATGCAGACCACAATTGTCTGTG | GTAGCGGTACACCCAGTGGCAA |
| GAAGGTCGGAGTCAACGGATTCATACCAGTGTCATCCATGCACTAA | ACTGCATTTATCTGGCCGGACCAAA |
| - | AGAAGTCACGGGCTTTGAGT |
| - | TGTGTTGACGAAGCAGCTTT |
| - | ATACACGGGAGAAGGATGAGGA |
| - | TTGCTAGGCCGTTCTGGATG |
| - | TCTCGACCTCATCCCCTGGT |
| - | CCCGGCACAGCACTTAAGTA |
| - | TTGAAGTAGGCATTCTCAAC |
| - | GTGTGGATGCACGTGTATTT |
| - | CCAGGAGTAGTACAAGGTCAC |

| | |
|---|---|
| - | CGAGTTGGGAAGAGATCGTAA |
| - | GGATGCCGTCGGTGTTGC |
| - | GGTAATAGACGCAGTTGGCAT |
| - | TGACCTCTCCGGACGCGA |
| - | CCTCCTTCTCACGACGCTCA |
| - | TTGCCTTGATCTGTCCCT |
| - | TTGCCTTGATCTGTCCCT |
| - | ATGATAATATCCATCGAGACAGG |
| - | GTGATGAGCTATCAATGG |
| - | TACCATCCACCTAACCTA |
| - | CAAGATTCATGCACACCAGATAC |
| - | CGTTTCTTGTAGCTCTGATCGGTG |
| - | GTGTGATGCCAGATTTTCTCCAT |
| - | AGGGGACGGTGCAGATGAA |
| - | CGTAACCCAAAATGCCCTTG |
| - | CGGACAGCAAAACGACCAAG |
